# Chitinase-1 inhibition reverses metabolic dysregulation and restores homeostasis in MASH animal models

**DOI:** 10.1101/2024.10.27.620490

**Authors:** Katarzyna Drzewicka, Katarzyna Głuchowska, Michał Mlącki, Bartłomiej Hofman, Irina Tuszynska, Tristram Ryan, Katarzyna Piwowar, Bartosz Wilczyński, Dorota Dymkowska, Barbara Dymek, Tomasz Rejczak, Kamil Lisiecki, Adam Gołębiowski, Adam Jagielski, Angelika Muchowicz, Dylan Ryan, Krzysztof Zabłocki, Luke A.J. O’Neill, Zbigniew Zasłona

## Abstract

OATD-01 is a chitinase-1 (CHIT1) inhibitor, reducing inflammation and fibrosis in animal models where chronic inflammation leads to tissue remodeling. CHIT1, predominantly secreted by macrophages, is overexpressed in metabolic-dysfunction-associated steatohepatitis (MASH). In this study, we demonstrated the efficacy of OATD-01 in two murine and a rat model of MASH. RNA-Seq analysis revealed that OATD-01 reversed MASH-dysregulated genes. Apart from the attenuation of inflammation and fibrosis, OATD-01 regulated metabolic processes such as lipid metabolism and glycolysis. We demonstrated that both genetic and pharmacological inactivation of CHIT1 resulted in inhibition of glycolysis and glucose uptake in primary macrophages. As a consequence, we observed increased ATP, lower citrate and increased acetate levels, resulting in a reduced IL-1β secretion. These results revealed the key role for CHIT1 in regulating metabolism. OATD-01 is a macrophage modulator that can directly restore metabolic balance and consequently inhibit inflammation and fibrosis, supporting its use for MASH treatment.

## Introduction

Chitotriosidase (CHIT1) is an ancient glycosidase, targeting GlcNAc-rich glycans, which abundantly modify proteins and to some extent also lipids (Kopitz, 2017; Reily et al., 2019). Previously we have shown that CHIT1 is expressed in pathologically activated macrophages, and inhibition of chitinolytic activity is beneficial in animal models of diseases where chronic inflammation leads to fibrosis, such as severe asthma (Sklepkiewicz et al., 2023), lung fibrosis (Koralewski et al., 2020; Sklepkiewicz et al., 2022), lung sarcoidosis (Dymek et al., 2022) and inflammatory bowel disease (Mazur et al., 2021, 2022). This was mostly thanks to the development of OATD-01, which is a small molecule inhibitor of CHIT1 (OATD-01) with drug-like properties detailed in our study from Koralewski et al., 2020. Its efficacy was proven in vitro, with an IC50 of 23 nM against human CHIT1 and 28 nM against murine CHIT1, as well as in the aforementioned animal models. Completion of preclinical development led to the clinical nomination of OATD-01, which was later proven safe in phase I clinical trials and is currently in phase II trials with patients suffering from lung sarcoidosis (the KITE study; NCT06205121).

In this study we demonstrated the efficacy of CHIT1 inhibitor (OATD-01) in metabolic dysfunction-associated steatohepatitis (MASH). MASH is characterized by fat accumulation in the liver (steatosis), which leads to damage of hepatocytes, followed by inflammation and fibrosis (Parola & Pinzani, 2024), presenting a major health problem in Western societies and an urgent need for efficient drugs. Specifically, pharmacological intervention to prevent the final outcome of the disease (F3/F4) would be beneficial, since these later stages involve fibrosis and liver cirrhosis that can be treated only by liver transplantation (Burra et al., 2020; Van Gaal et al., 2021). OATD-01 as an immunomodulatory drug with proven preclinical efficacy in reversal of fibrotic changes would fit this description.

The importance of macrophages, and specifically monocyte-derived macrophages in the development of MASH is well documented. Recent data illustrated the crucial role of both resident and recruited macrophages in the development of MASH (Kazankov et al., 2019; Remmerie, Martens, & Scott, 2020; Remmerie, Martens, Thoné, et al., 2020; Sakai et al., 2019; Seidman et al., 2020; Tran et al., 2020; Wan, Benkdane, Alons, et al., 2014; Wan, Benkdane, Teixeira-Clerc, et al., 2014). CHIT1 expression is activated in Kupffer-Browicz cells (resident liver macrophages) of MASH patients, compared to MAFLD patients or healthy donors (Malaguarnera et al., 2006). Moreover, very strong genetic evidence indicates that a 24-bp duplication in exon 10 of *Chit1* gene, leading to enzyme absence and loss of function, prevents MASH development (Di Rosa, Mangano, et al., 2013), further justifying therapeutic potential of CHIT1 inhibition.

Here, for the first time we provide mechanistic insights into OATD-01 protective action driven by a metabolic switch. Key pathways modulated by OATD-01 were revealed from the RNA sequencing data obtained from liver homogenates from a MASH model. Most upregulated pathways in MASH and reversed by OATD-01 treatment were all related to metabolic processes, including acetyl-CoA metabolism, triglyceride metabolism, fatty acid catabolism, cholesterol flux/biosynthesis, and glycolysis – confirmed later in a functional cellular assay. Taken together, our data implicated suppression of macrophage glycolysis-driven activation as a protective mechanism of OATD-01 in MASH and presented CHIT1 as a regulator of cellular metabolism.

## Results

### The inhibition of CHIT1 by OATD-01 is efficacious in animal models of MASH

To investigate the efficacy of OATD-01 in reducing MASH severity, we performed studies using therapeutic dosing regimen in three different MASH animal models with different etiology – the mouse STAM (streptozotocin + high-fat-diet), the DIAMOND (isogenic BL6/129 hybrid strain fed an obesogenic Western diet supplemented by high fructose and sucrose), as well as rat CDHFD (choline-deficient high-fat diet) models. In a 6-week-long STAM model (Fig. 1A), the therapeutic regimen of an administration of OATD-01 for 4 weeks at 100 mg/kg QD led to a systemic decrease in chitinolytic activity in the serum (Fig. 1B) resulting in improved all pathological changes in the liver (Fig. 1C-K). We have therefore provided evidence for the *in vivo* systemic target engagement and a positive effect of CHIT1 pharmacological inhibition determined by a histological assessment of MASH severity. Specifically, liver sections were evaluated for steatosis, lobular inflammation, and hepatocyte ballooning, which combined give the MAS score (Metabolic dysfunction-associated steatotic liver disease Activity) Score (Kleiner et al., 2005). OATD-01 administration improved all of these parameters (Fig. 1C-K). Inhibition of inflammation was assessed by reduced H&E staining (Fig. 1F, G), while for the analysis of fibrosis, we utilized computer-aided analysis of Picro Sirius Red (PSR)-stained sections (Fig. 1H, I). Since CHIT1 is mainly expressed in macrophages, we have performed immunohistochemical (IHC) staining detecting murine macrophage marker F4/80 (Fig. 1J, K). Quantitative analysis of images showed that OATD-01 decreased positive area fraction, which represents reduced infiltration of the liver with monocyte-derived macrophages. Overall, treatment with OATD-01 resulted in the improvement of liver morphology in the STAM model.

**Figure 1.**
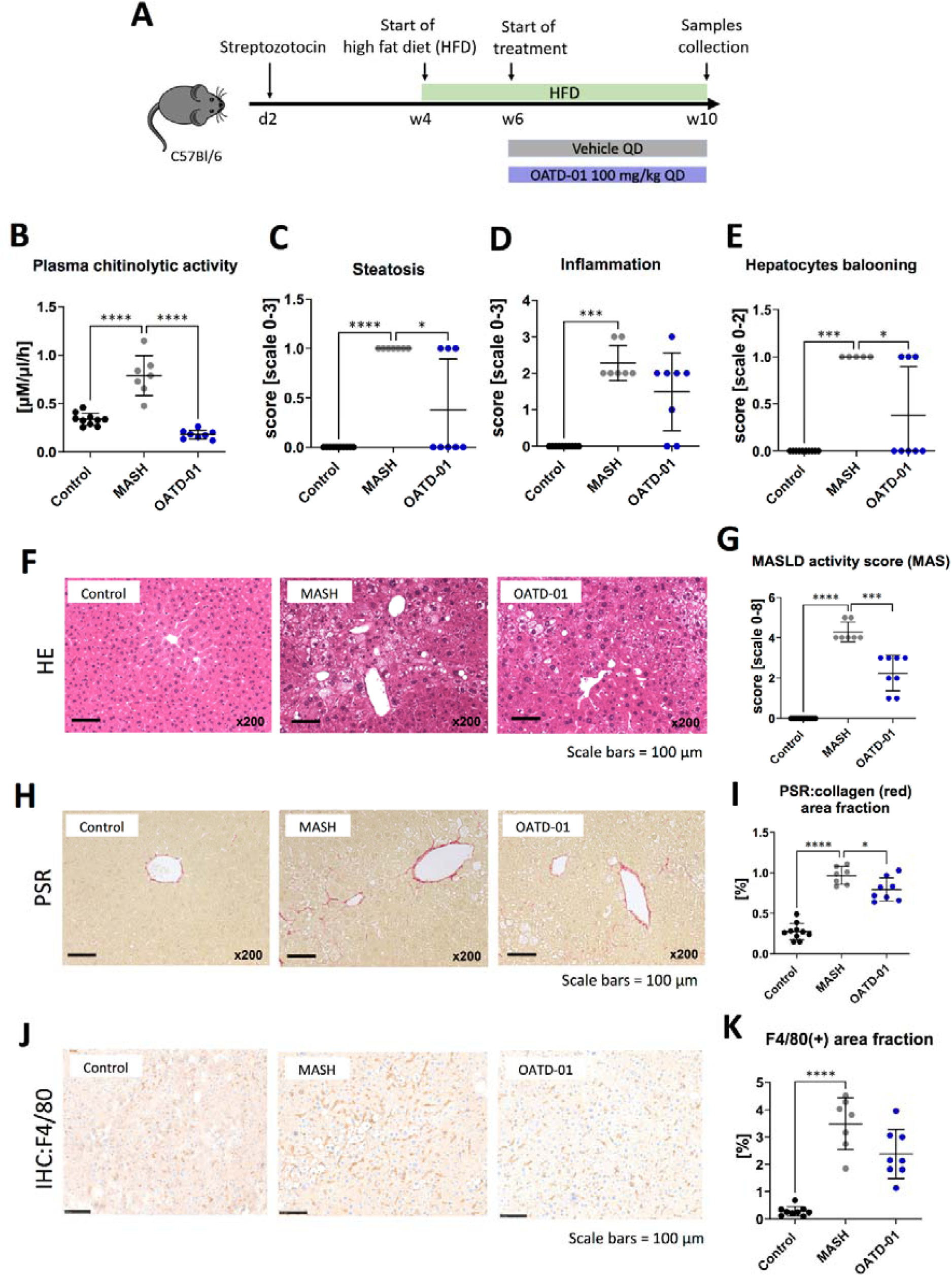
OATD-01 reduces MASH features in a 6-week-long STAM model (high fat diet treated animals with destroyed pancreatic islet’s beta cells). Datasets were collected from three groups: non-disease (depicted as control, black dots), STAM treated with vehicle (MASH, gray dots), and treated with OATD-01 at 100 mg/kg QD for 4 weeks (OATD-01, blue dots). In all graphs, data are presented as individual points with a median depicted with a 95% confidence interval, unless stated otherwise. Statistical significance of differences between groups: * p < 0.05, ** p < 0.01, *** p < 0.001, **** p < 0.0001. Scale bars reference for histological images are depicted below panels. (A) Scheme of the study. (B) Activity of chitinases in the serum. Data presented as means and standard deviation. (C)-(F) Histopathological evaluation of MASH severity, according to the Kleiner system, in hematoxylin and eosin (HE) stained sections of livers from animals in the study: steatosis (C), lobular inflammation (D) and hepatocytes ballooning (E). These scores sum up to the MAS score (metabolic dysfunction-associated steatotic liver disease activity score) (G). Representative pictures of HE stained sections (F). (H)-(I). Analysis of fibrosis burden in livers. Representative pictures of picrosirius red-stained section of livers from experimental groups (H). In red – depositions of collagen I and III. On graph (I) computer-aided analysis of red-positive area in pictures taken from the specimens (n=5 fields/animal, 200x magnification), presented as percent of field-of-view covered by red collagen (performed with ImageJ). Data presented as means and standard deviation; (J)-(K) Analysis of liver F4/80(+)-macrophages. Representative pictures of immunochemically (IHC) detected F4/80(+) cells in sections of livers from experimental groups (J). In brown–positive cells. Graph (K) shows a computer-aided analysis of the brown-positive area in pictures taken from the specimens (n=5 fields/animal, 200x magnification). Data are presented as percent of field-of-view covered by positive signal (performed with ImageJ). Data presented as means and standard deviation.

In a 24-week-long DIAMOND model (Fig. 2A), mice were fed with a high fat, high sucrose, and high cholesterol chow (Western Diet) and therapeutically administered with OATD-01 at 100 mg/kg QD for 16 weeks. Overall, the systemic changes, as determined by the analysis of serum biomarkers, showed that compound dosing improved the activities of ALT and AspAT, as well as levels of cholesterol and triglycerides (Fig. 2B-E). Moreover, in livers, OATD-01 reduced expression of MASH-related genes, namely *Tnf, Col1a1, Timp1, and Mmp9* (Fig. 2F-I). The beneficial effects of OATD-01 were in line with the abolished chitinase activity in the serum (Fig. 2J). After administration of OATD-01 histological assessment of livers revealed reduced all three principal pathologies in MASH – steatosis, lobular inflammation and hepatocyte ballooning (Fig. 2K-O), consequently, reducing combined MAS score (Fig. 2O). Assessment of liver fibrosis was determined on PSR-stained sections (Fig. 2P), using a simple scoring system (0-3) and demonstrated a reduction trend (Fig. 2Q). Overall, administration of OATD-01 in the DIAMOND model alleviated MASH severity, measured by serum biomarkers and liver MASH-related gene expression and histopathology.

**Figure 2.**
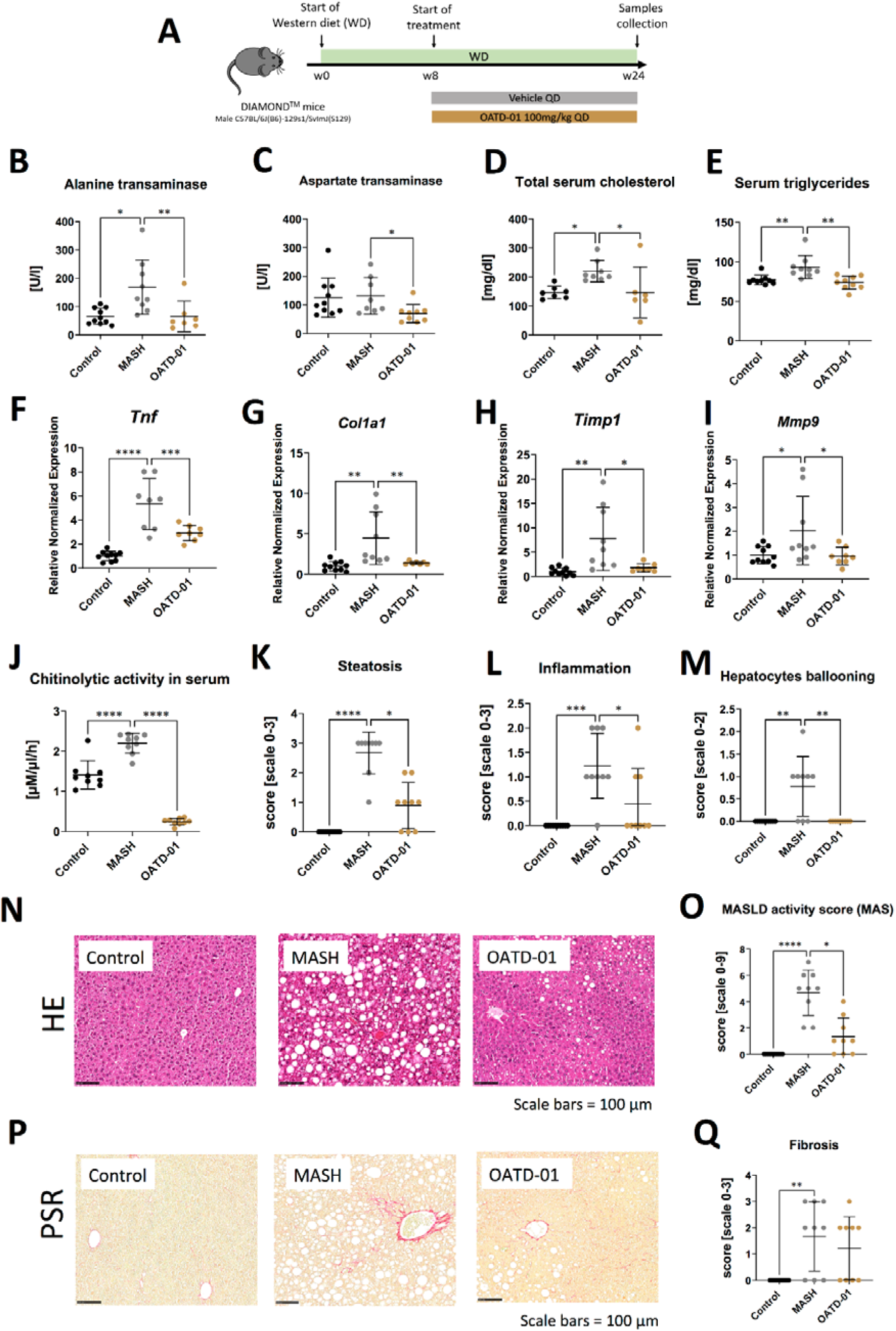
OATD-01 reduces MASH features in a murine model of a high fat, high sugar, and high sucrose diet for 24 weeks (DIAMOND model). Datasets are collected from three groups: healthy animals (depicted as controls, black dots), diet-fed DIAMOND mice treated with vehicle (MASH, gray dots) or treated with OATD-01 at 100 mg/kg QD for 16 weeks (OATD-01, brown dots). In all graphs, data are presented as individual points with a median depicted with a 95% confidence interval, unless stated otherwise. Statistical significance of differences between groups: * p < 0.05, ** p < 0.01, *** p < 0.001, **** p < 0.0001. Scale bars reference for histological images are depicted below panels. 0 (A) Scheme of the study. (B)-(E) Measurements of liver-related markers in serum: activity of ALT (B) and AspAT (C), total cholesterol (D), triglycerides (E). (F)-(I) Expression of genes involved in the development and progression of MASH in liver tissue collected from animals in the study: *Tnf* (F), *Col1a1* (G), *Timp1* (H), *Mmp9* (I). Datasets are normalized to the mean value from the control group. Data are presented as means with standard deviations. (J) Activity of chitinases in serum. (K)-(O) Histopathological evaluation of MASH severity, according to the Kleiner system, in hematoxylin and eosin (HE) stained sections of livers from animals in the study: steatosis (K), lobular inflammation (L) and hepatocytes ballooning (M). These scores sum up to the MAS score (O). Representative pictures of HE stained sections from control animals are shown on (N). (P)-(Q) Analysis of fibrosis burden in livers. Representative pictures of PSR-stained section of livers from experimental groups (P). In red – depositions of collagen I and III. On graph (Q) analysis of the scoring of fibrosis acc. to Kleiner guidelines.

Having determined the efficacy of OATD-01 in murine MASH models, we have tested its effectiveness in a rat CDHFD model. Rat is a species where recent mechanistic insights into metabolism facilitated the selection of biomarkers consistent with human biology and demonstrated a strong resemblance to human biochemical networks (Blais et al., 2017). Having established pharmacokinetic and pharmacodynamic properties of OATD-01 in rats and activity toward rat CHIT1 enzyme, we have moved towards efficacy assessment in the rat MASH model. Specifically, we verified the efficacy of OATD-01 in the CDHFD model, using a TGFβ1 receptor inhibitor, ALK5i, as a traditional antifibrotic reference drug to investigate anti-fibrotic properties of OATD-01 in the context of metabolic liver disease. Both drugs were administered in the therapeutic regimen for 6 weeks (Fig. 3A). OATD-01 and ALK5i demonstrated efficacy in all measured parameters such as liver-to-body weight ratio (Fig. 3B), serum biomarkers – AspAT (Fig. 3C), as well as biomarkers from ELF panel (enhanced liver fibrosis – measured serum levels of procollagen 1, TIMP-1 and hyaluronian) (Fig. 3D-F). Moreover, the evaluation of exposure to OATD-01 was performed in rats under steady-state conditions in week 8^th^ of the study (day 14^th^ of treatment with OATD-01). Upon once-daily administration at a dose of 100 mg/kg, OATD-01 exhibited high plasma exposure, with free compound concentrations exceeding the IC_90_ for the inhibition of the target enzyme CHIT1, even at trough levels (Fig. 3G-H).

**Figure 3.**
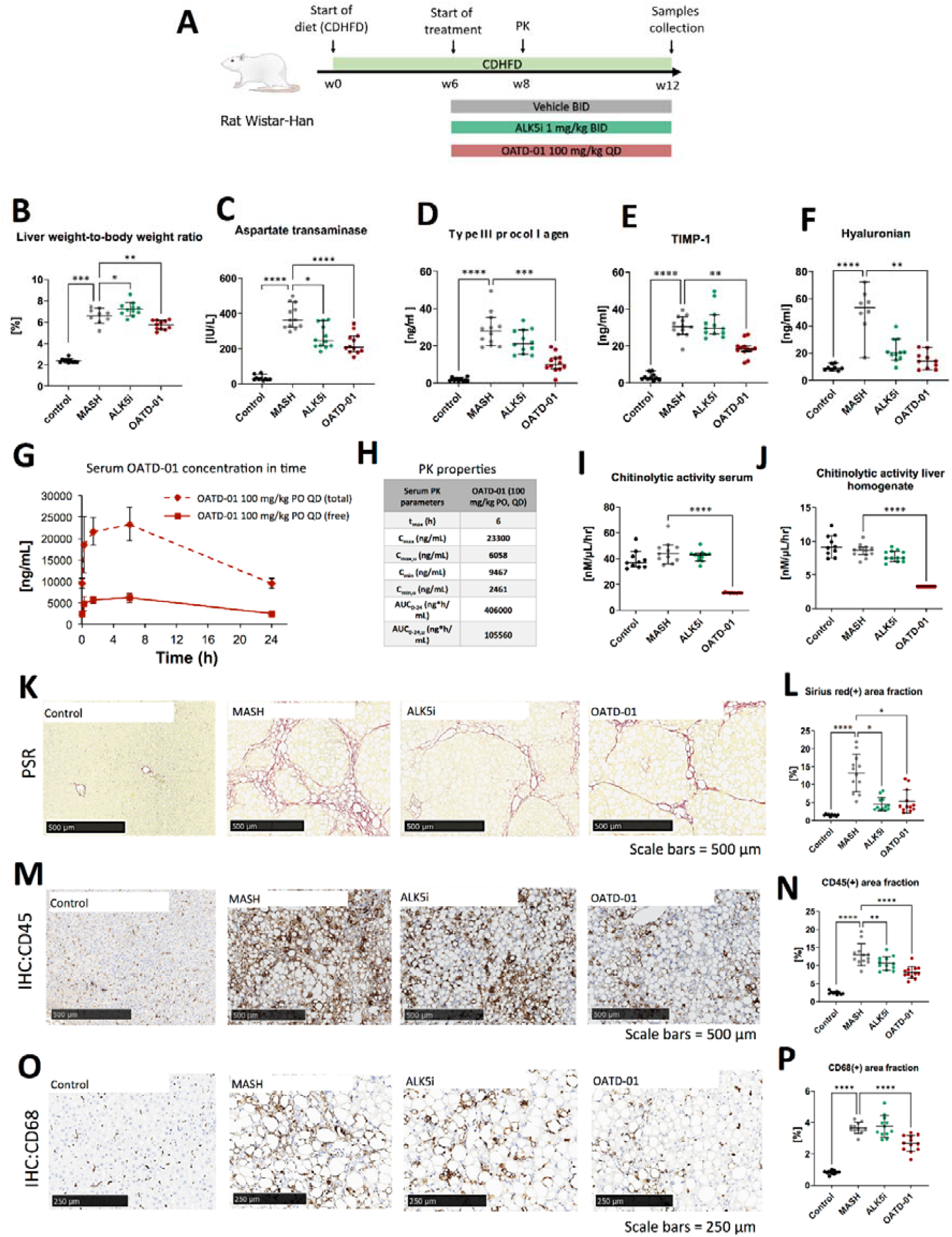
OATD-01 reduces MASH symptoms in a 12-week-long, choline-deficient, high fat diet-induced (CDHFD) rat model on MASH. Datasets were collected from three groups: non-disease animals (depicted as control, black dots), animals fed with the diet and treated with vehicle (MASH, gray dots), treated with AKL5i (1 mg/kg BID for 6 weeks, green dots) and administered with OATD-01 at 100 mg/kg BID for 6 weeks (OATD-01, purple dots). In all graphs, data are presented as individual points with a median depicted with a 95% confidence interval, unless stated otherwise. Statistical significance of differences between groups: * p < 0.05, ** p < 0.01, *** p < 0.001, **** p < 0.0001. Scale bars reference for histological images are depicted below panels. (A) Scheme of the study. (B) Liver weight-to-body weight ratio in rats in the study. Data presented with means with standard deviation. (C) Aspartate transaminase (AspAT) activity in serum collected from experimental animals. (D)-(F) Measurements of ELF (Enhanced Liver Fibrosis) system parameters in serum of experimental animals: type III procollagen peptide (PIIINP)(D), TIMP-1 (E), hyaluronian (F): (G)-(H) Pharmacokinetic properties of OATD-01 in the model, assessed by measurements of compound concentration in serum on the 14th day of treatment (week 8 of the study) with OATD-01 at 100 mg/kg BID, at timepoints: pre-dose, 0.25, 1.5, 6 and 24 hours post-dosing (n=3, sequential blood collection). In plot (G) the measured total concentration is presented as a solid line; the dotted line represents the “free fraction” of the compound, recalculated by considering plasma protein binding. These animals did not receive the second dose of the compound that day. Summary of the calculated PK properties in table (H); (I)-(J) Activity of chitinases in serum (I) and in liver homogenates (J) from animals in the study. (K)-(L) Analysis of fibrosis burden in livers from experimental animals. Representative pictures of the picrosirius red (PSR) stained section are presented in (K). In red – depositions of collagen I and III. In graph (L) whole slide computer-aided analysis of the red-positive area is presented as the percent of the liver section covered by red collagen (performed with Visiopharm Image Analysis Software). (M)-(N) Analysis of inflammation in livers from experimental animals. Representative pictures of immunochemically (IHC) detected CD45(+) cells in the section are presented in (M). In graph (N) the whole slide computer-aided analysis of the brown-positive area is presented as a percent of the liver section covered by a positive signal (performed with Visiopharm Image Analysis Software). (O)-(P) Analysis of CD68(+) macrophages infiltration of livers from experimental animals. Representative pictures of immunochemically (IHC) detected CD68(+) cells in the section are presented on (O). In brown–positive cells. In graph (P) whole slide computer-aided analysis of the brown-positive area is presented as the percent of liver section covered by positive signal (performed with Visiopharm Image Analysis Software).

To further characterize the pharmacodynamic effects of OATD-01, chitinolytic activity determination was employed as a direct endpoint, providing a reliable measure of CHIT1 inhibition. The results of the study demonstrate that OATD-01 reduced chitinolytic activity to the limit of detection in both serum and liver homogenates, confirming its primary pharmacological effects on chitinases (Fig. 3I-J). To assess the morphology of tissue samples, we performed a histological examination of fibrosis and cellular infiltration of tissue with immune cells (Fig. 3K-P). The whole slide computer-aided analysis of Picro Sirius Red-positive area showed the antifibrotic effect of OATD-01 as well as ALK5i treatment in the model (Fig. 3K-L). OATD-01 exhibited a stronger anti-inflammatory effect as seen by measurements on IHC:45(+)-stained sections (Fig. 3M-N). Importantly, analysis of IHC:CD68(+)-stained sections illustrated that administration of OATD-01, but not ALK5i reduced the number of liver macrophages in the tissue providing mechanistic differentiation between two compounds (Fig. 3O-P). Taken together, OATD-01 possesses antifibrotic and anti-inflammatory properties and inhibits in vivo macrophage accumulation proving its immunomodulatory potential for the treatment of MASH.

### MASH-specific gene signature is reversed by the treatment with OATD-01

To better understand genes and pathways involved in the beneficial effect of OATD-01 in MASH, we performed the bulk RNA-seq analysis from liver samples. We investigated the differential expression of genes from three rat groups: controls, MASH and MASH administrated with OATD-01. We have identified 6267 differently expressed genes (DEGs) (|log2(FC)| > 1, p-adjust < 0.05) between MASH and control rats. 4221 of them were up-regulated, while 2046 were down-regulated. Comparing MASH rats with the group treated by OATD-01 (MASH + OATD-01), the number of DEGs was 1304 with 229 up-regulated and 1075 down-regulated. Importantly, 956 out of 1075 (89%) of down-regulated genes after OATD-01 treatment were up-regulated in the MASH group, demonstrating that restorative process of OATD-01 observed by histopathological analysis is reflected by a clear reversal of a transcriptomic profile associated with MASH.

To verify how the rat CDHFD model resembled human MASH, we cross-referred 25 human biomarkers genes, whose expression is strongly associated with MASH (Govaere et al., 2020) with a dataset obtained from this experiment. 20 of human biomarkers have rat ortholog genes and all of them exhibited increased expression in the rat CDHFD model compared to controls, which is consistent with human MASH. Strikingly, the expression of all 20 biomarkers was reversed in rats treated with OATD-01 (Fig. 4A-B). These findings suggested that our CDHFD rat model was MASH-relevant and that further studies that aim to explore the effectiveness of OATD-01 in this context were translationally justified.

**Figure 4.**
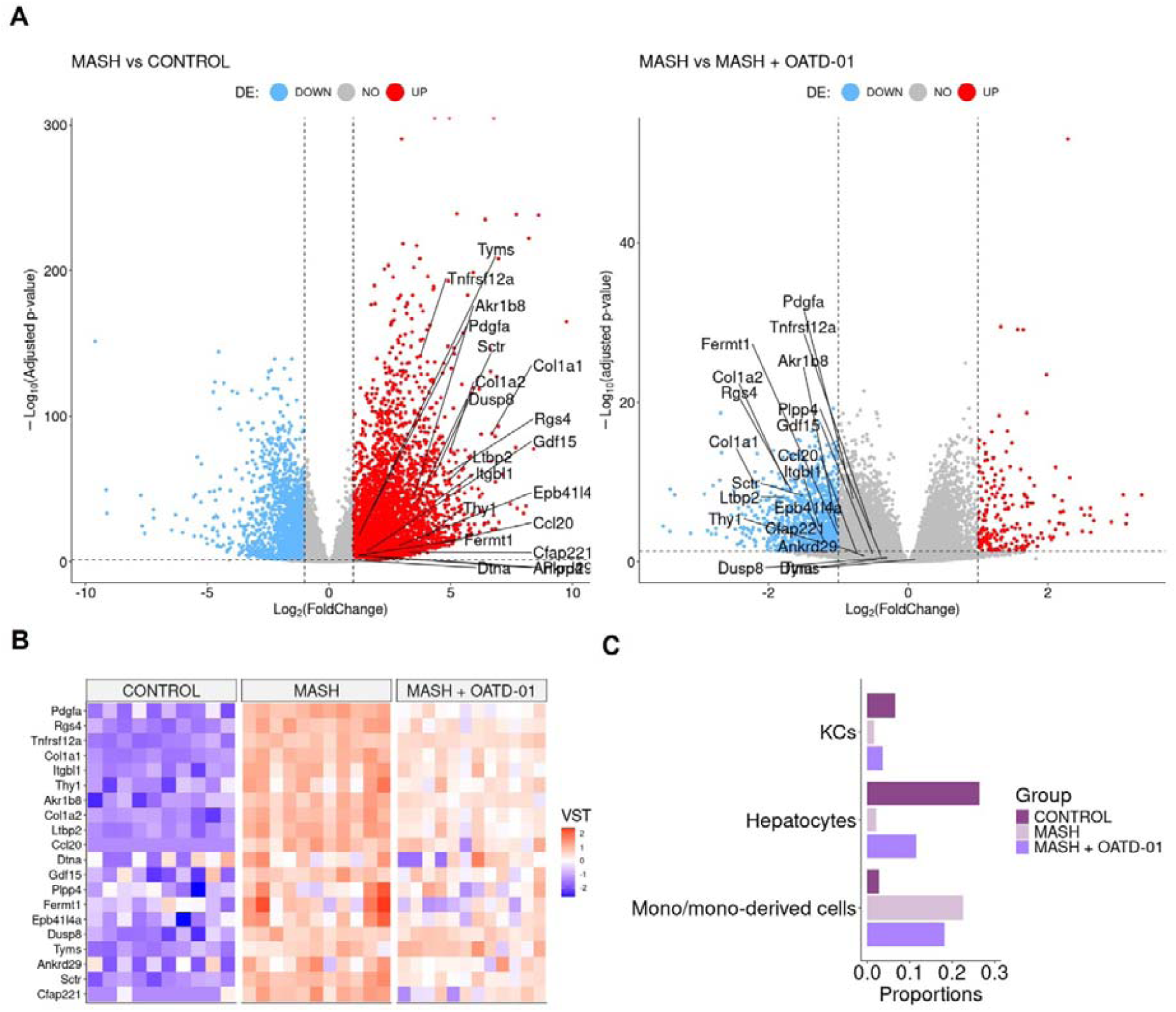
OATD-01 reverses the gene expression of MASH biomarkers and mitigates changes in cellular composition caused by MASH. (A) Volcano plots of differentially expressed genes in the MASH group vs Control group (on the left) and in the MASH group vs the MASH + OATD-01 group (on the right) in the MASH rat study. Biomarkers related to human MASH study are labeled. (B) Heatmap of 20 orthologs of human MASH-related biomarkers expression comparison between the three groups. Each column represents a single individual, and each row represents a gene expression measured in a scaled VST metric, which indicates the direction of expression changes. (C) Selected cell type populations derived from RNA-seq deconvolution results obtained from the livers of control, MASH, and MASH+OATD-01 rats. The full RNA-seq deconvolution results are shown in Supplementary Fig. 1.

As a next step we have performed RNA-seq deconvolution to determine the cellular composition of OATD-01-protected MASH livers (Fig. 4C). The analysis revealed that cholangiocytes, hepatocytes, endothelial cells, and monocyte-derived cells are major contributors to the cellular profile of the rat liver samples (Supplementary Fig. 1). Most significant changes were noted in the proportions of monocyte-derived cells between MASH livers and those treated with OATD-01. We confirmed that the protective effect of OATD-01 manifested in decreased monocyte-derived cells in the MASH livers (Fig. 4C), as previously shown by IHC staining (Fig. 3P). Additionally, healthier MASH livers after OATD-01 treatment had increased hepatocytes level (Fig. 4C).

### Functional enrichment analysis uncovers metabolic switch upon OATD-01 treatment

To define biological pathways induced in CDHFD-treated rats (MASH group) and reversed by OATD-01 treatment, we performed Gene Ontology (GO) functional study and Kyoto Encyclopedia of Genes and Genomes (KEGG) pathway enrichment analysis by applying the GSEA procedure. In the MASH group we found 447 significant enriched biological processes described in GO (31 KEGG pathways) and 394 processes in the OATD-01 treated group (48 in KEGG pathway) (Supplementary Tab. 1). Next, we performed a procedure of clustering the processes based on leading genes shared between processes (for details see Supplementary material on hierarchical clustering). These genes have the greatest influence on the enrichment score. It is evident that OATD-01 effectively reversed most clusters involved in MASH disease (Fig. 5A, B).

**Figure 5.**
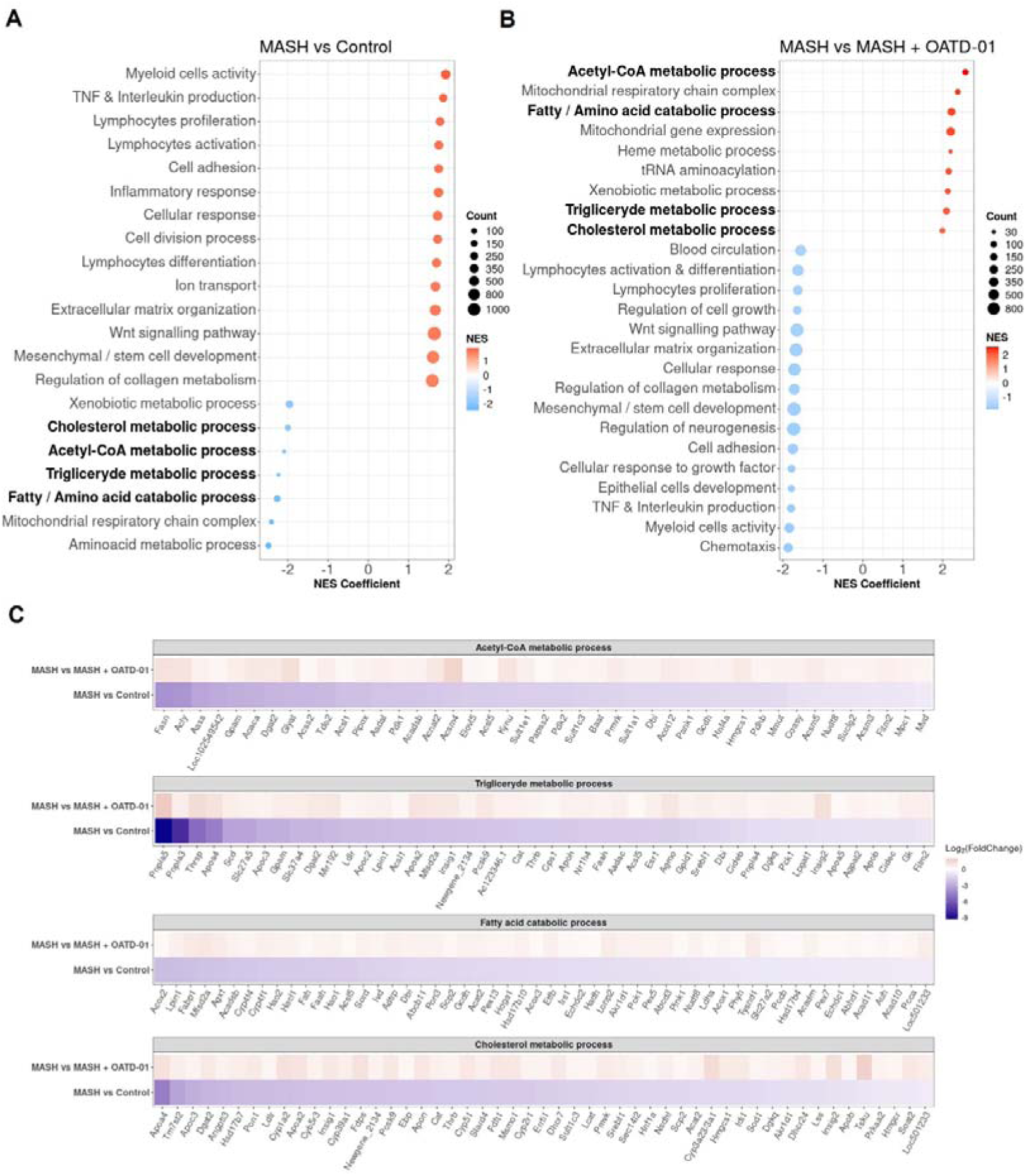
Clustering of GO biological processes from GSEA analysis shows that OATD-01 has a reversal effect, significantly influencing metabolism in the MASH rat study. (A, B) Clustered significantly changed processes in MASH vs Control (A) and MASH vs MASH + OATD-01 (B). NES describes the gene enrichment of the selected process in the gene ranking list concerning the permuted order of these genes. Here, the NES coefficient indicates the median NES of the cluster. Positive NES means that leading genes from the cluster are mostly upregulated, while negative NES indicates downregulation of the most leading genes from the cluster. The interesting, clustered processes related to lipid metabolism are highlighted. (C) Heatmaps showing the expression of leading genes in clustered processes are highlighted in (A, B). Gene expression changes are presented using log2(FoldChange) and visualized through a color gradient where blue indicates strong downregulation, and red indicates strong upregulation.

Based on the NES coefficient, which measures gene activation, the top 2 clusters with increased gene activity (NES >0) in MASH group were identified as *Myeloid Cell Activity* and *TNF & Interleukin production* (Fig. 5A) which reflect inflammatory aspects of this disease. The fibrotic processes in MASH were indicated by the upregulated expression of genes clustered as *Extracellular matrix organization* and *Regulation of collagen metabolism.* A significant proportion of genes in these clusters were reversed by the OATD-01 treatment, aligning with histologically confirmed anti-inflammatory and anti-fibrotic effects of the drug (see Venn diagrams in Supplementary Fig. 2; Fig. 3K-P). Strikingly, we identified 7 metabolic clusters as significantly decreased (NES <0) in MASH condition and reversed after OATD-01 treatment (Fig. 5A, B). We have drawn similar conclusions after GSEA analysis taking into consideration molecular functions, cellular components and KEGG pathways categories (Supplementary Tab. 2).

4 out of 7 metabolic clusters named *Acetyl-CoA metabolic process, Triglyceride metabolic process, Fatty/amino acid catabolic process and Cholesterol metabolism* were linked to lipid metabolism, dysregulation of which is pivotal in the pathogenesis of MASH (Rao et al., 2023). Therefore, we conducted a detailed analysis of the expression of leading genes in the aforementioned clusters, with an additional focus on the fatty acid catabolic process subcluster within the broader fatty/amino acid catabolic process (Fig. 5C). We observed that in the MASH group, there is a general decrease in the activity of genes involved in acetyl-CoA synthesis (e.g. *Pdhb, Acaca, Acly*). Specifically, a reduction in the utilization of this metabolite for de-novo fatty acid and triglyceride synthesis (e.g. *Fasn*, *Dgat2, Lpin1*), as well as fatty acid catabolism through oxidation in peroxisomes or mitochondria (e.g. *Acox2, Hao2, Hsd17b10*). Similar changes were observed for genes important for cholesterol synthesis (e.g. *Tm7sf2, Fdps, Srebf1*). The cholesterol metabolism pathway examined in the KEGG database revealed dysregulation of genes involved in cholesterol flux and lipoprotein packing (Supplementary Fig. 3). Notably, OATD-01 reversed the increased expression of *Abca1*, which is involved in cholesterol flux, *Lpl* - important for the hydrolysis of lipoproteins (Basso et al., 2003; Wu et al., 2021) and *Cd36* gene - implicated in fatty acids uptake from lipoprotein hydrolysis (Rada et al., 2020). CHIT1 inhibition restored the expression of the *Ldlr* gene, which is downregulated in MASH and crucial for the degradation of low-density-lipoproteins (LDLs) (Bieghs et al., 2012), indicating its effect on cholesterol cycling. Overall, we observed impaired recycling of acetyl-CoA metabolite and cholesterol flux in MASH conditions. OATD-01 treatment restored the expression of most genes in clusters related to lipid metabolism, as well as those involved in cholesterol recycling, thereby restoring proper lipid metabolism in the model.

### OATD-01 inhibits glycolysis in MASH condition and in activated BMDMs

Recognizing the effect of OATD-01 on genes involved in the synthesis of acetyl-CoA, like *Pdhb*, which is part of pyruvate dehydrogenase complex (Yonashiro et al., 2018), we wondered how glycolysis is altered in MASH and upon OATD-01 treatment. Our interest stems from the known influence of glycolysis on inflammation and repair processes during MASH progression (Li et al., 2023; Qu et al., 2023). We specifically examined the expression of genes assigned to the glycolysis pathway in the KEGG database. Indeed, gene expression profiles suggested alterations in glycolysis under MASH conditions (Supplementary Fig. 4). Specifically, among 62 genes assigned to the glycolytic pathway, as much as 18 were up-regulated (e.g. *Hk1, Pkm*) and 15 were down-regulated, including hepatocyte-expressed genes like e.g. *Gck1, Pklr* (Supplementary Fig. 4 and Fig. 6A), which can be aligned with the loss of hepatocytes in MASH condition and upregulation of the glycolytic pathway in MASH. OATD-01 treatment reversed the expression pattern of 18 genes, including the MASH-upregulated rate-limiting genes such as *Hk1-3* and *Pkm* and also *Slc2a1 -* which encodes glucose transporter 1 (GLUT1) involved in glucose uptake (Karim et al., 2014) (Fig 6A). In addition, the MASH-elevated expression of *Ldhb,* indicating anaerobic glycolysis, was lowered down by OATD-01 (Fig 6A). This data indicates a significant inhibitory effect of OATD-01 on the glycolysis pathway (Fig 6A).

**Figure 6.**
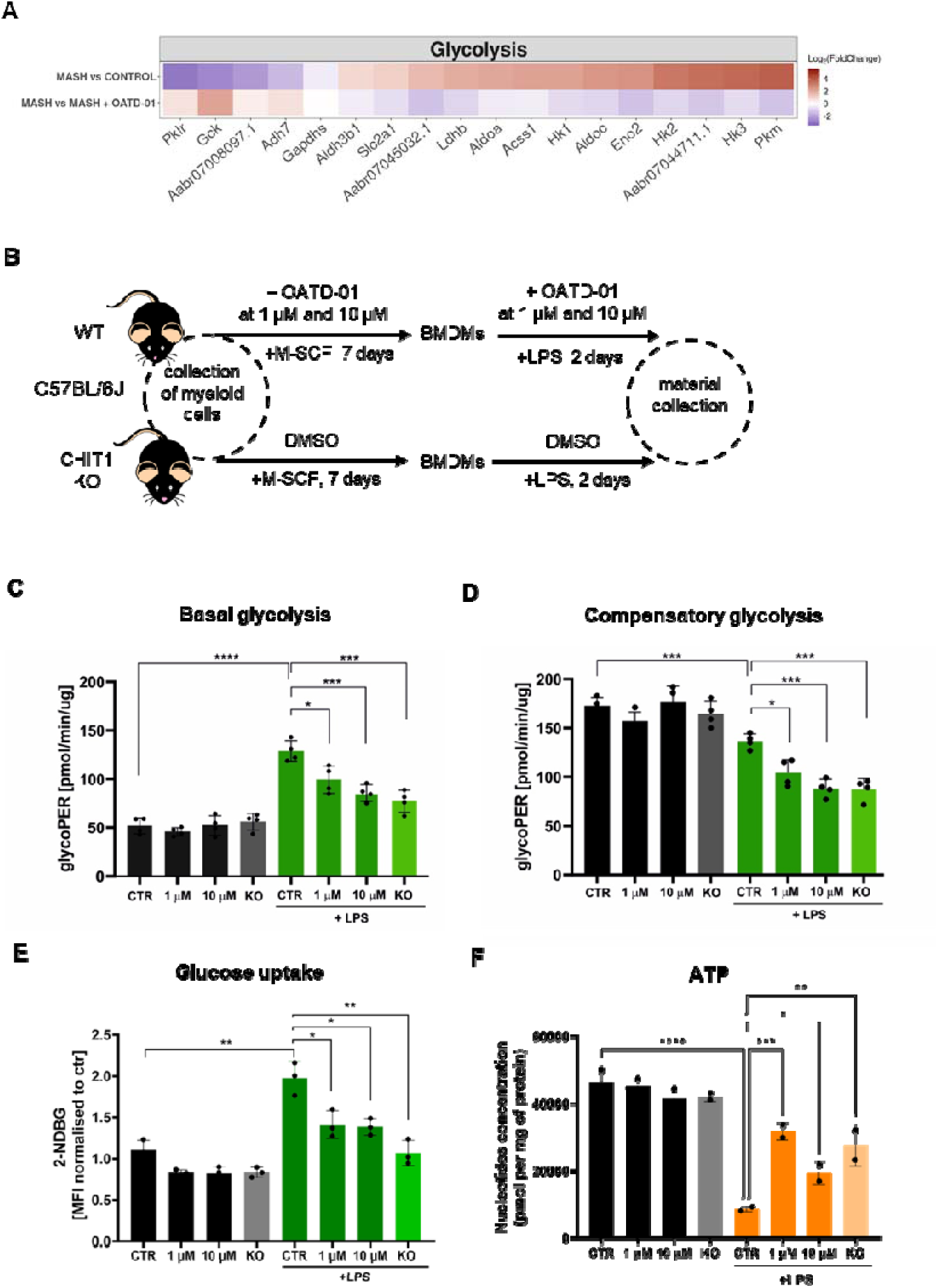
OATD-01 inhibits glycolysis in MASH rat study and in LPS-treated BMDMs. (A) Heatmaps displaying the expression of genes involved in glycolysis, altered in MASH (|log2(foldChange)| >1) and reversed by OATD-01. Genes were retrieved from KEGG database. Gene expression changes are presented using log2(FoldChange) and visualized through a color gradient where blue indicates strong downregulation, and red indicates strong upregulation. (B) Experimental protocol scheme for bone marrow-derived macrophages (BMDMs). BMDMs were derived from WT and CHIT1 KO mice. Upon collection of bone marrow cells, WT cells were treated with OATD-01 at two doses (1 µM and 10 µM) or with DMSO as a control during the differentiation period and polarization with LPS for 48h until material collection. Non-LPS treated cells were also prepared as a control of metabolic changes. (C) Measurement of basal glycolysis (n=2 biological replicates, each with two technical replicates) and (D) compensatory glycolysis (n=2 biological replicates, two technical replicates per biological replicate) by Seahorse assay, (E) assessment of glucose uptake with 2-NDB-labeled glucose (n=3 biological replicates), and (F) quantification of levels of total ATP in 48h LPS-stimulated BMDMs (n=2 biological replicates) prepared as described in (B). Data presented in (C-F) are shown as means and standard deviation. The statistical test used in (C, E) is an unpaired t-test, while in (F) it is a one-way ordinary ANOVA with Fisher’s Least Significant Difference (LSD) test for multiple comparisons. Statistical significance between groups is indicated as follows: * p < 0.05, ** p < 0.01, *** p < 0.001, **** p < 0.0001.

Since the transcriptomic data clearly shows that OATD-01 is inducing metabolic switch in MASH livers, we decided to test this functionally on CHIT1-originating cells, whose numbers are differentially regulated in our model, namely macrophages. Macrophages are an excellent cell type for this study - when stimulated with a proinflammatory stimulus like LPS, they switch their metabolism from oxidative to glycolytic, shutting off the TCA cycle (Cheng et al., 2014; Palsson-Mcdermott et al., 2015). We used bone marrow-derived macrophages (BMDMs) to establish the effect of CHIT1 on their metabolism by pharmacological (OATD-01) and genetic inactivation (CHIT1-KO) (Fig. 6B). Previous studies have shown that CHIT1 expression peaks during macrophage differentiation (Di Rosa, Malaguarnera, et al., 2013). Therefore, after isolating bone marrow cells, we administered OATD-01 and maintained it in the culture medium throughout the differentiation process until the end of polarization (Fig. 6B) with no effect on macrophage activation markers (Supplementary Fig. 5). Next, we used an extracellular flux analyzer to examine the effect of CHIT1 inhibition on glycolysis in naive and LPS-polarized BMDMs over 48 hours (Fig. 6C-D). As expected, LPS-treated macrophages displayed increased basal glycolysis (Fig. 6C). OATD-01 treatment dose-dependently reduced both basal and compensatory glycolysis (Fig. 6C-D). The same inhibition was observed in CHIT1-KO BMDMs, demonstrating a target-specific effect (Fig. 6C-D).

Since glycolysis depends on glucose uptake, we investigated this activity with the use of fluorescently labeled glucose (2-NDBG) measured in BMDMs by FACS. (Fig. 6E). The anticipated increase in a glucose uptake in LPS-stimulated BMDMs was dose-dependently reduced by OATD-01, similar to the effect observed in CHIT1-KO (Fig 6E). Keeping in mind that the effect of OATD-01 was only present when the compound was present throughout the macrophage differentiation process, we speculated that the consequences of CHIT1 inhibition on macrophage phenotype are not rapid. To explain the effect of OATD-01 on glucose uptake, we tested its impact on GLUT1 transporter expression, since it is a main glucose receptor responsible in macrophages for the facilitation of this process (Cornwell et al., 2023). Membrane protein levels were not affected in OATD-01-treated and CHIT1 KO macrophages (Supplementary Fig. 6) suggesting that OATD-01 acts by regulating GLUT1 activity in macrophages. It is known that GLUT1 activity can be inhibited by ATP (Blodgett et al., 2007). Therefore, we measured ATP levels and found that OATD-01 restored the LPS-induced reduction of ATP (Fig. 6F) keeping cells in a more energetic state.

Overall, OATD-01 inhibits the expression of genes involved in glycolysis in MASH condition, which we directly addressed in functional assays showing the inhibitory effect of our drug on glycolysis and glucose uptake in primary macrophages.

### OATD-01 inhibits immunometabolic changes in activated macrophages

In the next step, we explored how glycolysis inhibition affects downstream metabolites, focusing on those reported to play a key role in inflammation and MASH progression, namely citrate (Amjad et al., 2023; Tan et al., 2020) and acetate (Aoki et al., 2021; Xu et al., 2019). Citrate, originating from the TCA cycle, is a driver in this context (Amjad et al., 2023; Duan et al., 2021; Tan et al., 2020), whereas acetate, generated from pyruvate, has anti-inflammatory and therapeutic benefits (Aoki et al., 2021; Xu et al., 2019). As expected, LPS treatment resulted in an increase in citrate due to the shutdown of the TCA cycle (Fig. 7A). Both CHIT1 inhibition by OATD-01 and genetic deletion decreased citrate levels (Fig. 7A). Additionally, these treatments improved acetate levels, likely indicating a lower glycolytic flux (Fig. 7B). All of the above metabolic changes, initiated by the inhibition of glycolysis, should result in a decrease of IL-1β (Palsson-Mcdermott et al., 2015). Indeed, IL-1β secretion was lower in both CHIT1 KO and OATD-01-treated BMDMs (Fig. 7C). Altogether, the inhibition of CHIT1 diminished the metabolic switch that occurs in activated macrophages and in consequence reduced inflammation, as determined by IL-1β - a metabolically regulated cytokine relevant in MASH.

**Figure 7.**
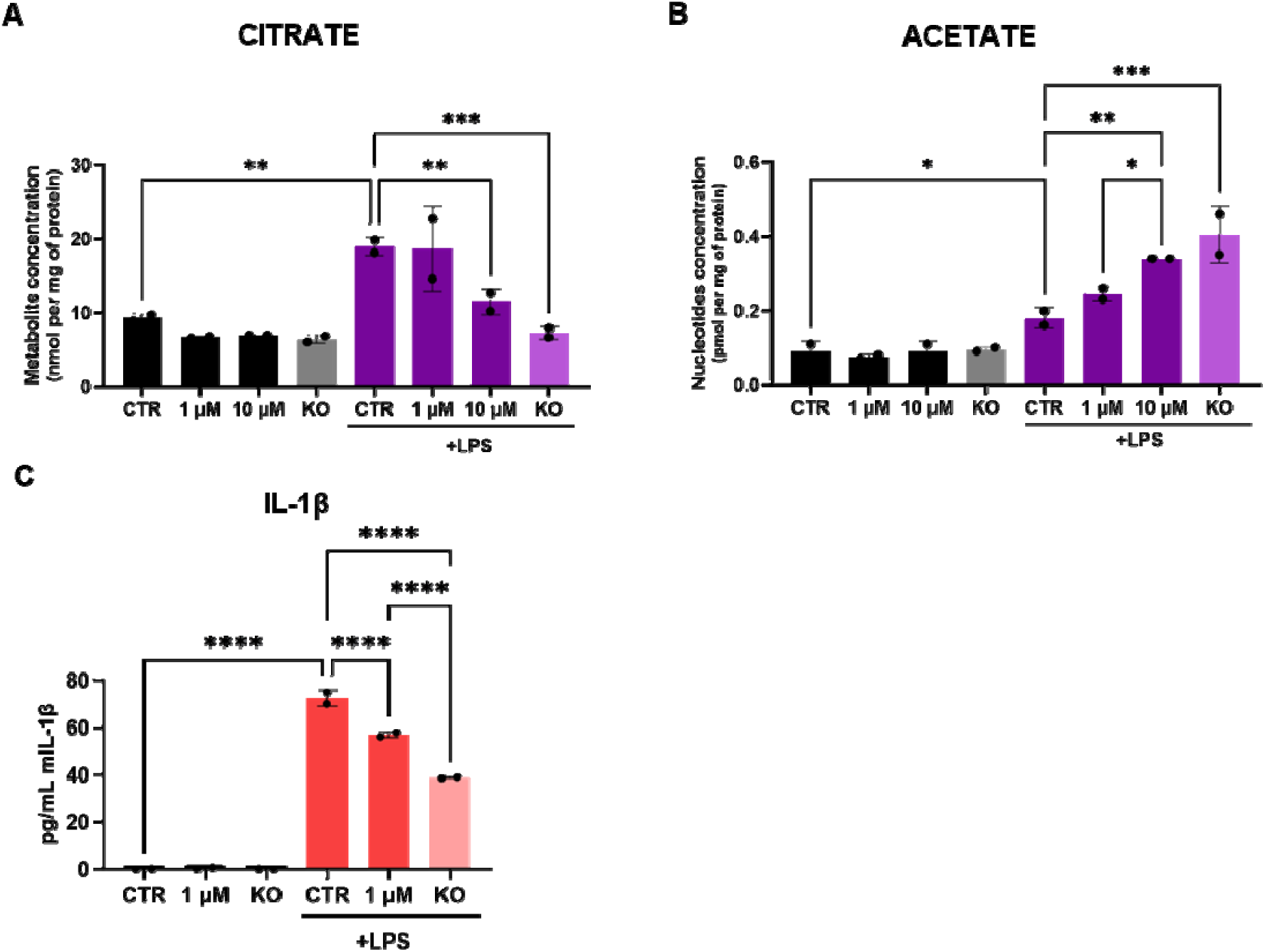
OATD-01 regulates immunometabolism in LPS-stimulated BMDMs. Levels of metabolites: (A) citrate (n=2 biological replicates), (B) acetate (n=2 biological replicates), and (C) levels of cytokine IL-1β (a representative experiment from n=2 biological replicates) measured in BMDMs prepared and stimulated with LPS for 48h as described in Fig. 6B. Data presented are shown as means and standard deviation. The statistical test is a one-way ordinary ANOVA with Fisher’s Least Significant Difference (LSD) test for multiple comparisons. Statistical significance between groups is indicated as follows: * p < 0.05, ** p < 0.01, *** p < 0.001, **** p < 0.0001.

## Discussion

In this study, we demonstrate the therapeutic efficacy of OATD-01, an inhibitor of CHIT1, in different animal models of MASH (STAM, DIAMOND, and CDHFD). We present OATD-01 as a global regulator of metabolic changes in MASH, including glucose uptake, glycolysis, acetyl-CoA metabolism, triglyceride metabolism, fatty acid catabolism as well as flux and biosynthesis of cholesterol (Fig. 8). Functional experiments performed on macrophages revealed that both genetic and pharmacological inactivation of CHIT1 resulted in inhibition of glucose uptake and glycolysis leading to a decreased IL-1β production. This knowledge has helped us to design the primary endpoint for the phase 2 proof-of-concept study for OATD-01 in lung sarcoidosis. We are using 18F-fluorodeoxyglucose positron emission tomography-computed tomography (18F-FDG PET-CT) to determine glucose uptake in lung granulomas reflecting pro-inflammatory activity with macrophages at the center of inflammatory cell clusters (the KITE study; NCT06205121). This is a novel and quantitative way to assess the anti-inflammatory characteristics of a drug whose mechanism of action is rooted in the cellular metabolic switch.

**Figure 8.**
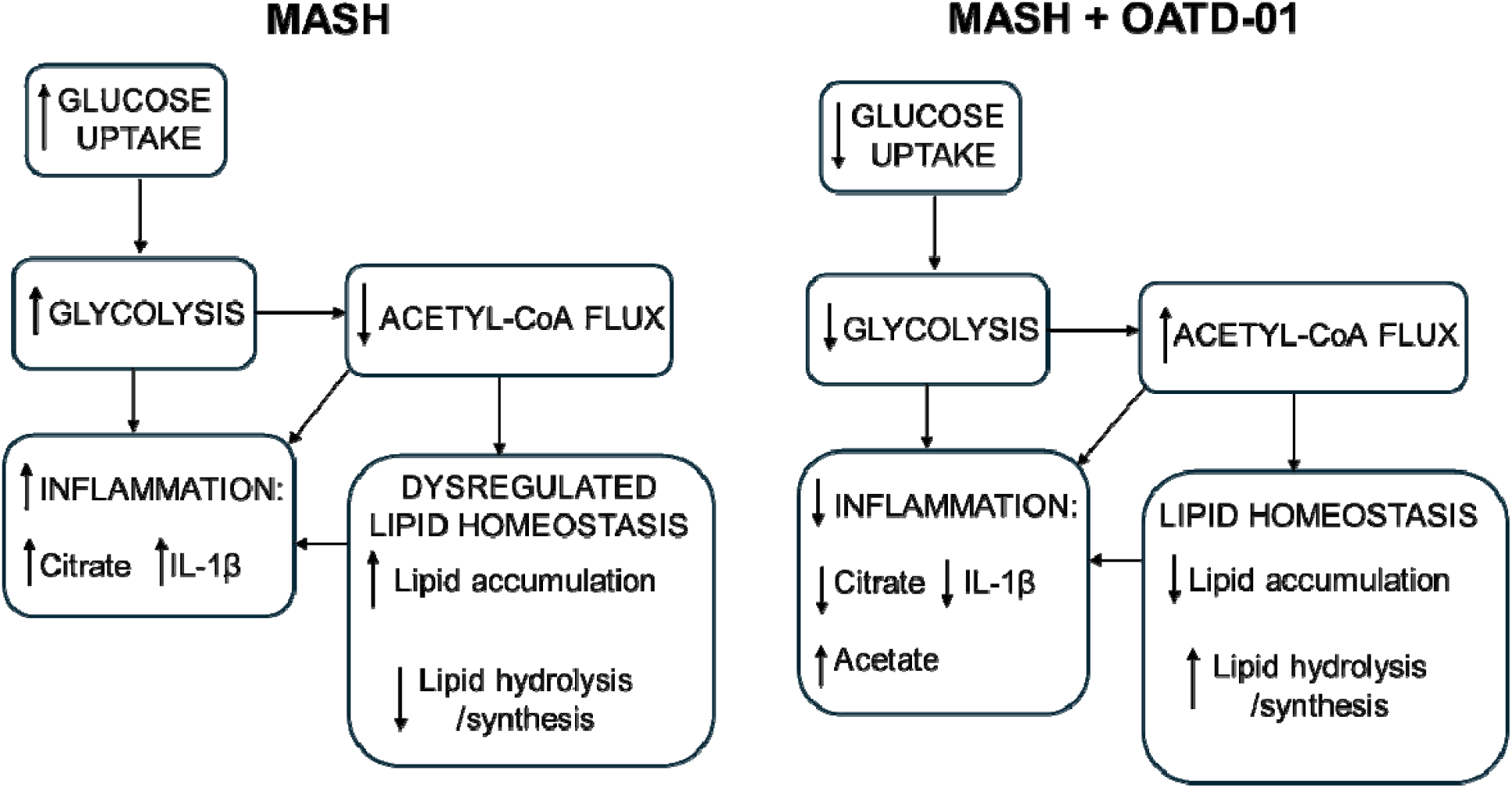
OATD-01 restores proper metabolic homeostasis in MASH condition. A comprehensive model of OATD-01 mechanism of action as a regulator of MASH-related metabolic pathways based on transcriptomic data from MASH rat study as well as metabolic characterization of OATD-01-treated macrophages. In the MASH condition, increased glucose uptake and glycolysis drive inflammation. There is reduced acetyl-CoA flux, leading to citrate accumulation and dysregulated lipid homeostasis, characterized by increased lipid accumulation and decreased lipid processing. This imbalance in lipids may also contribute to heightened inflammation. OATD-01 reverses these processes by directly targeting glycolysis, as observed in macrophages, resulting in decreased inflammation and improved lipid homeostasis.

Our RNA-Seq data mining revealed that the most upregulated pathway by OATD-01 in our GSEA analysis was *Acetyl-CoA metabolism*, which holds genes implicated in the synthesis and utilization of acetyl-CoA for de novo lipid formation. In addition, OATD-01 induced the upregulation of genes like *Acox2* (peroxisomal oxidase) and *Hsd17B10* (mitochondrial dehydrogenase) in *Fatty acid catabolism* cluster, driving the degradation of fatty acids. This suggests that OATD-01 can unlock the flux of acetyl-CoA, restoring the proper balance of lipid metabolism. The *Acox2* gene was highlighted in our study because it encodes a peroxisomal oxidase of branched-chain fatty acids and is critical for hepatic metabolic homeostasis (Y. Zhang et al., 2022). Knockout studies have shown that loss of *Acox2* can lead to liver cancer in mice (Y. Zhang et al., 2022), underscoring the liver-protective role of

OATD-01, which elevated the expression of this gene. Expression of *Acox2* is also induced by peroxisome proliferator-activated receptors (PPARs) and agonists of these receptors are important drugs currently being tested in clinical trials (Staels et al., 2023; Q. Zhang et al., 2021).

Acetyl-CoA synthesis is also the critical first step of the mevalonate pathway of cholesterol biosynthesis. Accumulation of cholesterol is a characteristic of MASH with hypercholesteremia being the major source of toxicity (Song et al., 2021). Cholesterol plays a key role in regulating critical immune processes (Ioannou, 2016) and OATD-01, as an immunomodulatory drug targeting macrophages, can partially exert its anti-inflammatory effect by regulating this pathway. Homeostatic cholesterol biosynthesis is necessary for immune resolution by CD4+ Th1 cells in a c-MAF-AP-1-dependent manner (Perucha et al., 2019), which drives IL-10 production that was shown to be protective against diet-induced fatty liver disease in mice (Cintra et al., 2008). Perturbations in cholesterol biosynthesis in macrophages can directly induce type I interferon signaling in a cGAS-STING-dependent manner (York et al., 2015) and trigger the production of proinflammatory cytokines (Ioannou et al., 2017). In addition, cholesterol accumulation has been implicated as a key stimulus for the activation of the NLRP3 (Duewell et al., 2010) and AIM2 (Dang et al., 2017) inflammasomes in macrophages, the latter occurring as a result of impaired mitochondrial metabolism leading to the release of mitochondrial DNA. Disequilibrium in cholesterol biosynthesis drives multiple inflammatory processes in MASH, identifying cholesterol metabolism as a key target in conditions such as MASH. This phenomenon has already been exploited in patients. Statins used in the clinic (Ayada et al., 2023; Dai et al., 2023) block cholesterol biosynthesis, and restore homeostatic lipid metabolism and mitochondrial bioenergetics (Dai et al., 2023; Park et al., 2016).

In addition to cholesterol biosynthesis, we observed changes in cholesterol flux and recycling induced by MASH and reversed after OATD-01 treatment. OATD-01 restored expression of MASH-depleted *Ldlr,* which encodes the low-density lipoprotein receptor critical for the maintenance of cholesterol homeostasis, and its depletion leads to hypercholesteremia and hepatic toxicity (Ishibashi et al., 1993). Notably, OATD-01 suppressed the MASH-mediated elevation in *Lpl* expression, which is associated with liver fibrosis via the propagation of cholesterol accumulation and inflammatory TLR4 signaling (Teratani et al., 2019). MASH rats exhibited altered expression of genes that regulate cholesterol and phospholipid efflux for the generation of high-density lipoprotein-cholesterol (*Abca1, Apoa1*), indicative of dysregulated cholesterol flux. These effects were limited by CHIT1 inhibition, further indicating the protective effects of OATD-01 via restoration of cholesterol biosynthesis. Another protein that was revealed to be reduced by CHIT1 inhibition was *Cd36*, which plays a key role in fatty acid uptake from lipoprotein metabolism. CD36 is a prognostic marker of MASH and an interesting therapeutic target (Rada et al., 2020).

The processes that we functionally addressed in this study are glucose uptake and glycolysis. These are hallmarks of inflammation and recognized drivers of MASH (Li et al., 2023). The therapeutically exciting interplay between CHIT1 and glycolysis has been shown previously when macrophages treated with 2-deoxyglucose exhibited reduced CHIT1 expression (Suzuki et al., 2016). Looking at downstream signaling we speculate that OATD-01 modulates a metabolic switch in macrophages, dependent on the activation of HIF-1α, a key factor in the pathogenesis of MASH (Chu et al., 2022; Holzner & Murray, 2021). Altered HIF-1α activity further increases glucose metabolism, for example by regulating GLUT1 expression. Possible reversal of HIF-1α stabilization by OATD-01 in MASH can contribute to the observed outcomes such as decreased glucose uptake, glycolysis, and IL-1β production.

OATD-01 showed efficacy in various preclinical models providing net anti-inflammatory and anti-fibrotic effects suggesting that it acts therapeutically on multiple pathways. The exact molecular mechanism is not clear, however, we know that CHIT1, as a glycosidase, can regulate various proteins and lipids glycosylation including those implicated in cellular metabolism. The glycosylation rate depends on the metabolic state, and impaired glycosylation is observed in many inflammatory and metabolic diseases, including MASH (Clarke et al., 2017; Drzewicka & Zasłona, 2024; Ochoa-Rios et al., 2022). Using in vitro macrophage culture experiments we tried to explain the effect of CHIT1 on glucose uptake by regulation of GLUT1 glycosylation. However, we failed to provide convincing evidence for the glycosylation status of GLUT1 upon OATD-01 treatment. LPS-stimulated macrophages might not be the optimal in vitro system to reveal autocrine CHIT1 secretion effects on target glycosylation proteins. In search of global protein targets affected by CHIT1 enzymatic ability to modify glycosylation co-culture system of macrophages and fibroblasts or in vivo experiments may provide the answer.

While we were not able to implicate changes in GLUT1 glycosylation as a direct explanation for the attenuated glucose uptake and glycolysis, thanks to transcriptomics data we have identified other interesting OATD-01 regulated genes. In MASH in vivo study, we have observed an elevated expression of glycolytic genes including *Hk1*, encoding hexokinase 1, encoding the first enzyme in glycolysis, and *Pkm2*, encoding pyruvate kinase M2, the last enzyme in the pathway, shown to be upregulated in macrophages of MASH patients (Qu et al., 2024). Interestingly, the functions of these proteins implicated in glycolysis can depend on glycosylation (Bacigalupa et al., 2018). OATD-01 decreased *Hk1 and Pkm2* expression, indicated restored glucose sensing and therefore decreased glycolysis. This data is supported by a study from Qu et al., 2024, where macrophage-specific genetic deletion of *Pkm2* ameliorated liver inflammation and fibrosis, as a consequence of reduced glycolytic activity.

Another increased glycolytic gene in MASH and reversed by OATD-01 was *Ldhb*, encoding lactate dehydrogenase B (LDHB), indicative of elevated lactate production and tissue damage (James et al., 1999). Hyperacetylation of LDHB enzyme was shown to drive the progression of MASH by impaired clearance of lactate (T. Wang et al., 2021). In our study, the transcriptomic data suggests that lactate production might also be suppressed through an OATD-01-induced reduction of *Ldhb* expression.

Apart from the global effect of OATD-01 on metabolism in MASH models, we provided data on the direct regulation of cellular metabolism in macrophages. Activation of macrophages is characterized by a metabolic switch from oxidative phosphorylation to glycolysis, with decreased TCA cycle activity accompanied by elevated pentose phosphate pathway (PPP) flux and lactate production (Kelly & O’Neill, 2015). During MASH, polarized macrophages exhibit a glycolytic, pro-inflammatory, and pro-fibrotic phenotype. Restoring oxidative phosphorylation and an anti-inflammatory phenotype in macrophages has been shown to limit inflammation and fibrosis in MASH mice (Liu et al., 2023). Inhibition of CHIT1 limits glycolysis and glucose uptake that is accompanied by the accumulation of glycolysis-driven acetate, which can be a substrate for lipid metabolism (Moffett et al., 2020). Acetate is known to reduce inflammation by regulating the inflammasome (Xu et al., 2019). Additionally, it serves as a positive regulator of lipogenesis through the activation of *Fasn, Acaca* genes (Gao et al., 2016), which we observed to be downregulated in MASH and restored after OATD-01. In line with our data, citrate accumulated in macrophages stimulated with LPS (Jha et al., 2015; De Souza et al., 2019). This metabolite is known to amplify inflammatory response through activation of NLRP3 in ALI mice (Duan et al., 2021). Therefore, the effect of the OATD-01 on citrate level can explain the observed reduction in IL-1β production. Citrate is also a clinically important metabolite as circulating citrate levels have been identified as a marker of liver fibrosis in MASH patients (Amjad et al., 2023). The citrate level is regulated by ACLY enzyme, which hydrolyses citrate into oxaloacetate and acetyl-CoA (Dominguez et al., 2021). This enzyme is crucial in regulating macrophage inflammatory responses in vitro (Verberk et al., 2021). In the MASH rat study, a down-regulation of the gene encoding this enzyme, suggests an impairment of citrate levels, which OATD-01 addresses by restoring *Acly* expression levels. Genetic and pharmacological CHIT1 inhibition resulted in a restoration of citrate to the levels observed during homeostasis. This was followed by an increase in TCA cycle flux, confirmed by a significant rise in ATP levels. ATP is a negative regulator of GLUT1 activity; therefore, the diminished glucose uptake is most likely due to the negative feedback loop (Blodgett et al., 2007). This indicates that CHIT1 inhibition can effectively shift metabolism towards a more energetic state, which can be beneficial for the treatment of metabolic disorders, as proven by drugs like metformin or the newly FDA-registered drug for MASH, resmetirom (Harrison et al., 2024; Lamoia & Shulman, 2021).

The data obtained using pharmacological and genetic approaches allowed us to reveal a new and unexpected role of CHIT1 in regulating cellular metabolism. Mechanistically, we observed a strong effect of CHIT1 inhibition on glycolysis in activated macrophages, which has broad therapeutic implications in disorders where chronic inflammation leads to tissue remodeling. Collectively, our data identify CHIT1 as a novel regulator of macrophage immunometabolism and underscore the beneficial effects of CHIT1 inhibition in MASH.

## STAR Methods

### Key resources

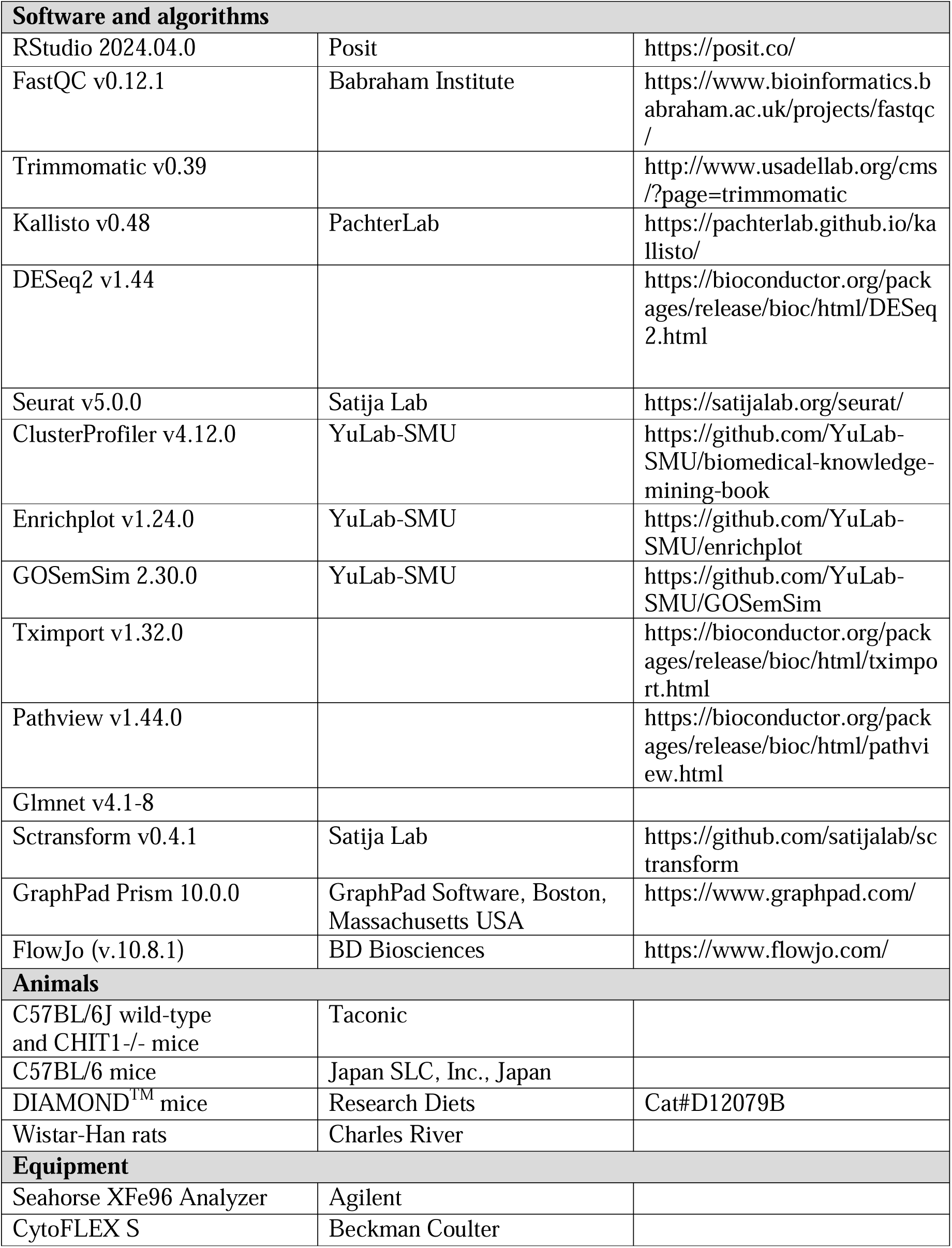
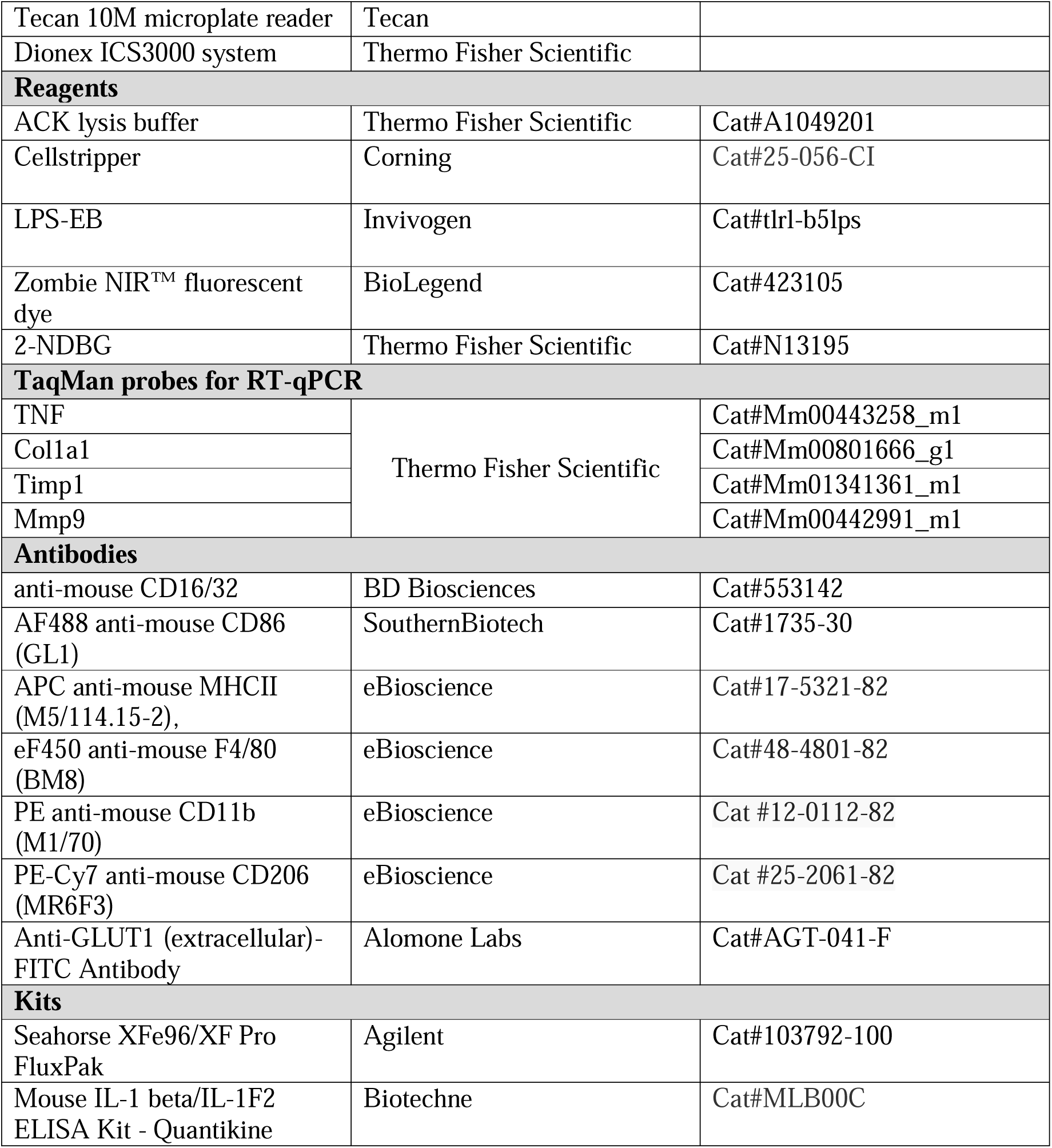

### Animal models

#### STAM

Efficacy study in STAM model (Stelic animal model) was performed at SMC Laboratories (Japan) with their standard protocols, as previously described (Fujii et al., 2013). Animals used in the study were cared for according to local guidelines: Act on Welfare and Management of Animals (Ministry of the Environment, Act No. 105 of October 1, 1973), Standards Relating to the Care and Management of Laboratory Animals and Relief of Pain (Notice No.88 of the Ministry of the Environment, April 28, 2006) and Guidelines for Proper Conduct of Animal Experiments (Science Council of Japan, June 1, 2006).

2-day-old C57BL/6 male pups from SPF females (Japan SLC, Inc., Japan) were subcutaneously injected with 200μg of streptozotocin (STZ, Sigma-Aldrich, USA). After 4 weeks animals were put on *ad libitum* high fat diet (HFD32, CLEA Japan, Inc., Japan). Control littermates (group “Control”) were fed with a standard diet (CE-2; CLEA-Japan Inc., Japan). At week 6, animals on HFD were randomized into two groups (n=10) – “MASH” and “OATD-01”. The latter group was administered orally once a day with 100 mg/kg OATD-01 in vehicle 0.5% (w/v in water) carboxymethylcellulose sodium salt (CMC, Nacalai Tesque, Inc., Japan) in dose-volume 10 ml/kg. The “Control” and “MASH” groups were treated the same way with vehicle. At 10 weeks of age animals were weighed and sacrificed, blood was collected for plasma, liver was dissected, weighed, and divided for various analyses.

Liver samples for histopathology and fibrosis evaluation were fixed in Bouin’s solution, processed to paraffin blocks, cut into 4 μm sections on rotary microtome (Leica Microsystems, Germany) and stained hematoxilin and eosin (HE) or picrosirious red (PSR).

MASLD Activity Score (MAS) was assessed blindly according to the criteria of Kleiner (Kleiner et al., 2005). Shortly, bright field images of HE-stained sections were randomly captured around the central vein using a digital camera (DFC295; Leica) at 50- and 200-fold magnification. MAS in 1 field/sample was evaluated. Steatosis score and other scores were calculated at 50- and 200-fold magnification, respectively.

Liver fragments for immunohistochemistry were embedded in OCT and cut on cryostat (Leica Microsystems, Germany) into 5 μm sections and fixed with acetone. Endogenous peroxidase were blocked by incubation with 0.03% H2O2, and unspecific epitopes with Block Ace (Dainippon Sumitomo Pharma Co. Ltd., Japan). The sections were incubated with anti-F4/80 antibody (BMA Biomedicals, Switzerland), followed by biotin-conjugated secondary antibody (VECTASTAIN Elite ABC kit, Vector laboratories) and ABC reagent. Enzyme-substrate reactions were performed using 3, 3’-diaminobenzidine/H2O2 solution (Nichirei Bioscience Inc.).

For quantitative analyses of fibrosis and inflammation areas, bright field images of Sirius red-stained and F4/80-immunostained sections were randomly captured around the central vein using a digital camera (DFC295; Leica) at 200-fold magnification, and the positive areas in 5 fields/section were measured using ImageJ software (National Institute of Health, USA).

#### DIAMOND model

Efficacy study in the DIAMOND™ mouse model was performed by Sanyal Biotechnology LLC (USA), with their standard protocols, as previously described (Asgharpour et al., 2016; Asgharpour & Sanyal, 2022). The study was conducted by all rules set forth by the International Association of Assessment and Accreditation of Laboratory Animal Care (iAAALAC) and the United States Office of Laboratory Animal Welfare (US OLAW) and had oversight from the Institutional Animal Care and Use Committee (IACUC, Protocol number: 19-009-OAT-01). Twenty DIAMOND^TM^ male mice were put on *ad libitum* western diet, WD (Research Diets, D12079B), and sweetened water (3.25% D-fructose and 2.25% D-glucose, SW). At week 7 mice were randomly assigned into groups: “MASH” and “OATD-01”. The latter group was administered orally once a day with 100 mg/kg OATD-01 in vehicle 0.5% (w/v in sweetened water) carboxymethylcellulose sodium salt (CMC, Nacalai Tesque, Inc., Japan) in dose-volume 20 ml/kg. 10 “Control” DIAMOND^TM^ male mice, fed *ad libitum* with a standard diet (Teklad Diets: LM-485/TD7012) and normal water, as well as “MASH” group, were administered with the vehicle in the same scheme as “OATD-01” group.

At the 24^th^ week animals were weighed and sacrificed, blood was collected for serum, liver was dissected, weighed, and divided for various analyses.

Serum transaminases activities were measured with VetScan VS2 Chemistry Analyzer (Abaxis/Zoetis), serum cholesterol and triglycerides levels were measured with Cholestech LDX blood lipid analyzers (Alere/Abbott). The chitinolytic activity was measured as described before (Sklepkiewicz et al., 2022).

For gene expression analysis, fragments of livers, preserved in RNAlater (Invitrogen) were homogenized and RNA was isolated using RNeasy MiniKit (Qiagen) according to the manufacturer’s protocol. Equal amount of RNA was used for RT PCR using High-Capacity cDNA Reverse Transcription Kit (Applied Biosystems). Gene-specific TaqMan Assays and TaqMan Gene Expression Master Mix (Applied Biosystems) were used for gene expression measurements. Real-time PCR Detection System CFX-384 (BIO-RAD) was used for PCR and relative gene expression was calculated based on 2-ΔΔCT method (Schmittgen & Livak, 2001). Primers used (Thermo Fisher Scientific Inc.): TNF: Mm00443258_m1; Col1a1: Mm00801666_g1; Timp1: Mm01341361_m1; Mmp9: Mm00442991_m1

For histopathology, fragments of livers were fixed in 10% neutral buffered formalin, processed to paraffin blocks, cut into 5 μm sections and stained with HE and Picrosirius red with standard protocols.

Steatosis, inflammation and hepatocyte ballooning, and subsequent NAS score, were analyzed blindly on HE-stained sections and fibrosis on PSR-stained section, according to the criteria of Kleiner (Kleiner et al., 2005).

#### Rat CDHFD model

Rat model was performed by Galapagos NV (Belgium), using their standard protocols and according to local regulations on ethics of animal use in research studies. Wistar-Han male rats from Charles River (France) were fed a control diet (A12450KR, Research Diet, New Brunswick, USA; “Control” group) or a choline-deficient, L-amino acid-defined, high-fat diet (CDHFD) (A16092003, Research Diet, New Brunswick, USA) for 12 weeks. At week 6 animals on CDHFD were randomly assigned to 3 groups: “MASH”, “ALK5i” and “OATD-01”.

“MASH” and “Control” animals were administered for 6 weeks with vehicle 0.5% (w/v) carboxymethyl cellulose, “ALKi” positive control animals were administered with the inhibitor of TGFb signaling at 1 mg/kg twice a day, and “OATD-01” was administered with OATD-01 at 100 mg/kg once a day, in volume 5 ml/kg.

At day 14th of treatment with OATD-01, in precise timepoints after formulation administration, blood samples were collected for compound concentration measurement and pharmacokinetic (PK) profiling.

At day 86 animals were sacrificed and samples of blood and fragments of livers were collected.

AST activity was measured in serum samples with Biochemical Strip Analyzer Spotchem. ELF (enhanced liver fibrosis) panel was analyzed by ELISAs (Rat TIMP-1 DuoSet® ELISA, DY580, lot P217394; Hyaluronan DuoSet® ELISA, DY3614-05, lot P296405; Rat N-Terminal Procollagen III propeptide (PIIINP), NOVUS, NBP2-76630, lot QLELV1THYM).

For histopathology, fragments of livers were fixed in 10% neutral buffered formalin, processed to paraffin blocks, cut into 5 μm sections and stained with immunohistochemistry to detect CD45 (Abcam, ab10558, lot GR3384334-1) and CD68 (Abcam, ab125212, lot GR77386-38). Analysis of IHC stained slides were performed by Visiopharm Image Analysis Software, Denmark)□

### Chitynolytic activity

Chitinolytic activity in rodent plasma and liver homogenates was measured using an established assay protocol (Sklepkiewicz et al., 2022). Shortly, 1 µl of plasma per well was mixed with 96 µM substrate - 4-methylumbelliferyl β-D-N,N’-diacetylchitobioside hydrate in assay buffer (0.1 M citrate, 0.2 M dibasic phosphate, 1 mg/ml BSA, pH 2 or pH 6) and incubated in a 96-well black microtiter plate with shaking in the dark, at 37°C for 60 minutes followed by addition of stop solution (0.3 M glycine/NaOH Buffer, pH 10.5). Substrate hydrolysis product - 4-methlyumberlliferone was measured fluorometrically using Tecan 10M microplate reader (excitation 355 nm/emission 460 nm). The chitinase activity was calculated using a standard curve of 4-methlyumberlliferone.

### Bioinformatics pipeline analyzing gene expression from bulk RNA-seq in Rat CDHFD model

Bulk RNAseq was performed by Galapagos NV (Belgium) upon liver sample collection from the Rat CDHFD model. Raw reads were processed by Trimmomatic software (Usadel, B. (2014)) to remove adaptor sequences and poor-quality reads. Transcript expression quantification was performed using kallisto v0.48 software (Bray et al., 2016) using Rnor_6.0 reference transcriptome as an index (GCA_000001895.4). Gene-level expression estimates were calculated using tximport R package (Soneson et al., 2015)

### RNA-seq Differential Expression

Before analysis, genes with no reads were filtered out of the dataset. The remaining genes were used to perform differential expression analysis using the DESeq2 package (v1.44.0), which uses the negative binomial generalized linear model to estimate logarithmic fold change along with the statistical significance of gene expression changes. The comparisons were performed between MASH vs Control groups and MASH vs MASH + OATD-01. P-values were adjusted by Benjamini-Hochberg correction. Genes without calculated p-values were filtered out for downstream analysis and log2(FoldChange) were shrunk according to the DESEq2 workflow. Counts were then normalized using Variance Stabilizing Transformation (VST).

### Gene Set Enrichment Analysis & Processes clustering

All retained genes were used for functional enrichment analysis done by GSEA algorithm implemented in ClusterProfiler package (v4.12.0) on Gene Ontology annotations retrieved from org.Rn.eg.db package (v3.19.1) and on KEGG pathways annotation downloaded online by ClusterProfiler package. Ranked gene lists by their log2(Fold Change) were used as an input. The analysis was run with default parameters, except p-adjust method, which was set to Bonferroni correction. Due to large numbers statistically, significant biological processes returned by GSEA, there was performed semantic similarity analysis to group processes that share similarities in terms of genes involved in particular processes. For this purpose, Gene Ontology annotations were obtained by GOSemSim package (v2.30.0), and pairwise similarities were calculated using the Jiang method performed by enrichplot package (v1.24.0). Then pairwise similarities of the enriched terms were clustered using a hierarchical clustering approach. The obtained clusters were then named based on the processes assigned to each cluster, and the genes belonging to them were merged. The clusters were analyzed for gene similarity within each process. Some clusters contained several subclusters, which were analyzed separately (see Supplementary material on hierarchical clustering). To get information about the NES coefficient for each cluster, the median NES from the processes comprising the cluster was calculated.

### Mice MASLD single-cell RNA-seq & deconvolution

To explore liver cell type heterogeneity and composition, we analyzed publicly available single-cell and single-nucleus RNA-seq data derived from mouse livers (GSE192742). Mice were divided into groups fed a standard diet and a Western diet inducing MASH, both for 24 weeks. Counts were processed using the Seurat v5 (Satija et al., 2015) standard workflow. As the cells were derived from different sources of digestion (ex-vivo and nuclei), counts were batch-corrected to eliminate this source of variation. We employed the Variance Stabilizing Transformation approach implemented in sctransform (Satija et al., 2022) to remove unwanted effects and correct counts. These corrected counts were used for further analysis, including normalization, variable feature selection, clustering, and integration. For the deconvolution analysis, single-cell and single-nucleus RNA-seq expression data were aggregated to obtain summarized expression profiles of each gene per cell type. Deconvolution analysis was performed for each group of rats (Control, MASH, OATD-01) using a Ridge regression model performed by glmnet package (Hastie T. et al. 2010). Negative coefficients were zeroed, and proportions were calculated from the remaining regression coefficients.

### Generation, Polarization, and Culture Conditions of BMDMs

Cells were cultured in a complete experimental medium continuously from isolation to the endpoint. The complete experimental medium consists of DMEM (Dulbecco’s Modified Eagle Medium), 10% heat-inactivated FBS, 1% penicillin/streptomycin, 50 ng/ml of colony-stimulating factor (mM-CSF), and OATD-01 at concentrations of 1 μM and 10 μM, or DMSO, respectively. DMSO was used as the vehicle control.

Bone marrow was retrieved from the femurs and tibias of 6- to 10-week-old male C57BL/6J wild-type and CHIT1-/- mice purchased from Taconic. After isolation, the bone marrow was filtered through a cell strainer (70 µm), washed with PBS, and subjected to red blood cell lysis using ACK lysis buffer (Thermo Fisher, A1049201), followed by neutralization in culture medium. The obtained bone marrow cells were counted and seeded at a density of 1 × 10^6 cells/ml on a petri dish in a final volume of 10 ml. On days 3 and 5, 10 ml and 5 ml of medium were added, respectively. After 6 days, cells were considered macrophages, detached from vessels using Cellstripper (Corning), counted, and reseeded for downstream applications. After reseeding, when the cells were attached, they were stimulated with 100 ng/ml of LPS (LPS-EB, Invivogen) for M1-like (M(LPS)) subtype generation.

For the Seahorse assay, cells were plated at a density of 1 × 10^5 cells/well in 100 μl of complete experimental medium and grown in Seahorse XF96 polystyrene tissue culture plates (Agilent).

For flow cytometry-based analysis of Glut1 and glucose uptake, cells were plated at a density of 0.8 × 10^6 cells/well in 2 ml of experimental medium in a 6-well plate format.

For metabolite analysis, cells were seeded at a density of 1 × 10^6 cells/ml on a petri dish in a final volume of 10 ml.

### Seahorse Assay

Before measurement, cells were incubated in Seahorse XF DMEM assay medium (without phenol red and sodium bicarbonate) containing 10 mM glucose, 1 mM pyruvate, and 2 mM glutamine in a non-CO2 incubator at 37□°C for 1□hour. The oxygen consumption rate (OCR) was measured every 3 minutes with a mixing time of 3 minutes in each cycle, with 3 cycles per step using the Seahorse XFe96 Analyzer (Agilent). The Cell Mito Stress Test was used to assess mitochondrial function. The sequential addition of oligomycin A (1.0□µM), FCCP (1.5□µM), and antimycin A/rotenone (0.5□µM) dissolved in DMEM assay medium allowed for the calculation of OCR linked to ATP production, maximal respiration capacity, and spare respiratory capacity. Basal respiration was measured before the injection of oligomycin A. The simultaneous measurements of the extracellular acidification rate (ECAR) allowed for the calculation of basal glycolysis and compensatory glycolysis. Finally, the data were normalized according to the total protein content in each well.

### Flow cytometry based analysis

#### Phenotyping of BMDMs

Flow cytometry analysis was applied to characterize cell types. Briefly, after collecting and washing in PBS, cells were subjected to staining with specific antibodies. First, cells were stained for viability using Zombie NIR™ fluorescent dye (BioLegend, #423105). After washing in PBS, Fc receptors on the cell surface were blocked by incubation in purified anti-mouse CD16/32 (BD Biosciences, #553142) diluted in FACS buffer (PBS containing 2% FBS, 0.5 mM EDTA). Then, to quantify the expression of macrophage cell surface receptors, cells were incubated with a combination of the following anti-mouse conjugated antibodies from SouthernBiotech:AF488 anti-mouse CD86 (GL1) and eBioscience: APC anti-mouse MHCII (M5/114.15-2), eF450 anti-mouse F4/80 (BM8), PE anti-mouse CD11b (M1/70), and PE-Cy7 anti-mouse CD206 (MR6F3). After washing, cells were resuspended in FACS buffer and the fluorescent intensity of each antibody was measured.

#### Glut1 Cell Surface Protein Expression

For Glut1 cell surface protein expression, an antibody designed for detection of the extracellular domain was used (Alomone Labs, #AGT-041-F). After stimulation with LPS for 48 hours, BMDMs were washed with PBS, detached as previously described, stained with Zombie NIR, followed by blocking with FcBlock and labeling with Glut-FITC conjugated antibody. Flow cytometry was performed using the CytoFLEX S (Beckman Coulter) system. Data were analyzed using FlowJo (v.10.8.1).

### Glucose Uptake

To study cellular metabolism and the rate at which cells absorb glucose, a glucose uptake assay using the fluorescent glucose analog 2-NDBG (Thermo Fisher, #N13195) and the protocol described by (Palmer et al., 2016), was employed. Briefly, at 48 hours of LPS stimulation, cells were washed with PBS, detached from vessels as previously described, and incubated with 2-NBDG at a final concentration of 1.46 µM for 30 minutes.

### Metabolites analysis

#### Preparation of samples for IC-MS analysis

Cells were washed with warm (37°C) 0.3 M mannitol, ensuring that all mannitol was removed after the wash. Then a solution of ice-cold methanol:acetonitrile (2:1:1, v/v/v) was used for extraction as described in (Bennett et al., 2008). Following extraction, samples were centrifuged (5 minutes, 4°C, 13,000 rpm) and the precipitates were resuspended in 100 mM NaOH + 0.125% Triton X-100 for protein quantification using the Bradford method. The supernatants were stored at -80°C until analysis.

#### IC-MS

Anionic metabolites were separated by ion chromatography using a modification of the method described in (Bhattacharya et al., 1995). The separation employed a Dionex ICS3000 system equipped with a KOH eluent generator and a Dionex ADRS 600 suppressor. Two connected Dionex IonPac AS11-HC columns were used to achieve separation with a KOH gradient. Chromatograms for ATP, ADP, and AMP were obtained by monitoring eluent absorbance at 260 nm using a UV detector placed before the suppressor on the system. All other metabolites were detected using a Waters ZQ mass spectrometer coupled to the IC system. The electrospray source was utilized, and data acquisition was performed for negative ions. Potential differences in ionization efficiency due to matrix and contamination effects were addressed by spiking the samples with tartrate as an internal standard before IC separation. All metabolite quantifications were performed relative to this internal standard (Schaub et al., 2006; W. Wang et al., 2003).

### ELISA for IL-1β

ELISA for IL-1β was performed according to the manufacturer’s protocol (Cat#MLB00C; Biotechne) using frozen cell supernatants collected from BMDMs, which were either stimulated with LPS for 48 hours or left unstimulated.

### Statistical analysis

Data were analyzed using GraphPad Prism v. 10.0.0 (GraphPad Software, Boston, Massachusetts, USA). To evaluate differences between groups for in vivo read-outs, an ordinary one-way ANOVA with Holm-Šídák correction for multiple comparisons was applied in cases of normal distribution, while the Kruskal-Wallis test with Dunn’s correction for multiple comparisons was applied for non-normal distributions. For BMDM experiments, either an unpaired t-test or parametric one-way ANOVA with Fisher’s test for multiple comparisons was used to evaluate differences between groups. P-values < 0.05 were considered significant and noted with asterisks (* for P < 0.05, ** for P < 0.01, *** for P < 0.001, **** for P < 0.0001).

## Supporting information

Supplementary material of hierarchical clustering

## Supplementary Figure Legends

**Supplementary Figure 1. OATD-01 restores cellular composition affected in MASH condition from the MASH rat study.** Deconvolution of RNAseq dissecting proportions of different populations of cells in livers in control, MASH, and OATD-01-treated rats.

**Supplementary Figure 2. OATD-01 reverses the expression of a significant number of genes implicated in MASH pathogenesis.** Venn Diagrams of selected clustered GO processes indicated by GESEA analysis of transcriptomic data from control, MASH, and MASH + OATD-01 groups in the MASH rat study.

**Supplementary Figure 3. OATD-01 regulates cholesterol flux by reversing changes in gene expression observed in MASH condition from the rat study.** The cholesterol metabolism pathway from the KEGG database, with indicated genes found in the transcriptomic analysis, shows changes in gene expression for MASH vs Control and MASH vs MASH + OATD-01. Gene expression changes are presented using log2(FoldChange) and visualized through a color gradient, where blue indicates strong downregulation and red indicates strong upregulation.

**Supplementary Figure 4. Glycolysis is altered in MASH and regulated by OATD-01.** Heatmaps displaying the expression of genes involved in glycolysis and TCA cycle from MASH rat study. Genes were fetched from the KEGG database. Gene expression changes are presented using log2(FoldChange) and visualized through a color gradient where blue indicates strong downregulation, and red indicates strong upregulation.

**Supplementary Figure 5. OATD-01 and CHIT KO do not affect the phenotype of BMDMs stimulated with LPS**. FACs analysis of LPS-treated BMDMs from WT and KO mice.

**Supplementary Figure 6. OATD-01 and CHIT1 KO do not affect levels of GLUT1 transporter on the surface of macrophages.** FACs analysis of GLUT1 extracellular domain in LPS-treated BMDMs for 48h from WT and KO mice. Data presented are shown as means and standard deviation. The statistical test is an unpaired t-test. Statistical significance between groups is indicated as follows: * p < 0.05, ** p < 0.01, *** p < 0.001, **** p < 0.0001.

## Supplementary information

▪ Table S1. Excel file containing the list of gene ontology terms found in the transcriptomic analysis of the rat MASH study.
▪ Table S2. Excel file containing the list of KEGG pathways found in the transcriptomic analysis of the rat MASH study
▪ Supplementary material on hierarchical clustering. The PDF file with the graphic explanation of the clustering process related to Fig. 5
▪ Figures S1-S6

## Author contributions

K.D. and Z.Z designed the experiments, interpreted the data, and wrote the paper; K.G. conducted the experiments on BMDMs; M.M. designed in vivo experiments; B.H. analyzed RNA-seq data; D.R designed RNA-seq analysis experiments; A.M. designed and supervised BMDM experiments; I.T. and B.W supervised RNA-seq data.; K.P. performed ELISA experiments; D.D. and K.Z. performed Seahorse analysis; T. Ryan designed metabolic experiments; A.J. performed metabolic measurements; L.A.J.O. revised the project and gave intellectual assistance; B.D., K.L, T.Rejczak., and A.G. developed OATD-01 drug for this study.

## Acknowledgments

We thank our funding organizations for supporting this research through three projects: (1) “Preclinical research and clinical trials of a first-in-class development candidate in the therapy of asthma and inflammatory bowel disease” (POIR.01.01.01-00-0168/15), acronym IBD, (2) “Development of a ‘first-in-class’ small molecule drug candidate for treatment of idiopathic pulmonary fibrosis through chitotriosidase inhibition” (POIR.01.01.01-00-0551/15), acronym IPF, both co-financed by European Union through the European Regional Development Fund within the Smart Growth Operational Programme and (3) “Preclinical and clinical development of drug candidate OATD-01, for the treatment of sarcoidosis patients” (MAZOWSZE/0128/19), acronym SARCO, as part of the “Path for Mazovia” competition co-financed by the National Centre for Research and Development from national funds. B.D. was supported by the Ministry of Science and Higher Education, Poland (50// DW/2017/01/1).

