## Supplementary Figure 1-6 for "Chitinase-1 inhibition reverses metabolic dysregulation and restores homeostasis in MASH animal models"

**Supplementary Figures**

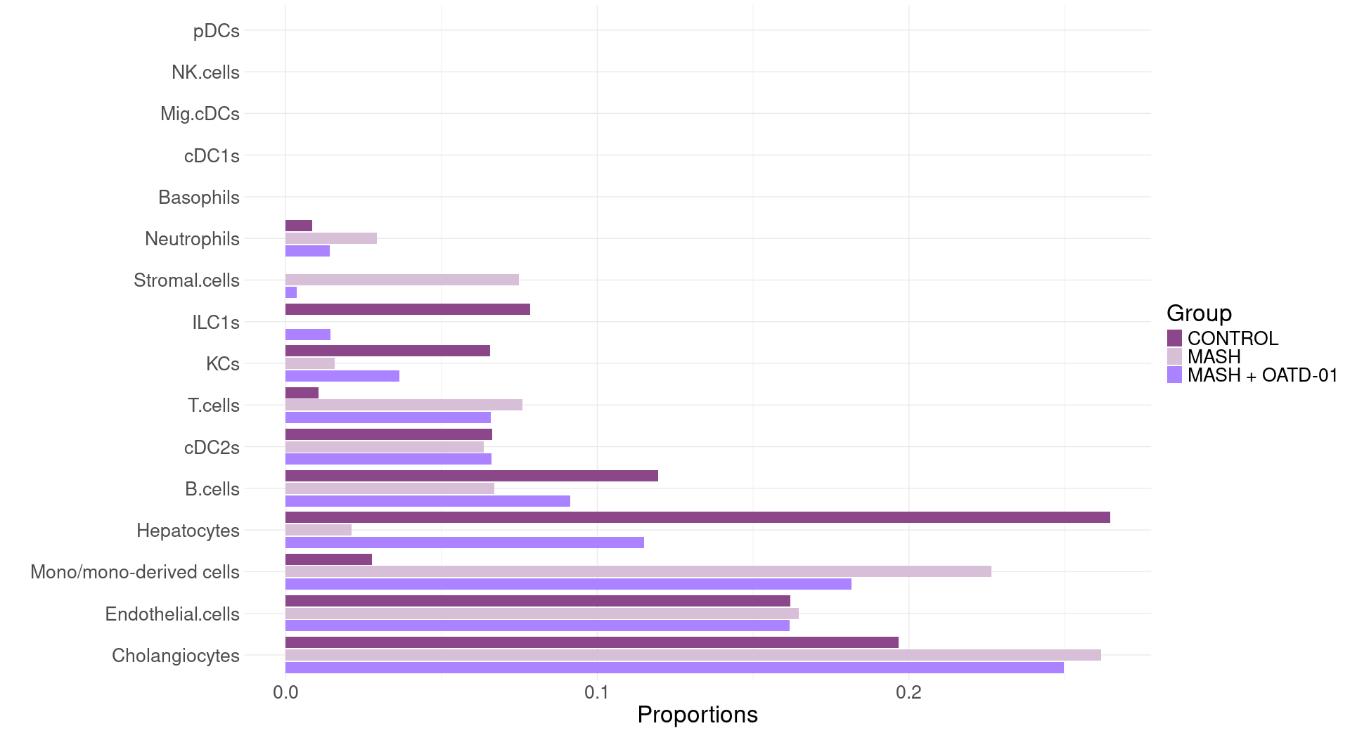

**Supplementary Figure 1. OATD-01 restores cellular composition affected in MASH condition from the MASH rat study.** Deconvolution of RNAseq dissecting proportions of different populations of cells in livers in control, MASH, and OATD-01-treated rats.

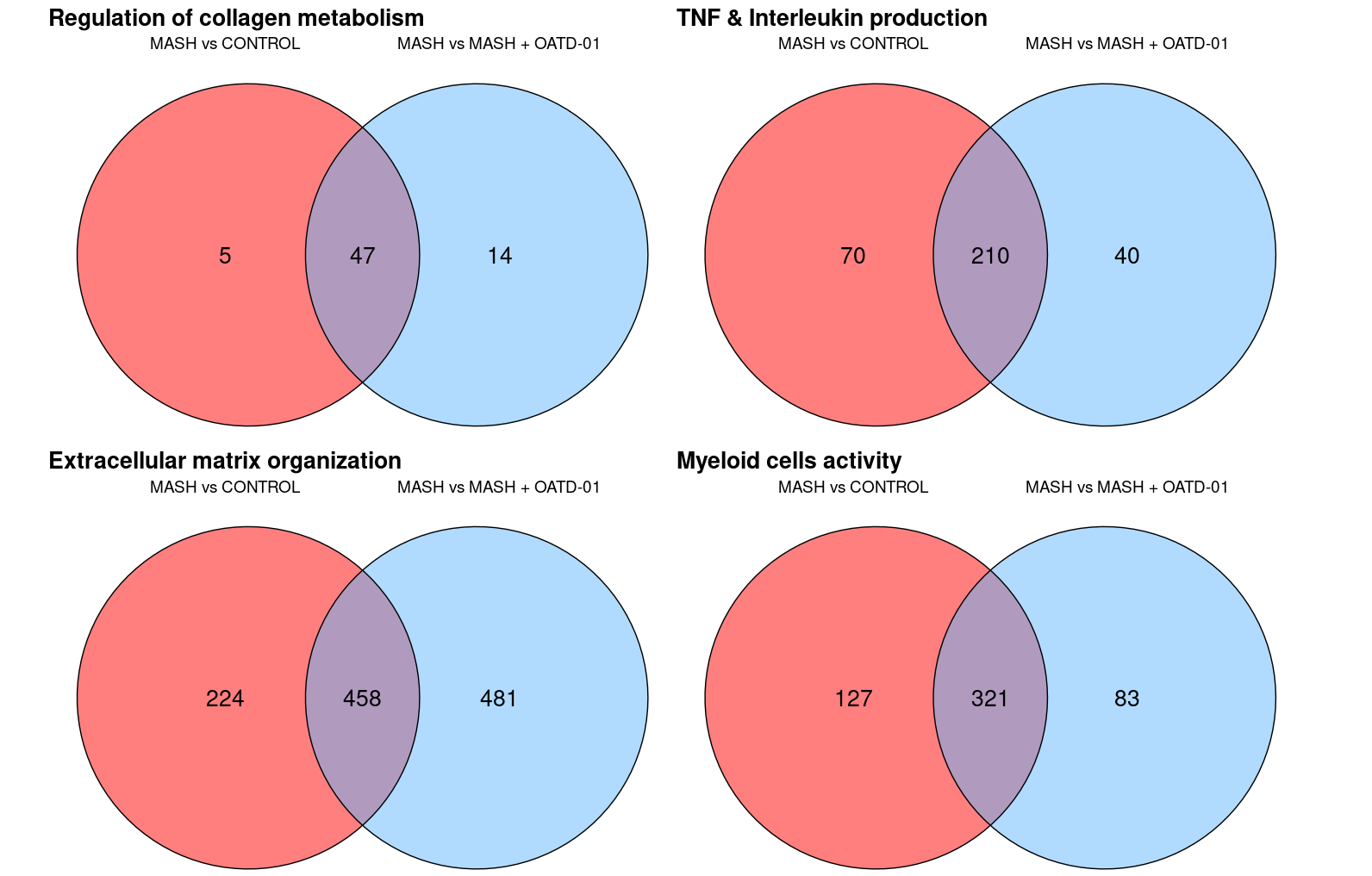

**Supplementary Figure 2. OATD-01 reverses the expression of a significant number of genes implicated in MASH pathogenesis.** Venn Diagrams of selected clustered GO processes indicated by GESEA analysis of transcriptomic data from control, MASH, and MASH + OATD-01 groups in the MASH rat study.

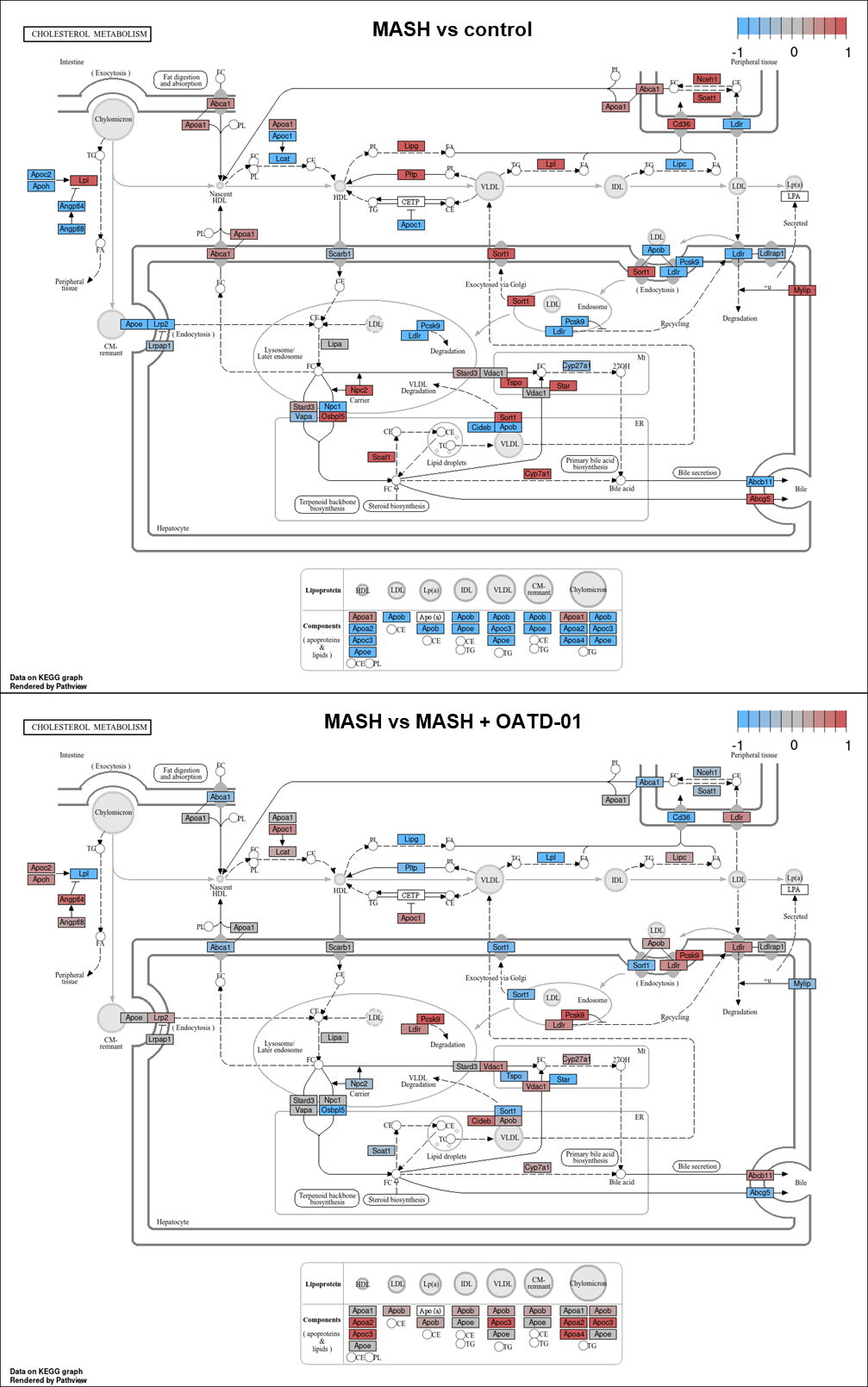

**Supplementary Figure 3. OATD-01 regulates cholesterol flux by reversing changes in gene expression observed in MASH condition from the rat study.** The cholesterol metabolism pathway from the KEGG database, with indicated genes found in the transcriptomic analysis, shows changes in gene expression for MASH vs Control and MASH vs MASH + OATD-01. Gene expression changes are presented using log2(FoldChange) and visualized through a color gradient, where blue indicates strong downregulation and red indicates strong upregulation.

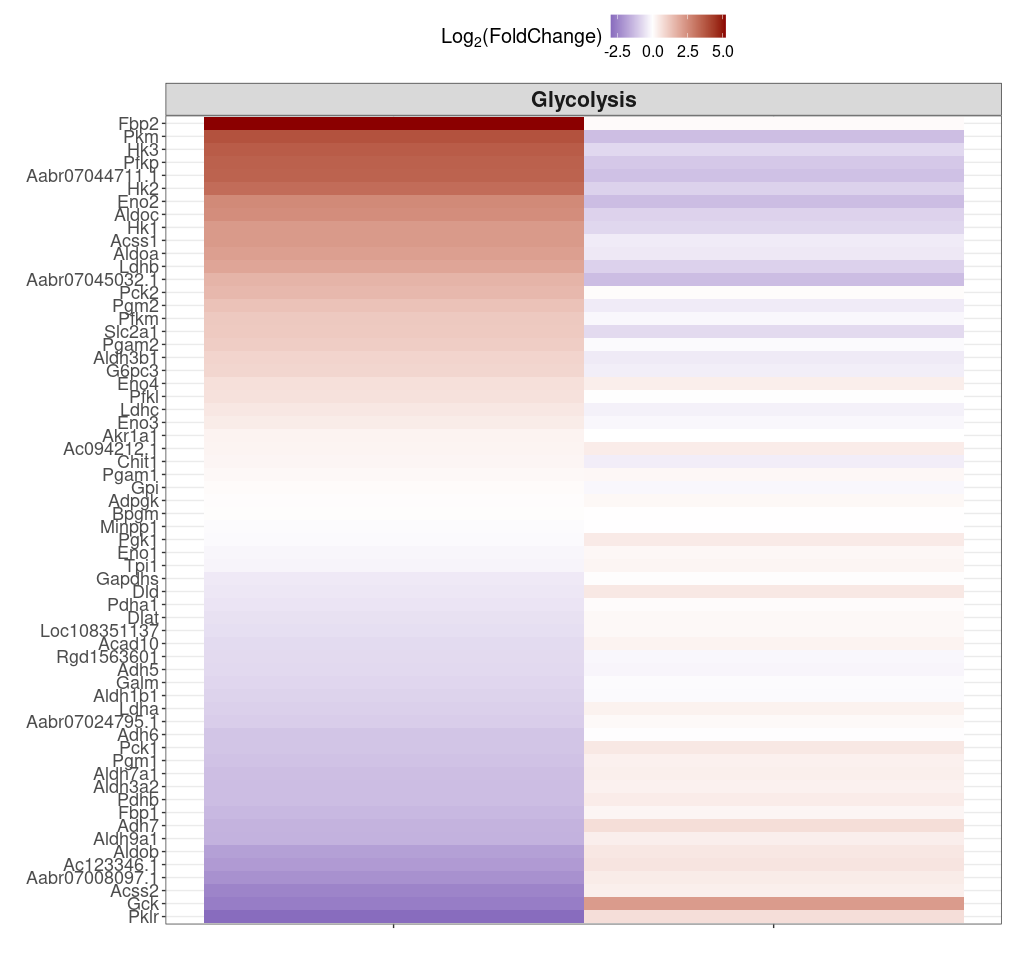
**Supplementary Figure 4. Glycolysis is altered in MASH and regulated by OATD-01.** Heatmaps displaying the expression of genes involved in glycolysis and TCA cycle from MASH rat study. Genes were fetched from the KEGG database. Gene expression changes are presented using log2(FoldChange) and visualized through a color gradient where blue indicates strong downregulation, and red indicates strong upregulation.

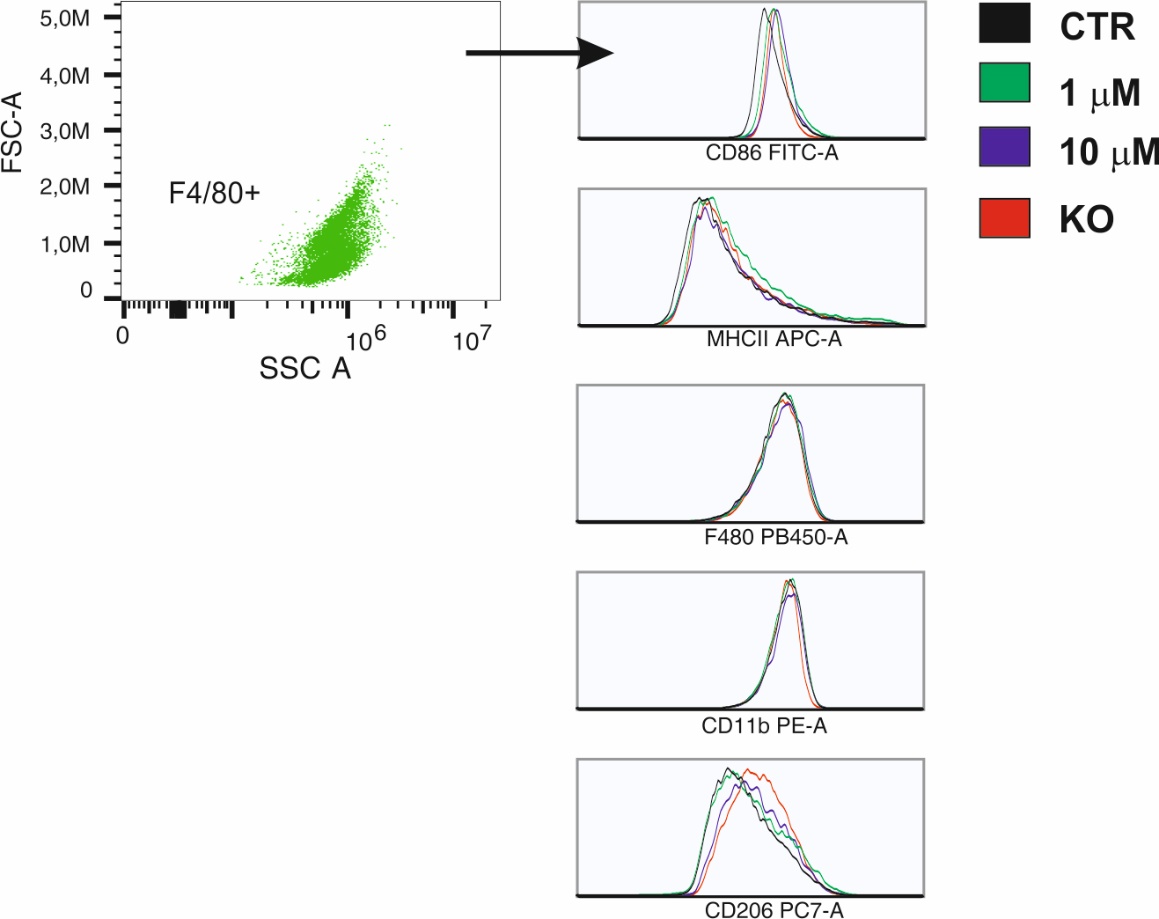

**Supplementary Figure 5. OATD-01 and CHIT KO do not affect the phenotype of BMDMs stimulated with LPS**. FACs analysis of LPS-treated BMDMs from WT and KO mice.

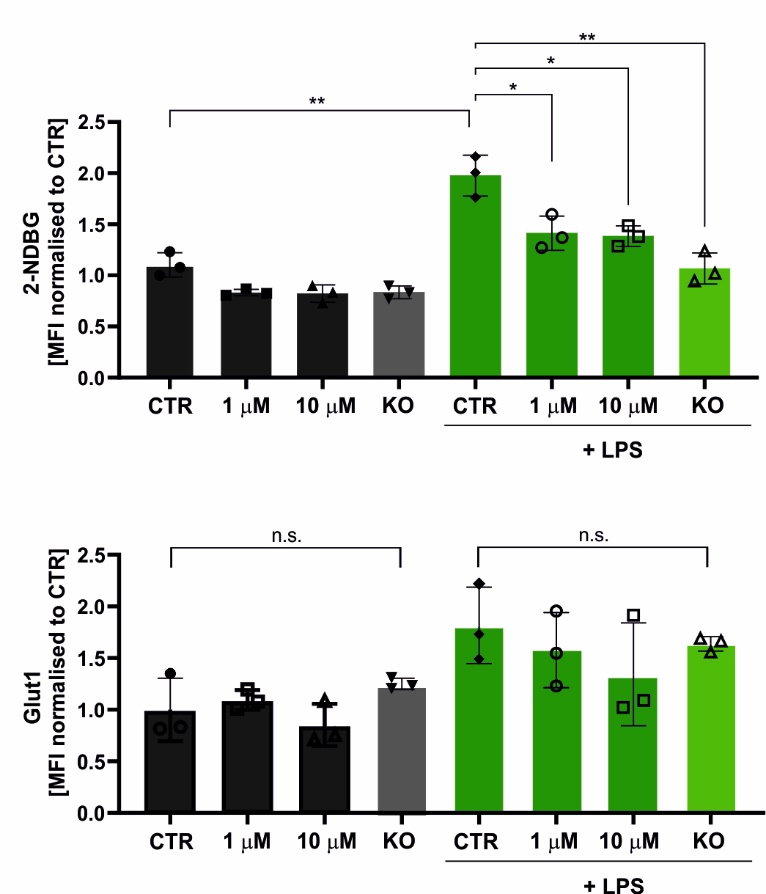

**Supplementary Figure 6. OATD-01 and CHIT1 KO do not affect levels of GLUT1 transporter on the surface of macrophages.** FACs analysis of GLUT1 extracellular domain in LPS-treated BMDMs for 48h from WT and KO mice. Data presented are shown as means and standard deviation. The statistical test is an unpaired t-test. Statistical significance between groups is indicated as follows: * p < 0.05, ** p < 0.01, *** p < 0.001, **** p < 0.0001.
