## Supplementary material of hierarchical clustering for "Chitinase-1 inhibition reverses metabolic dysregulation and restores homeostasis in MASH animal models"

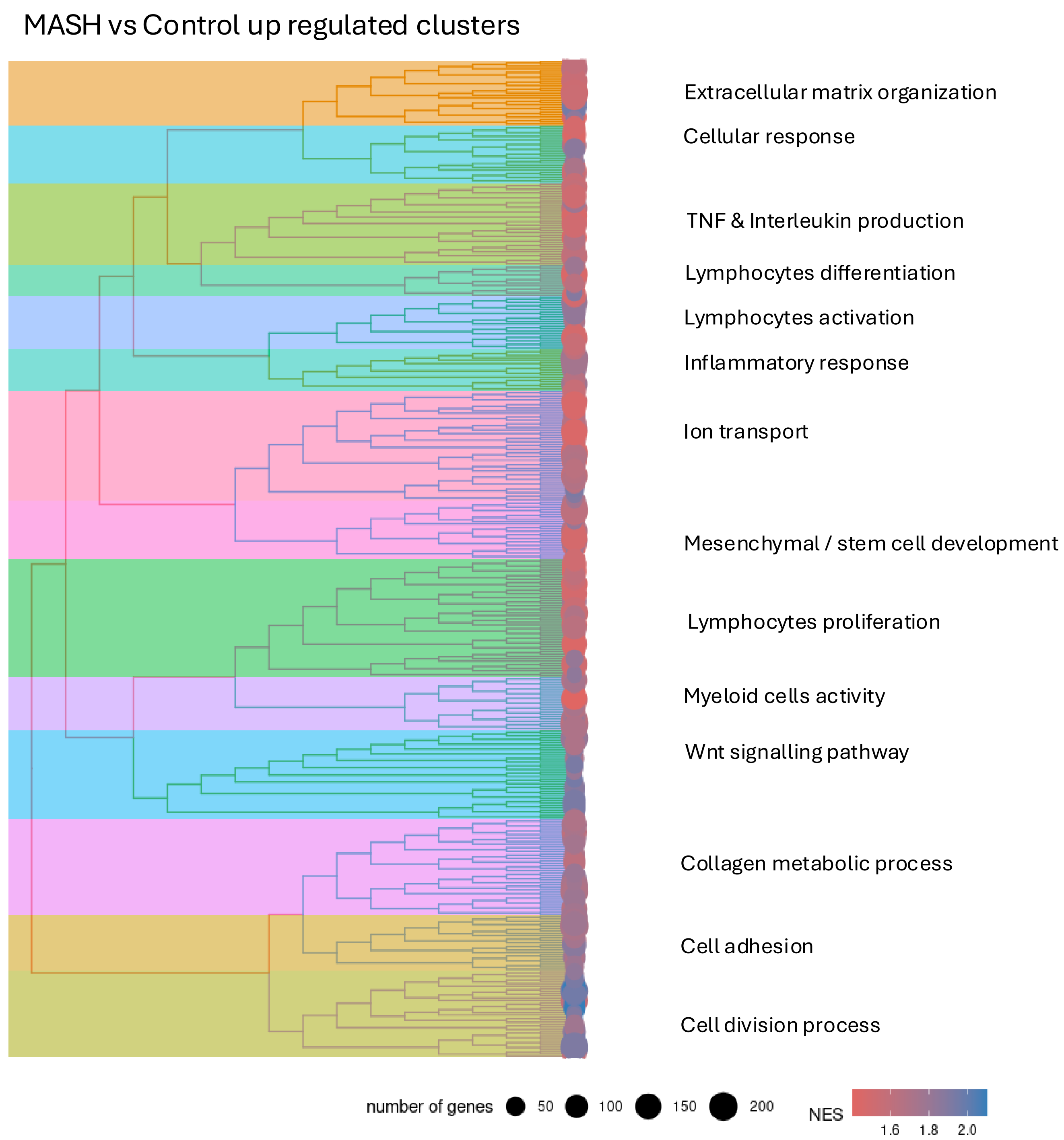
MASH vs Control up regulated clusters
Extracellular matrix organization
Cellular response
TNF & Interleukin production
Lymphocytes differentiation
Lymphocytes activation
Inflammatory response
Ion transport
Mesenchymal / stem cell development
Lymphocytes proliferation
Myeloid cells activity
Wnt signalling pathway
Collagen metabolic process
Cell adhesion
Cell division process
number of genes
50
100
150
200
NES
1.6
1.8
2.0

MASH vs Control down regulated clusters

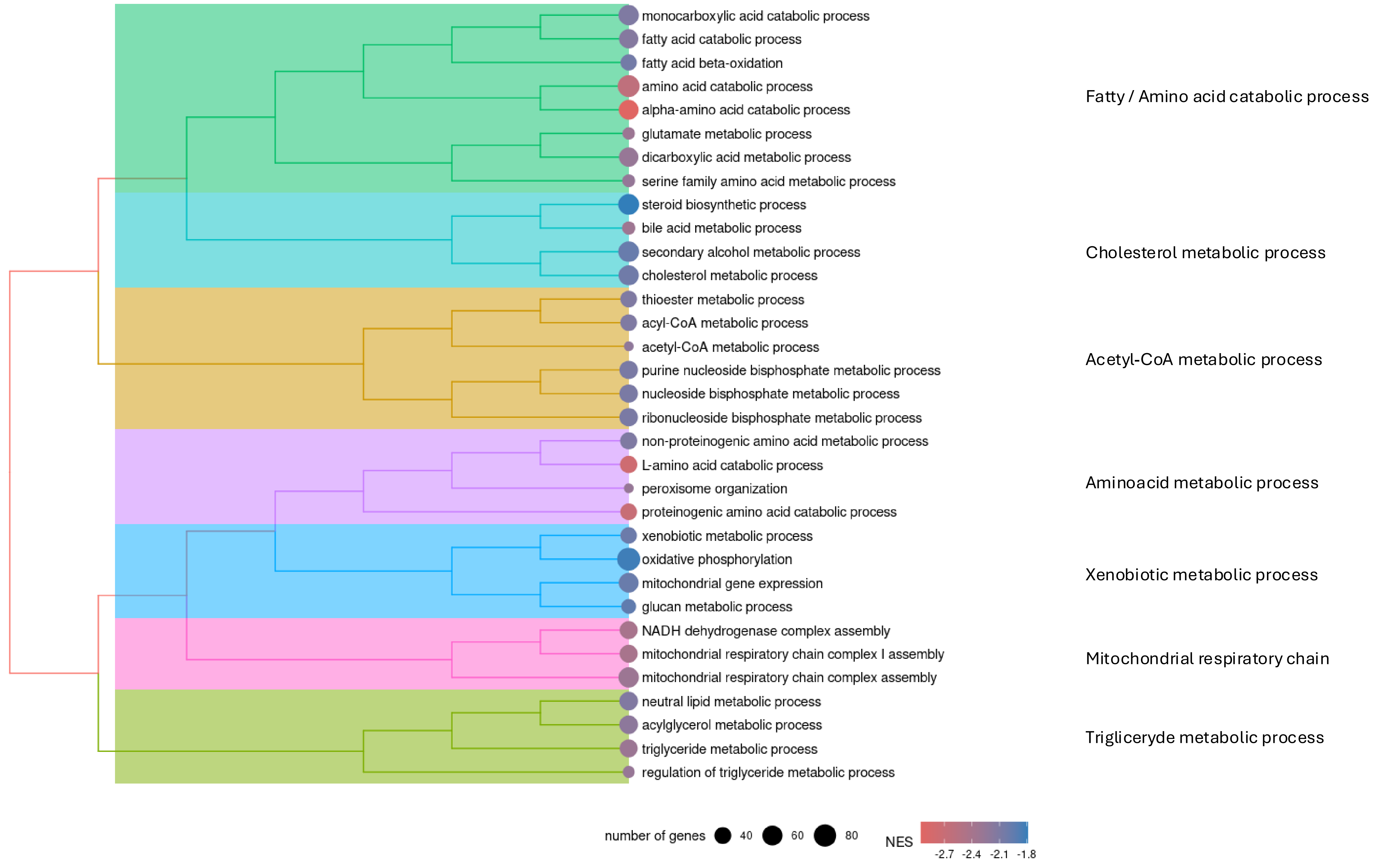

MASH vs MASH + OATD-01 up regulated clusters

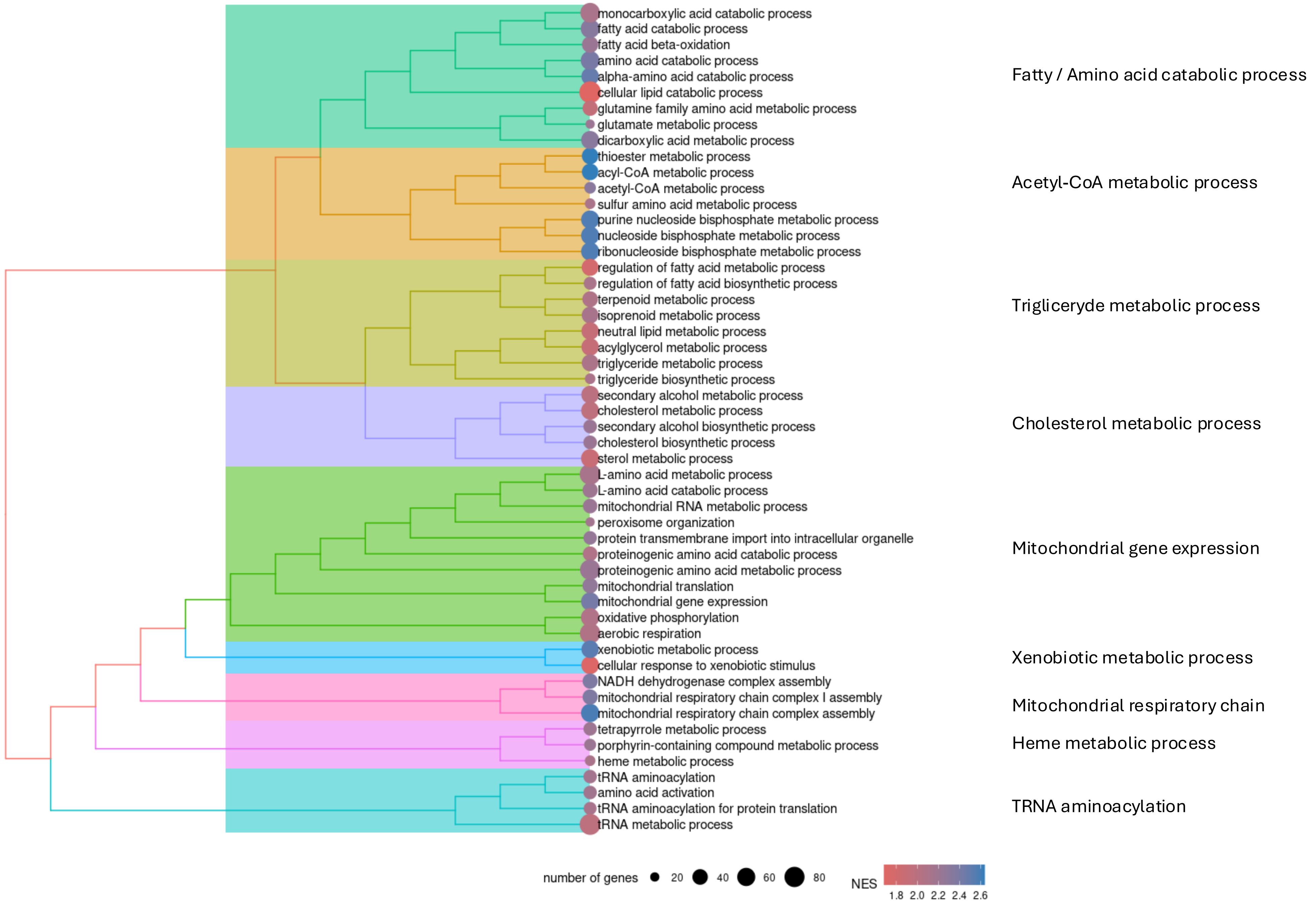

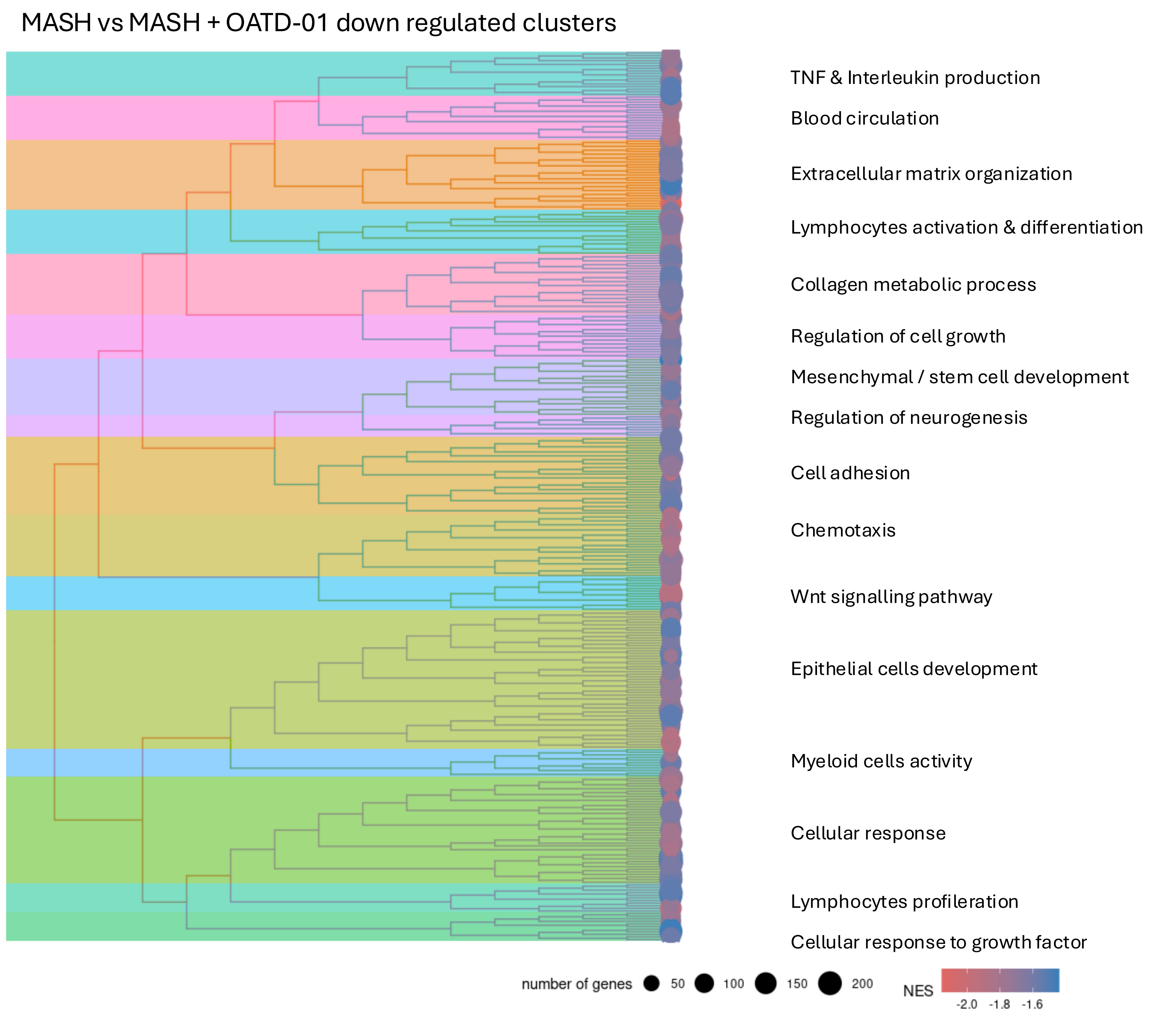
MASH vs MASH + OATD-01 down regulated clusters
TNF & Interleukin production
Blood circulation
Extracellular matrix organization
Lymphocytes activation & differentiation
Collagen metabolic process
Regulation of cell growth
Mesenchymal / stem cell development
Regulation of neurogenesis
Cell adhesion
Chemotaxis
Wnt signalling pathway
Epithelial cells development
Myeloid cells activity
Cellular response
Lymphocytes profileration
Cellular response to growth factor
number of genes
50
100
150
200
NES
-2.0
-1.8
-1.6

### MASH vs Control - Extracellular matrix organization cluster

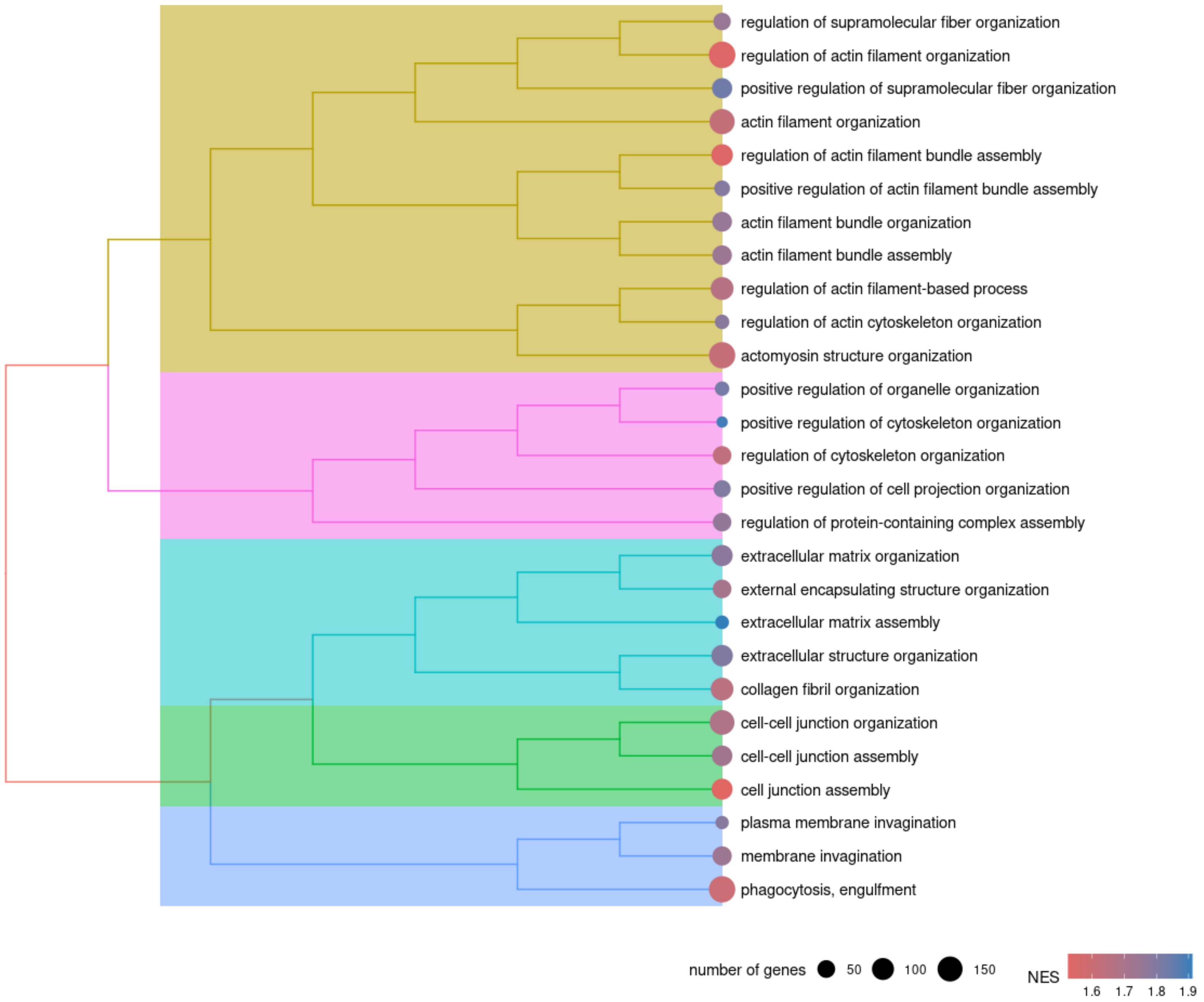

### MASH vs Control Inflammatory response cluster

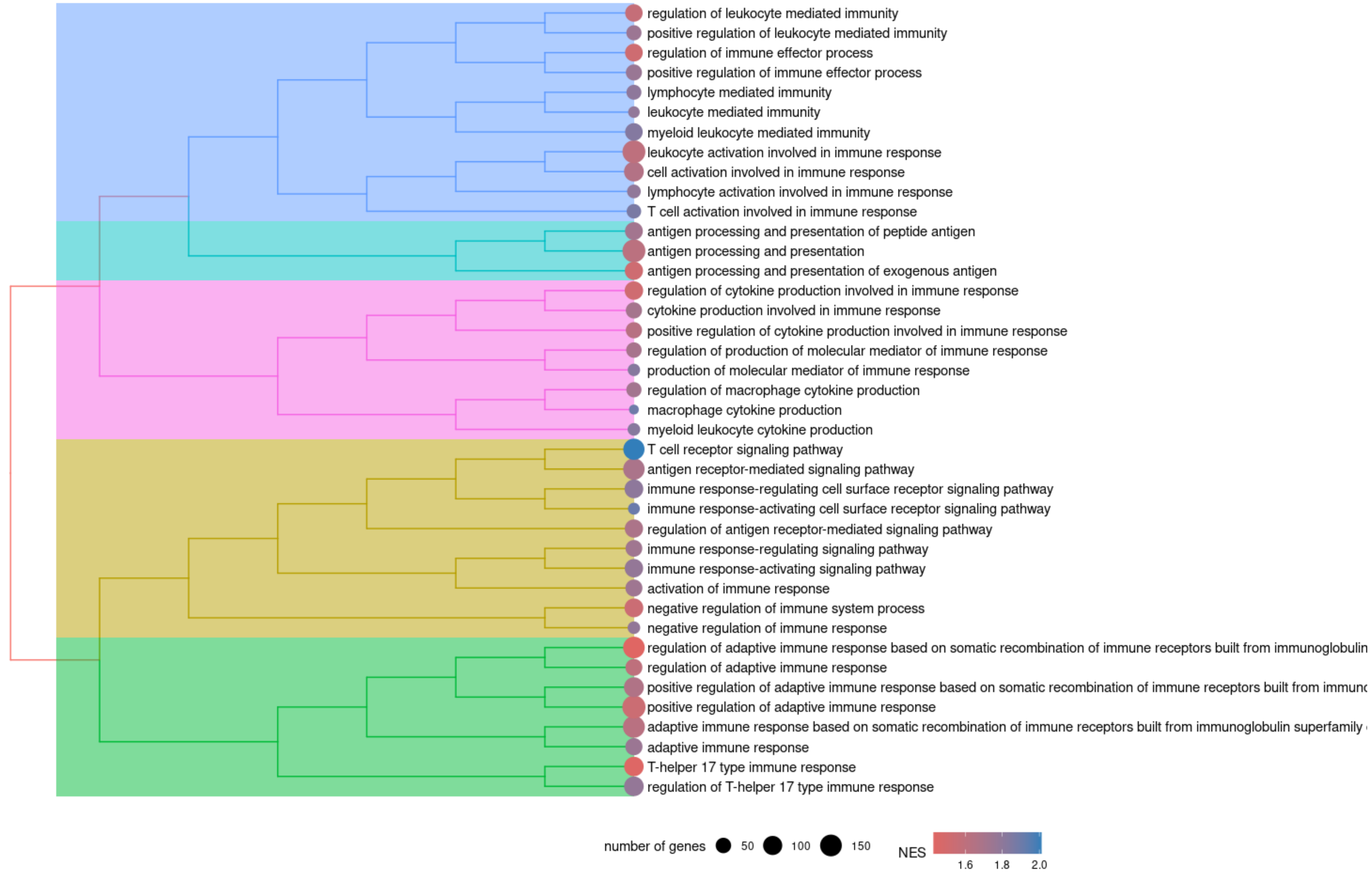

### MASH vs Control Cellular response - cluster

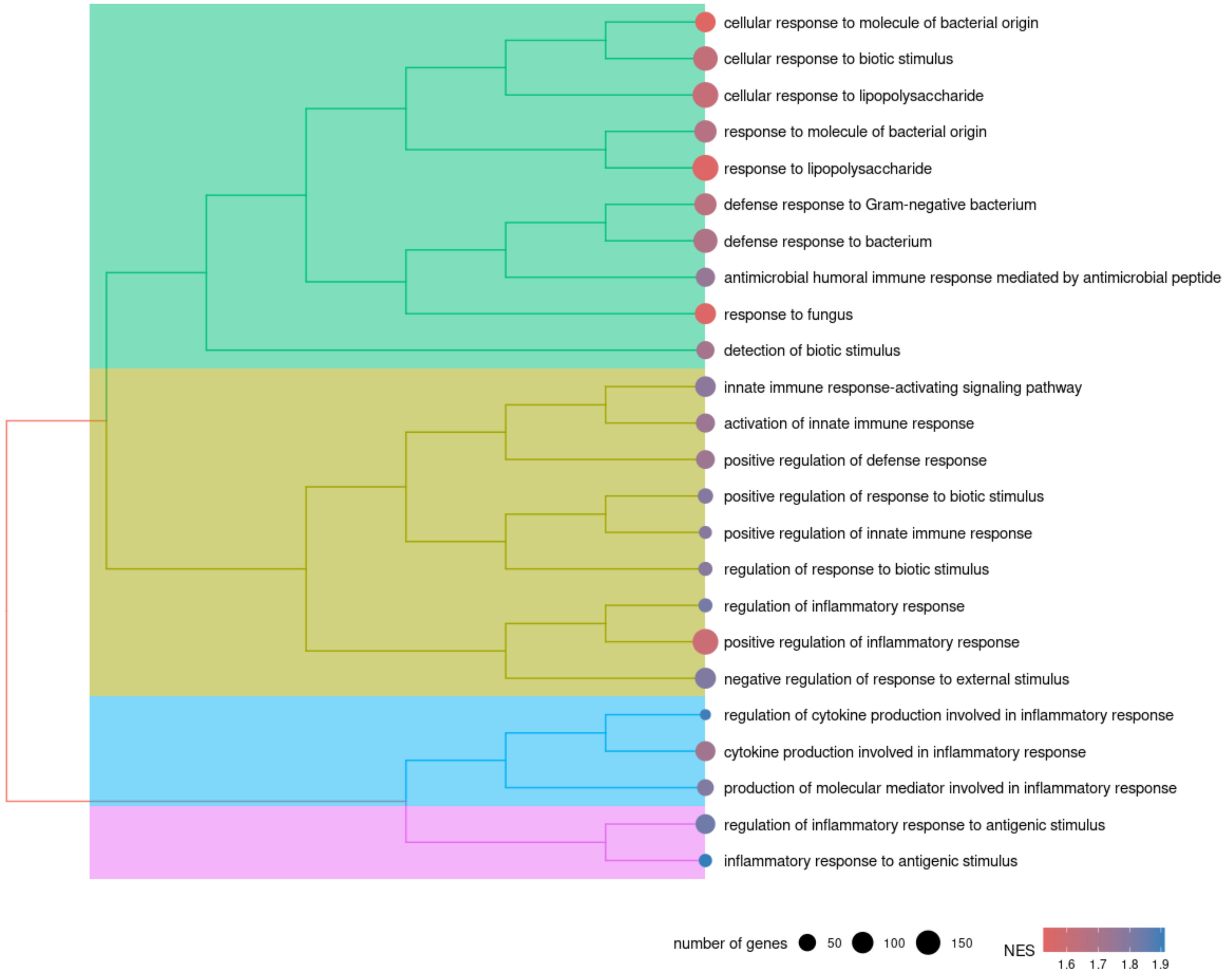

### MASH vs Control - Wnt signalling pathway cluster

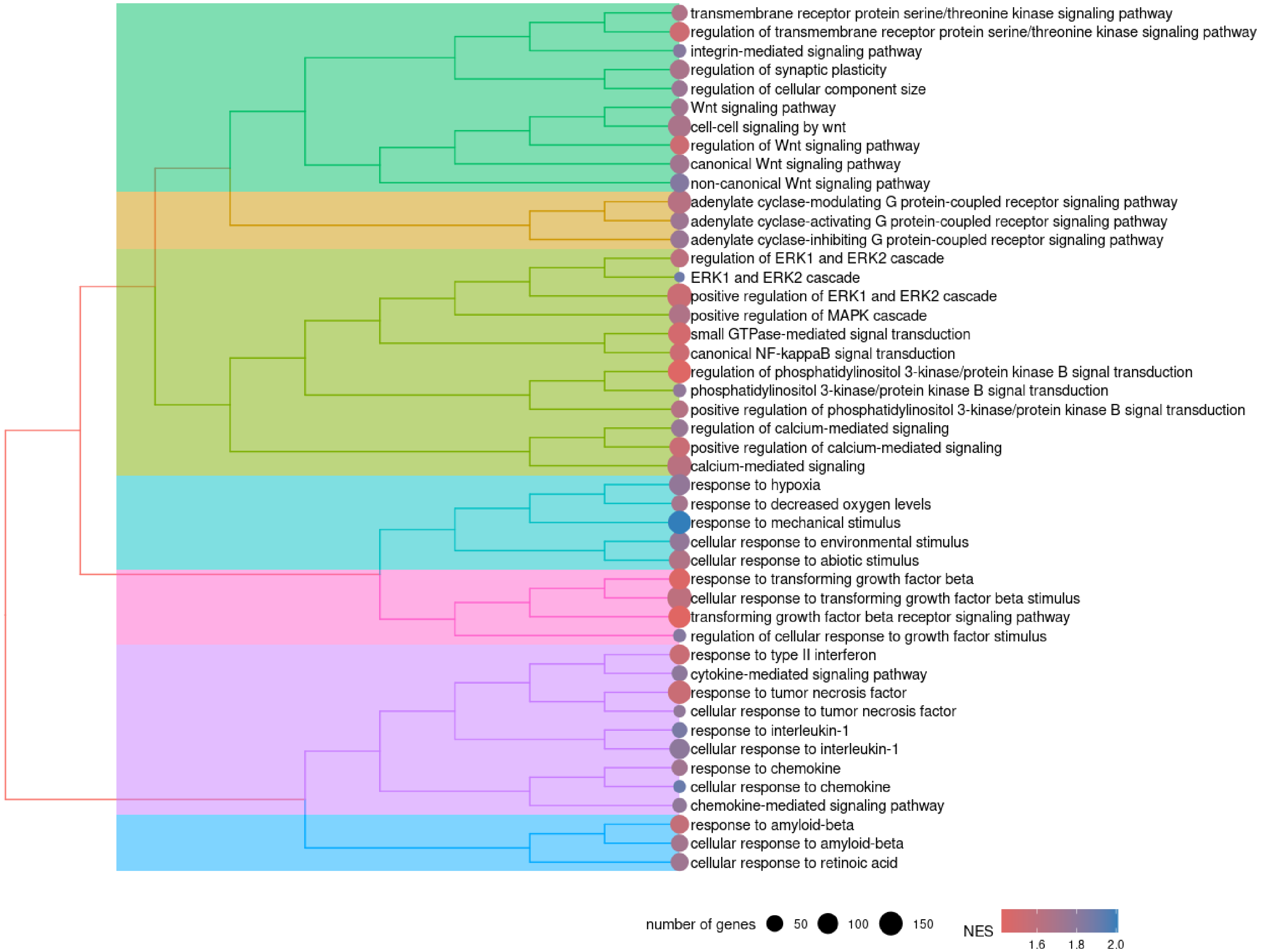

### MASH vs Control - Lymphocytes activation cluster

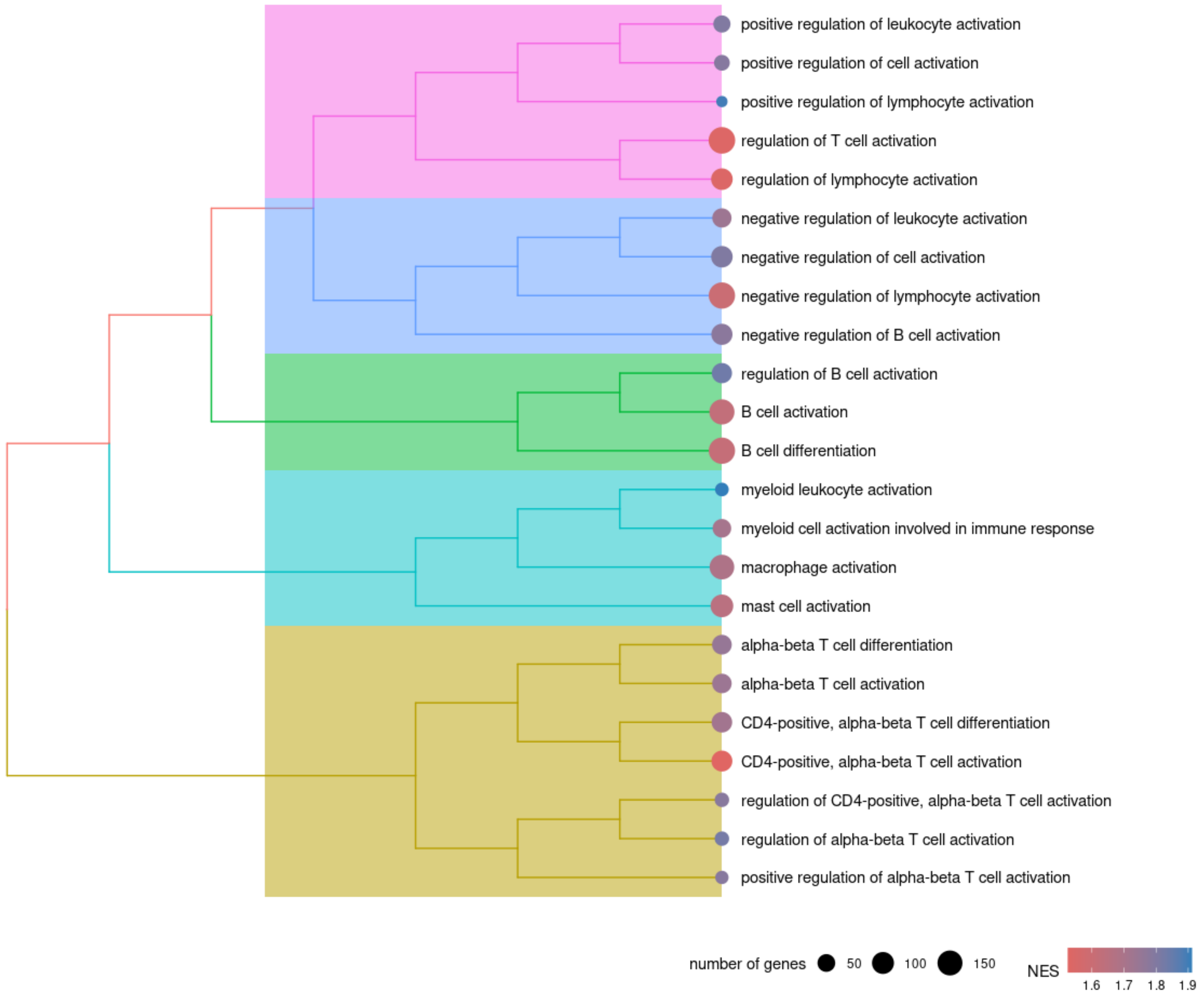

### MASH vs Control - Myeloid cells activity cluster

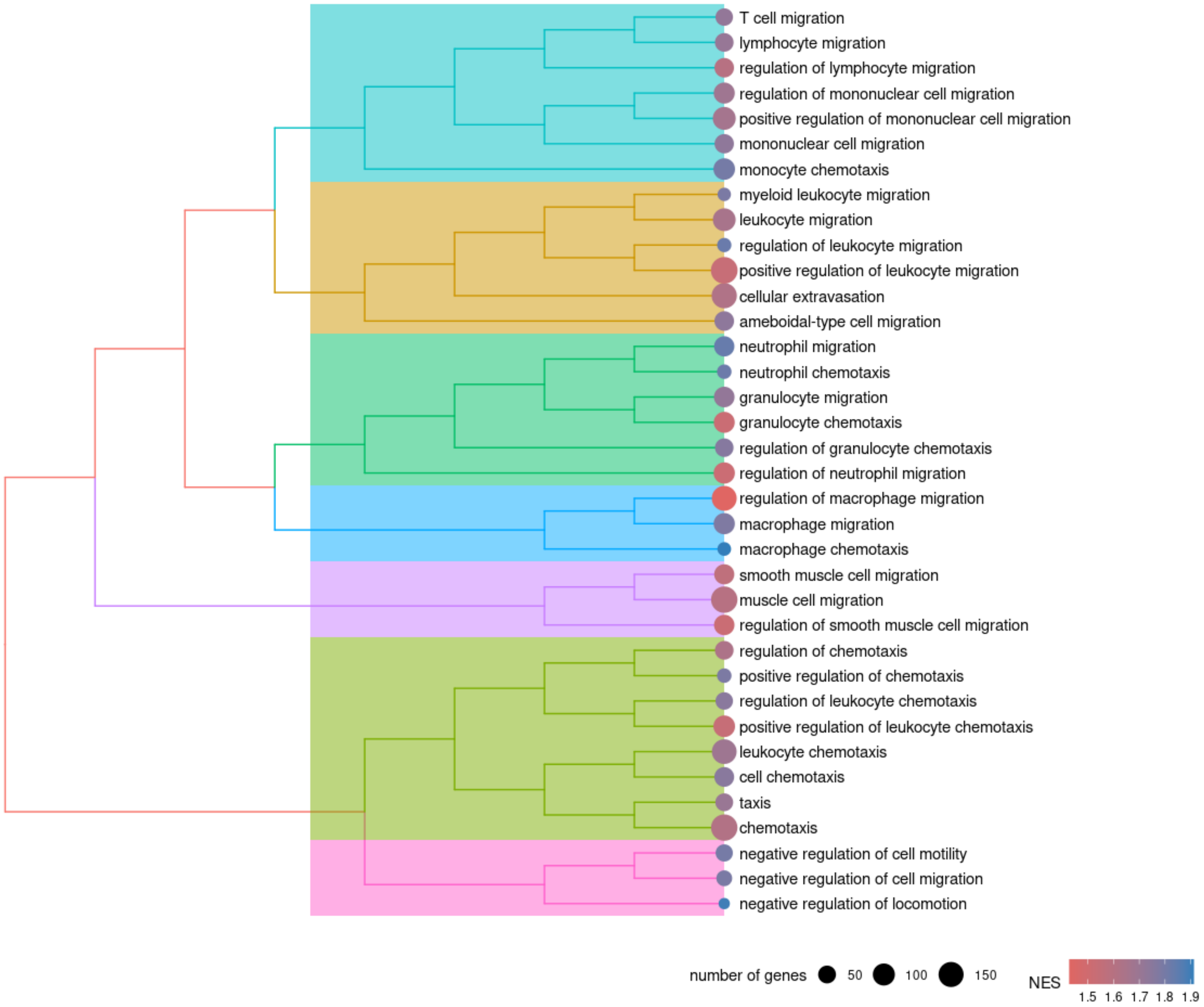

### MASH vs Control - Regulation of collagen metabolism cluster

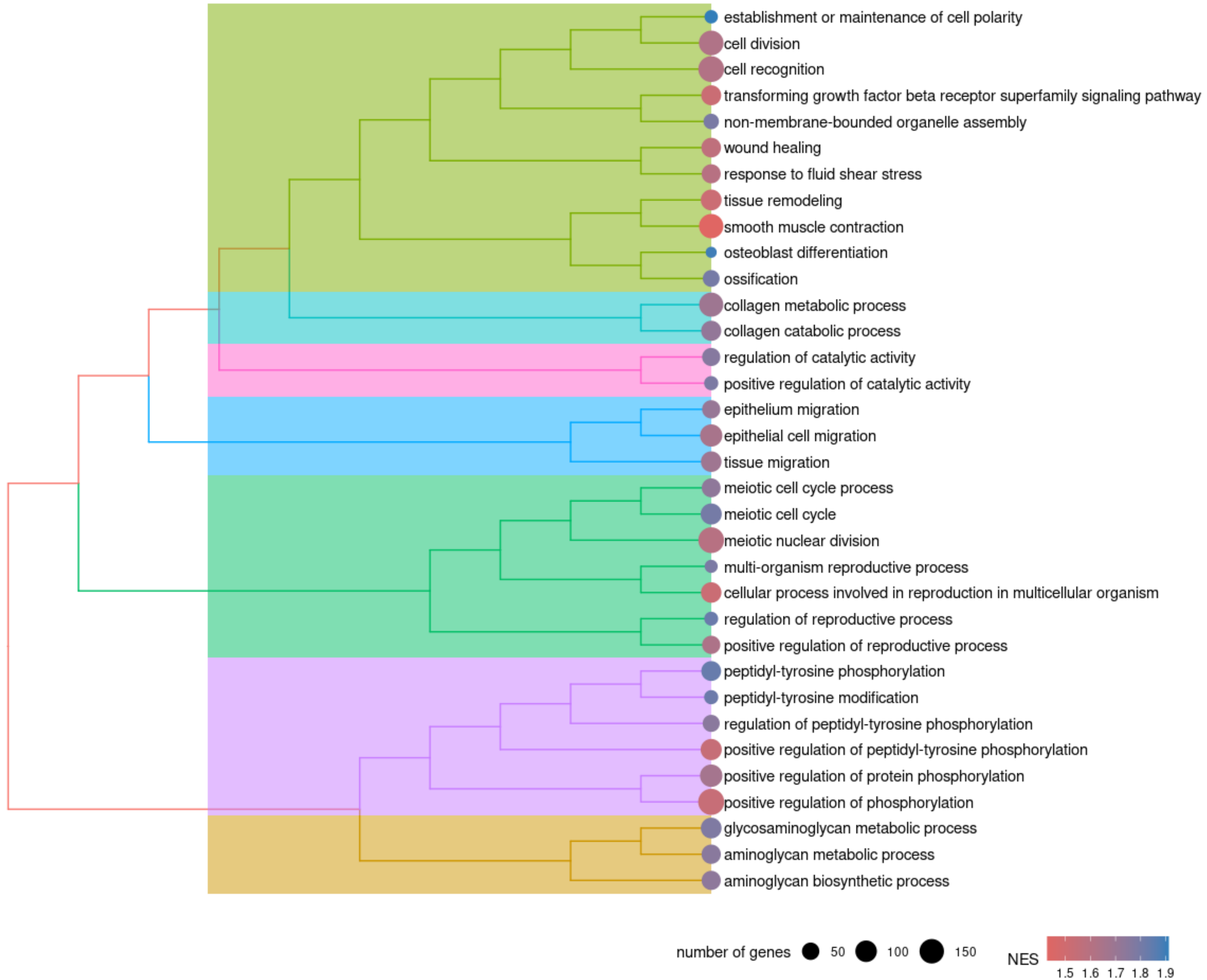

### MASH vs Control - Mesenchymal / stem cell development cluster

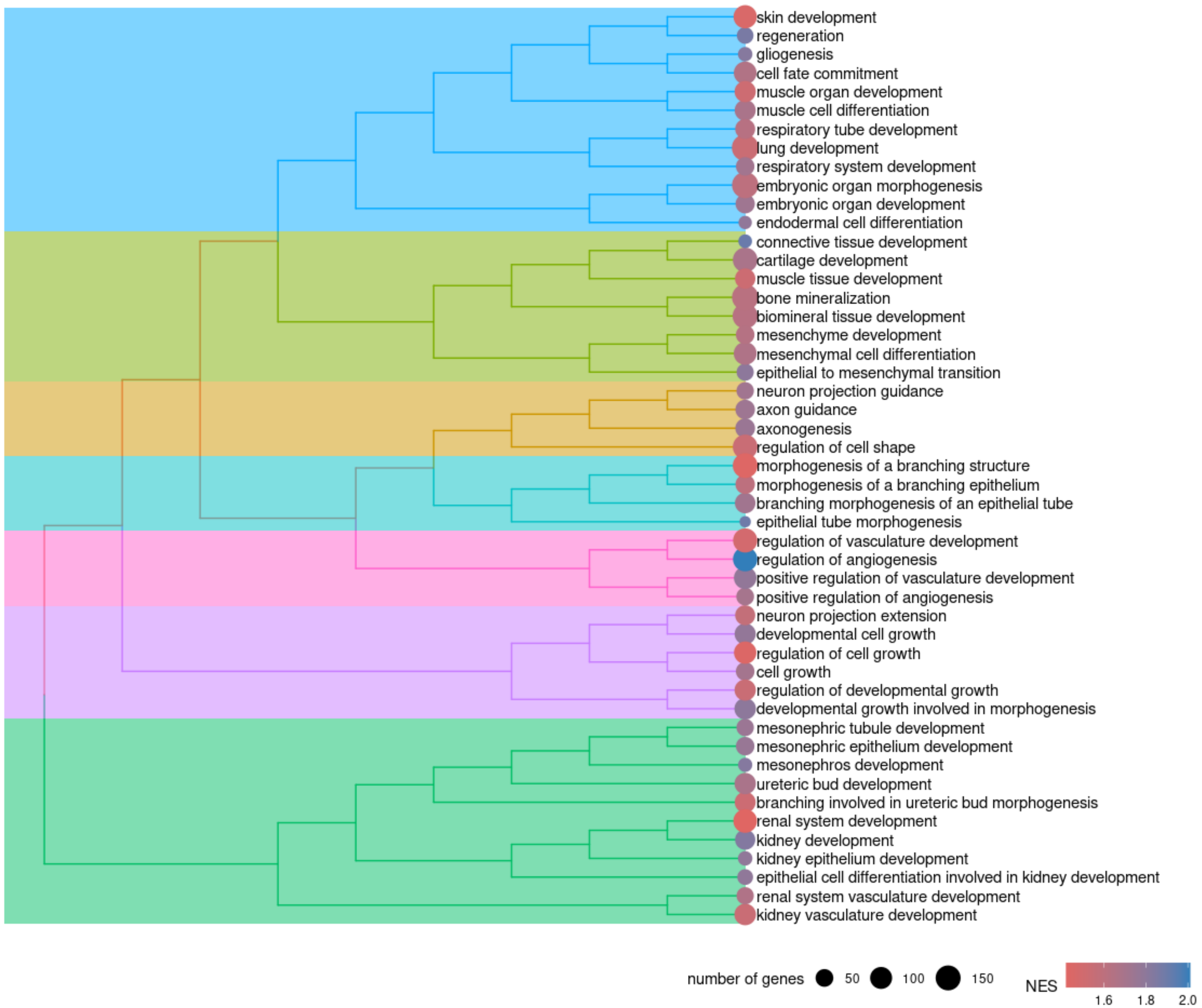

### MASH vs Control - Ion transport cluster

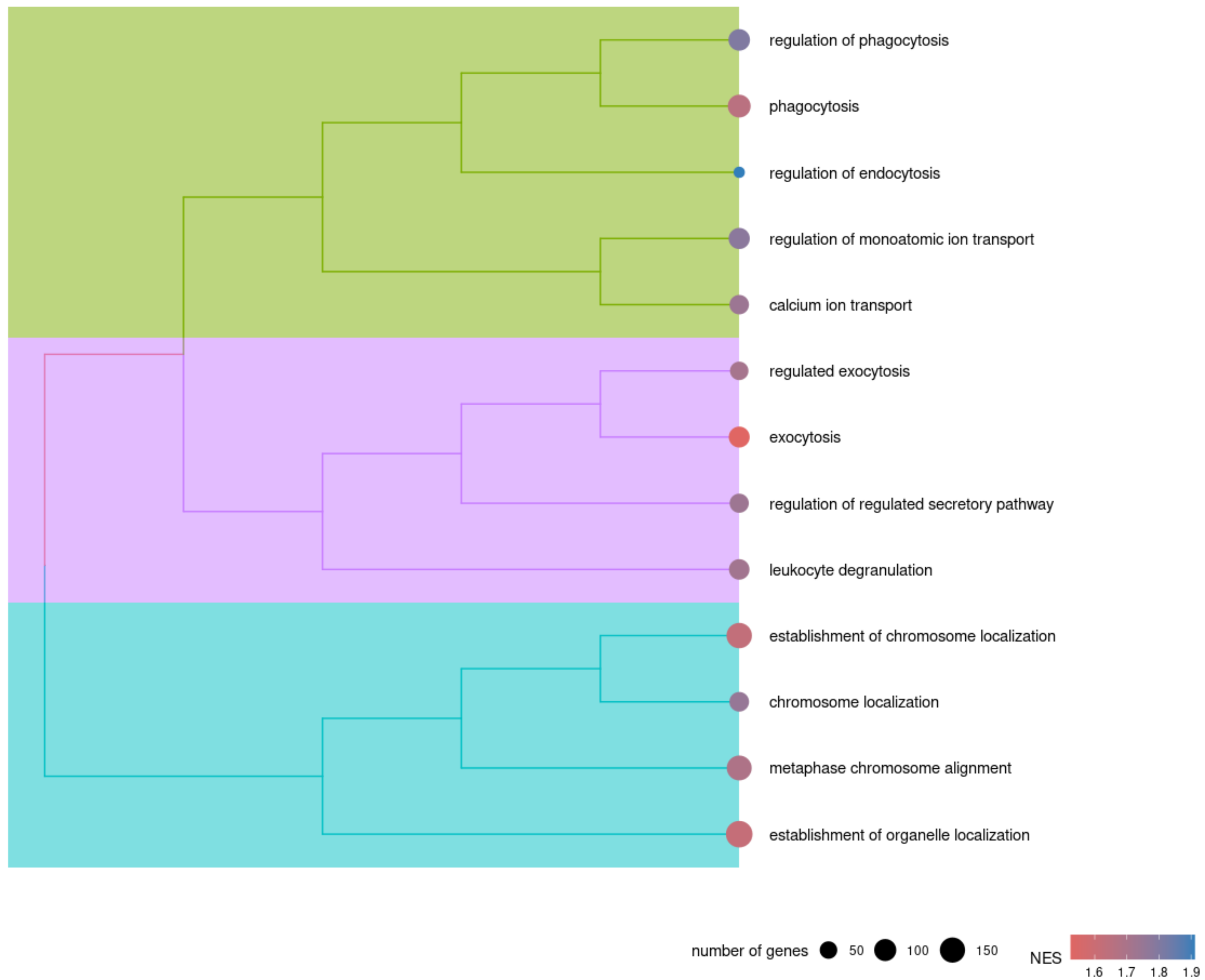

### MASH vs Control - Cell adhesion cluster

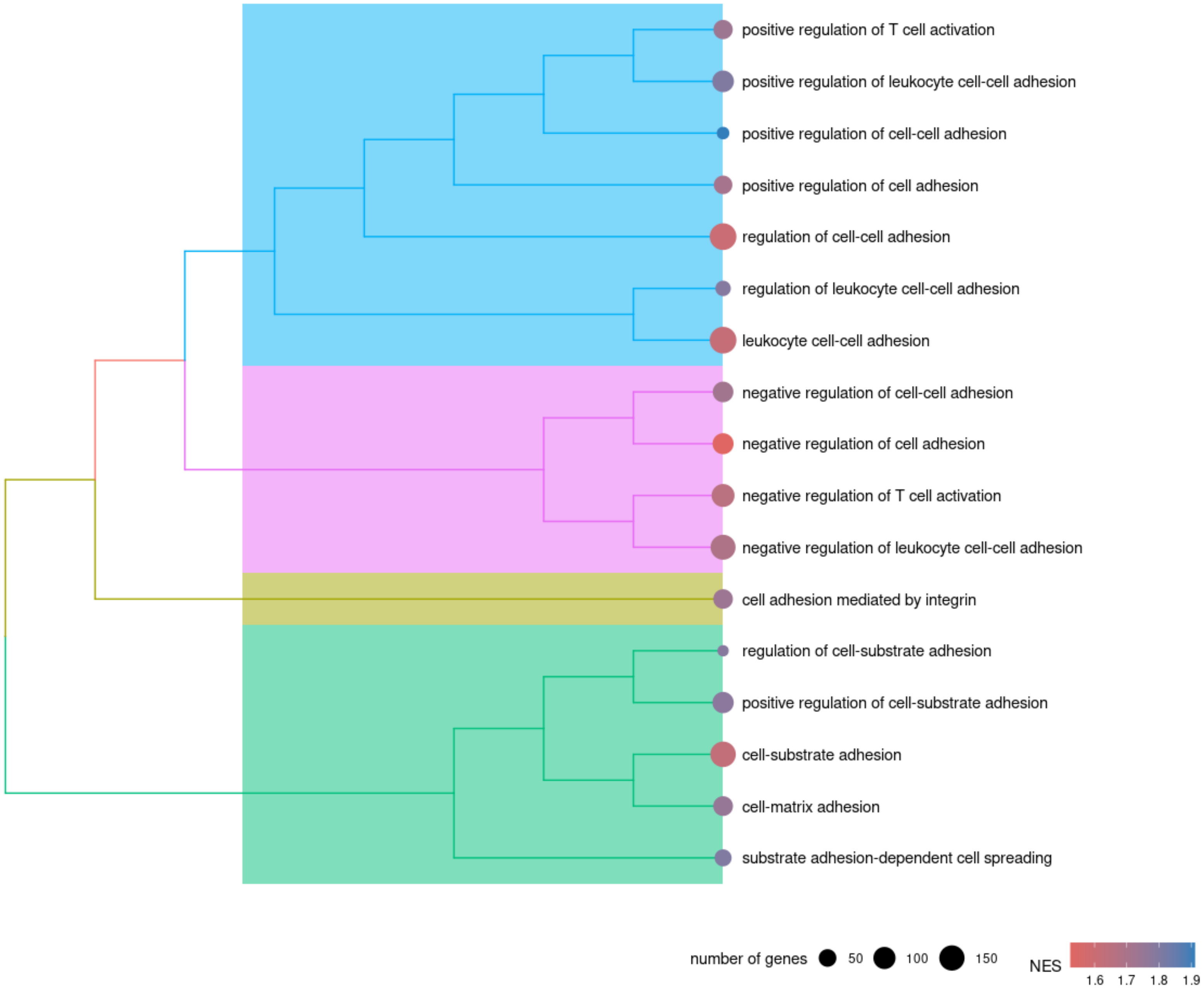

### MASH vs Control - Cell division process cluster

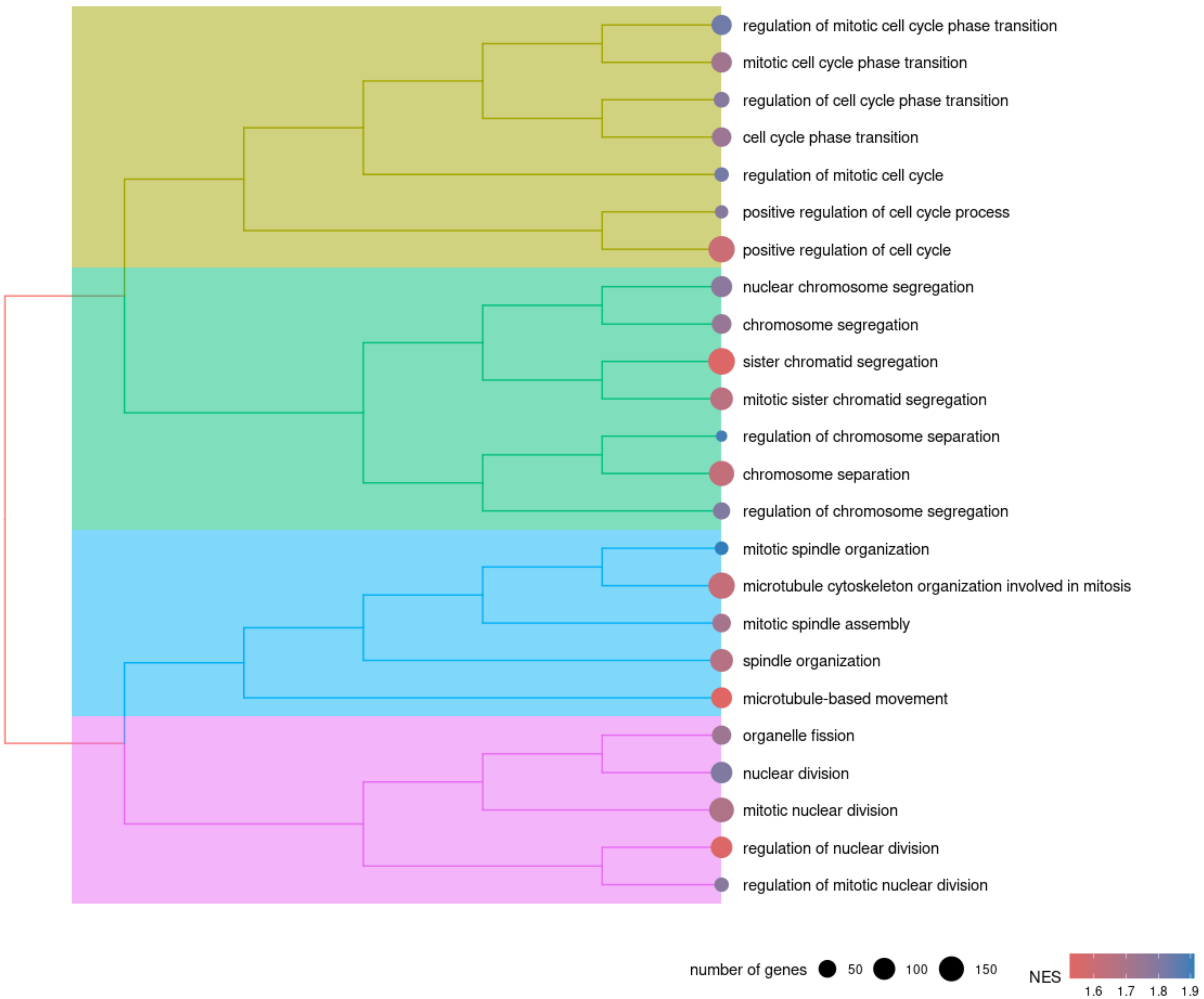

### MASH vs Control - TNF & Interleukin production cluster

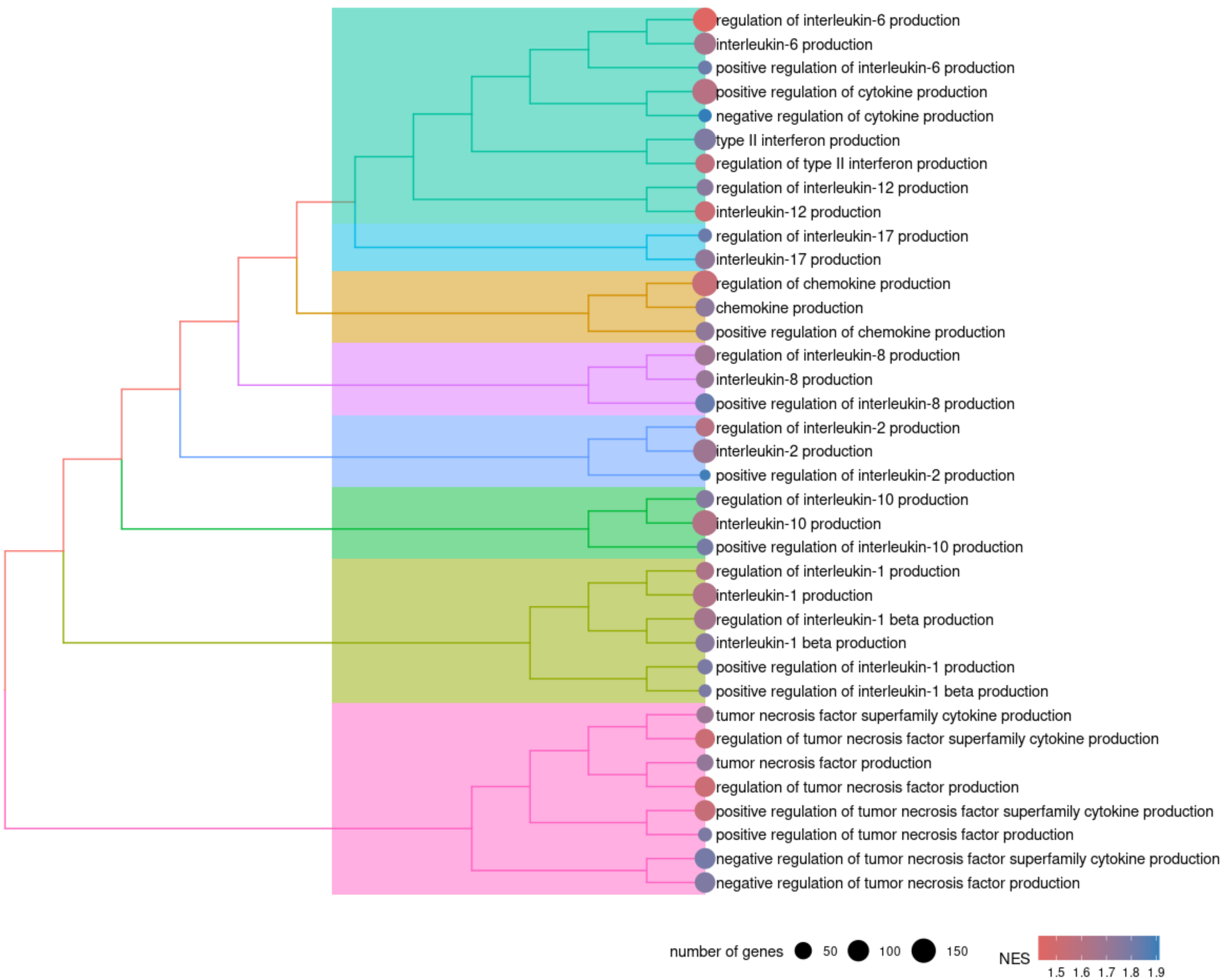

### MASH vs Control - Lymphocytes profileration cluster

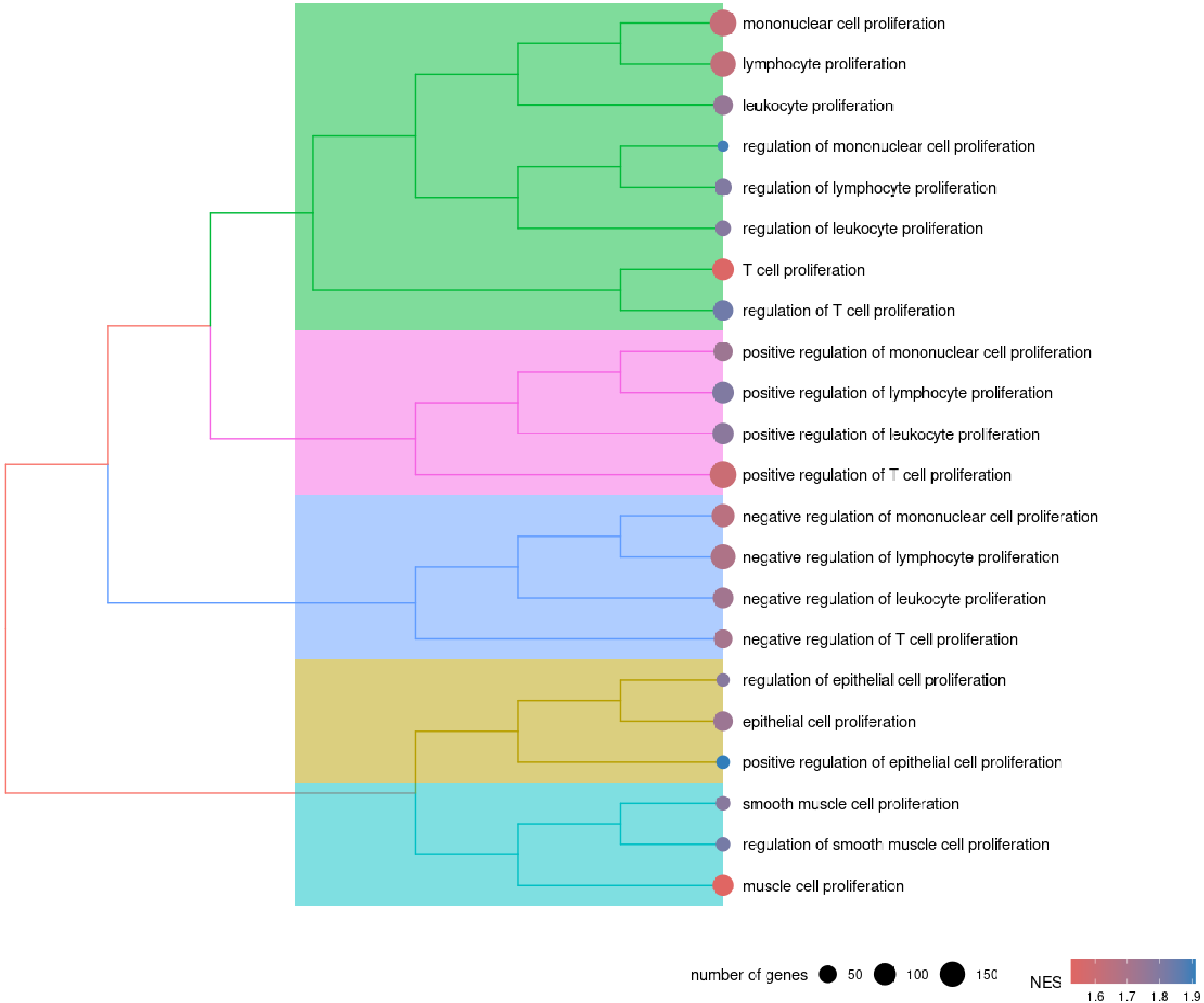

### MASH vs Control - Lymphocytes differentiation cluster

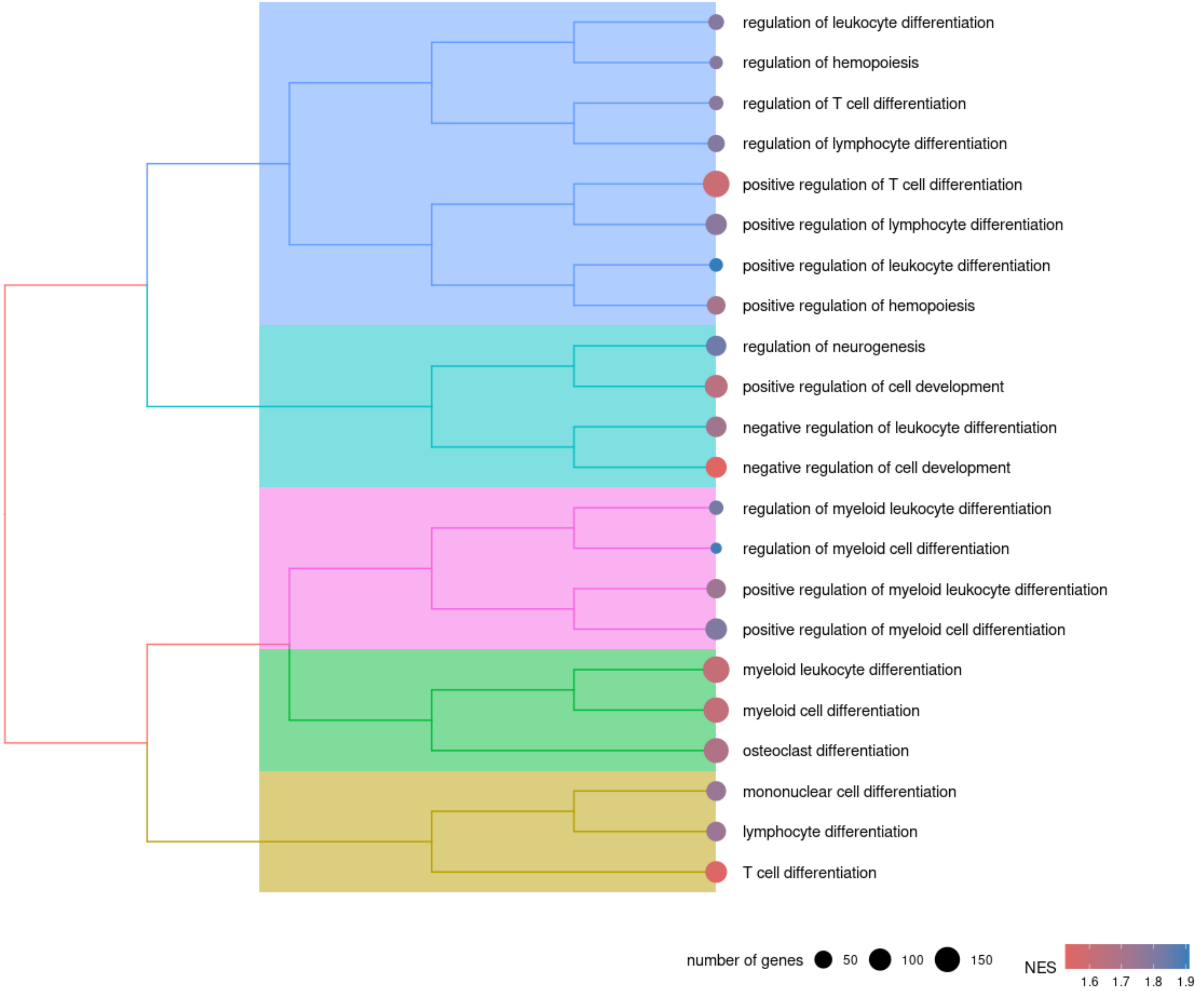

### MASH vs MASH + OATD-01 - Extracellular matrix organization cluster

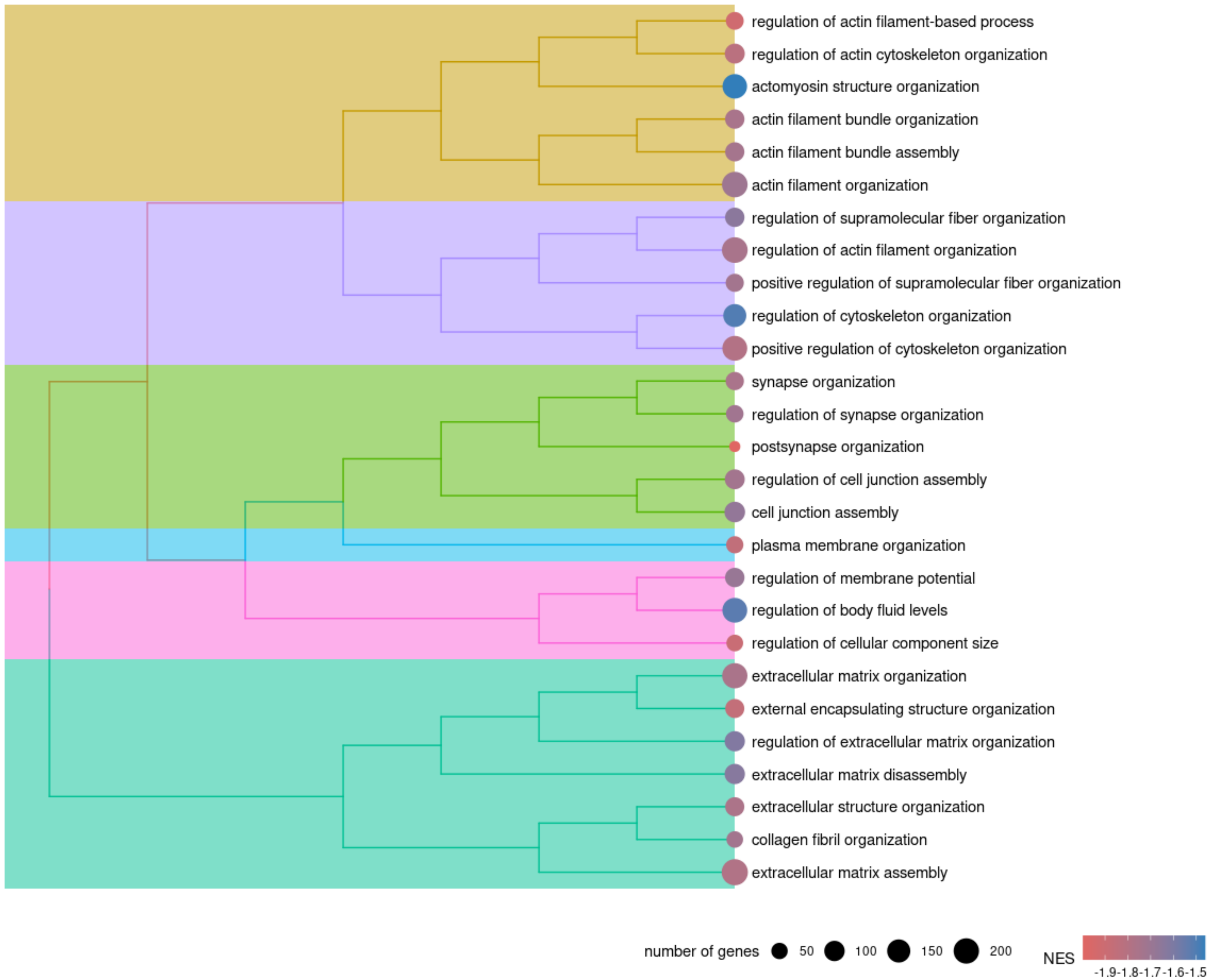

### MASH vs MASH + OATD-01 - Lymphocytes activation & differentiation cluster

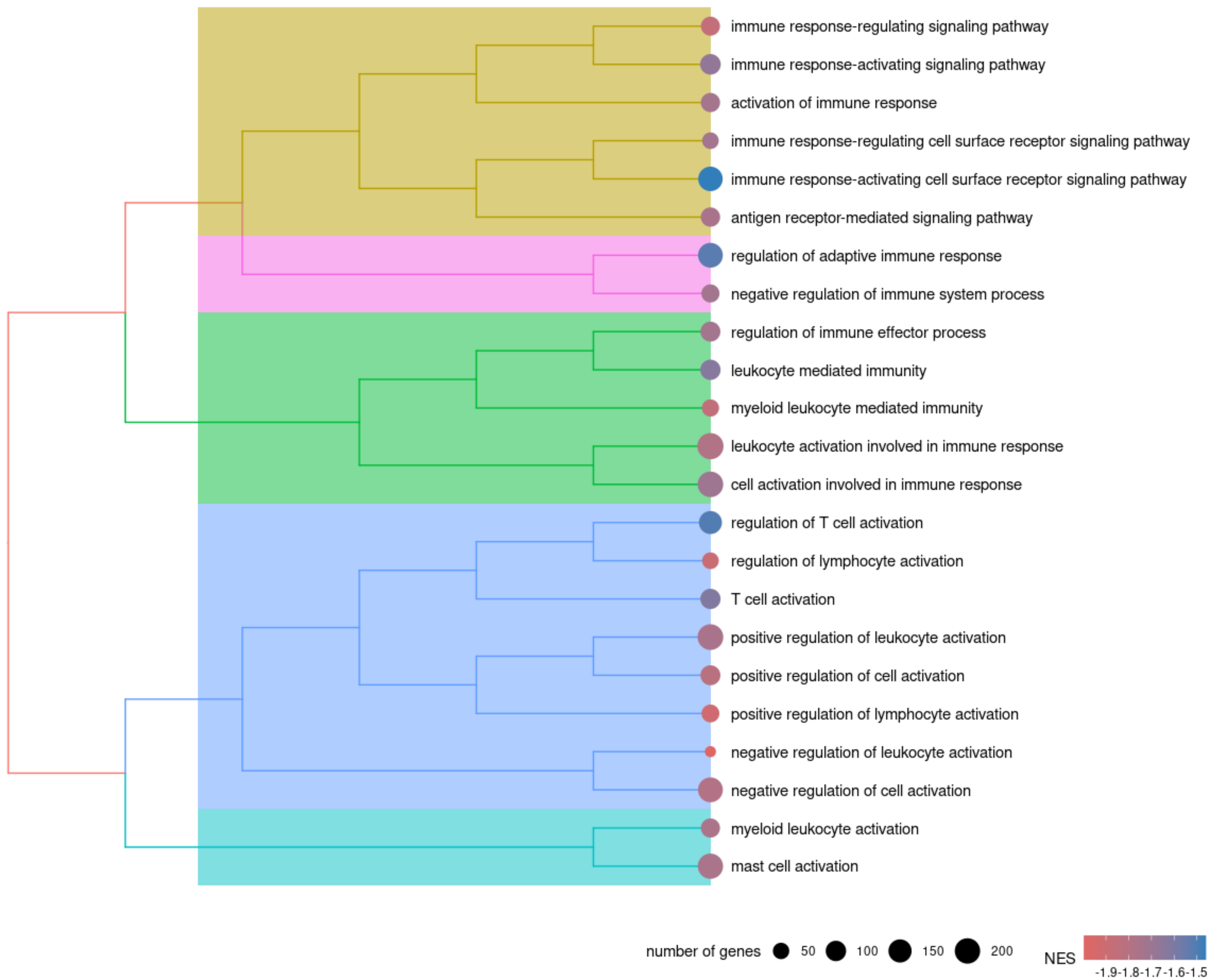

### MASH vs MASH + OATD-01 - Wnt signalling pathway cluster

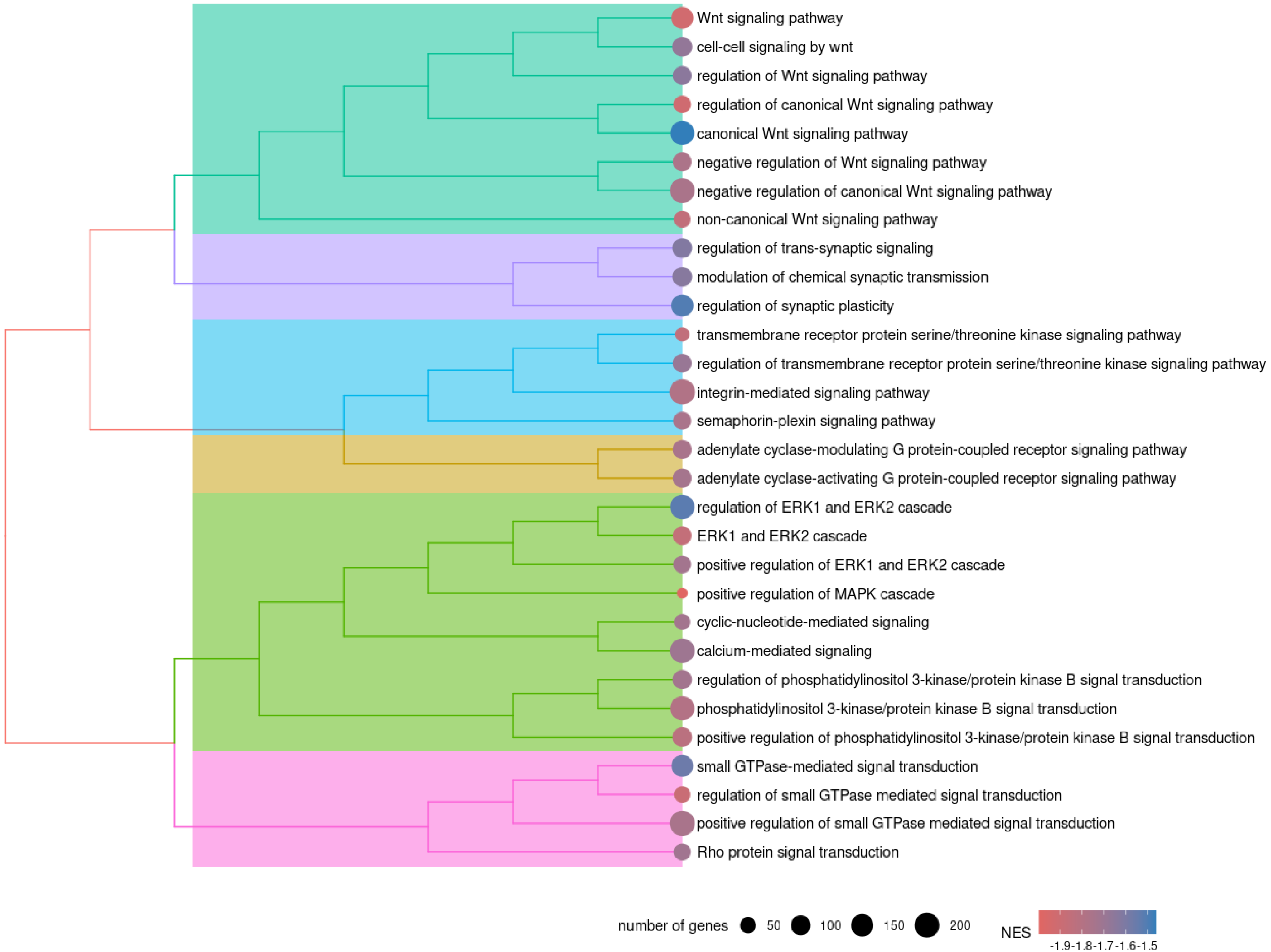

### MASH vs MASH + OATD-01 - Myeloid cells activity cluster

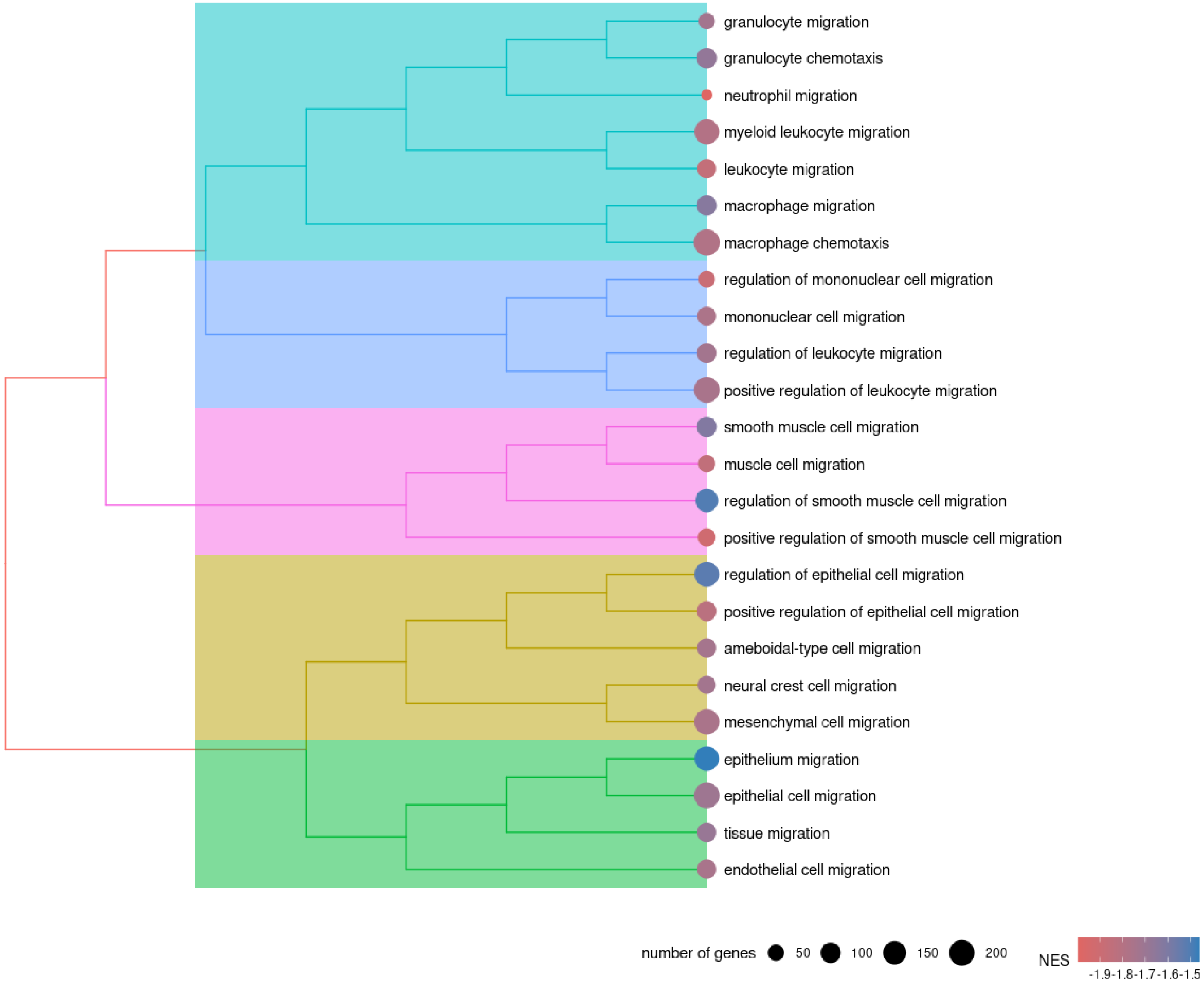

### MASH vs MASH + OATD-01 - Mesenchymal / stem cell development cluster

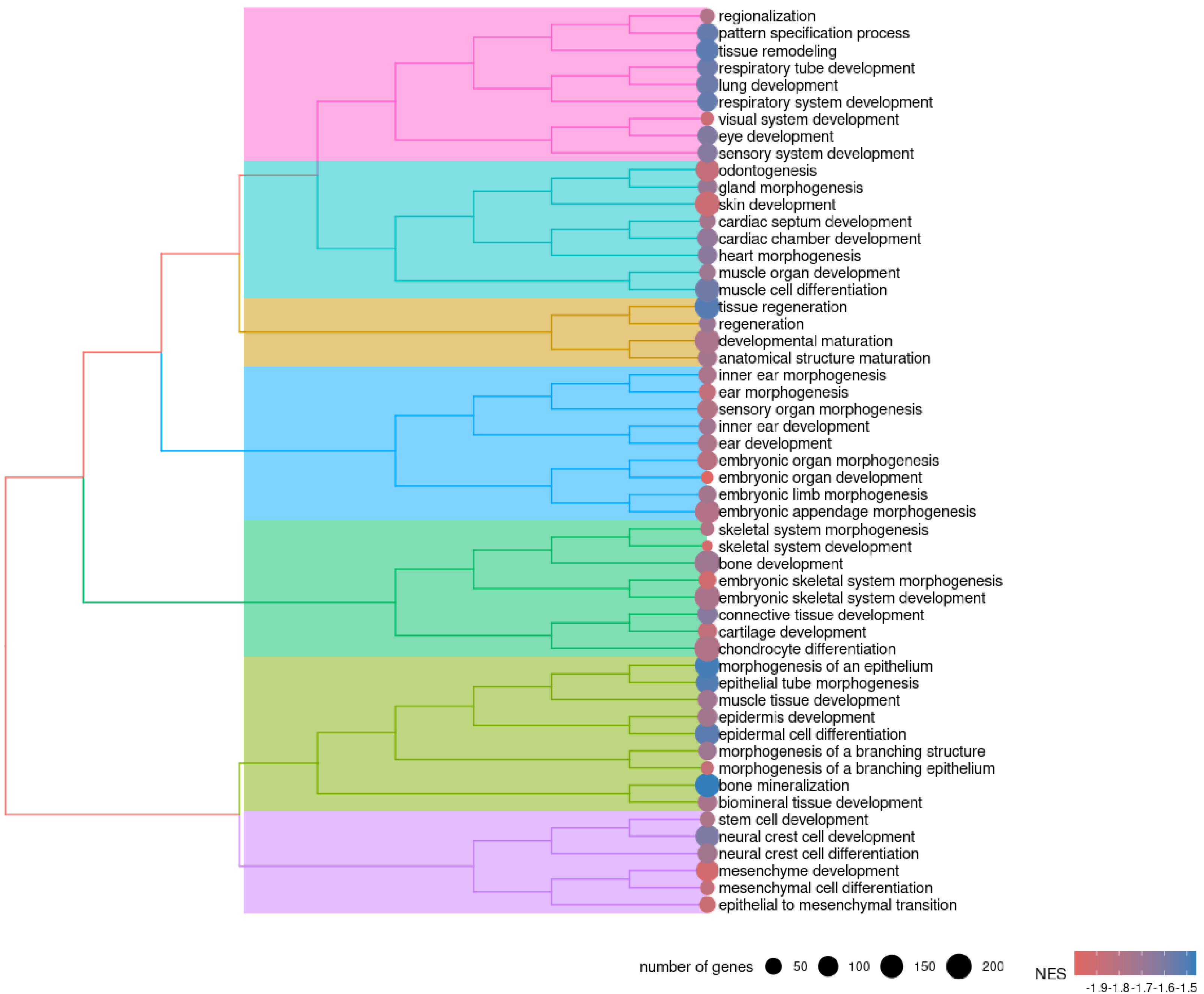

### MASH vs MASH + OATD-01 - Regulation of neurogenesis cluster

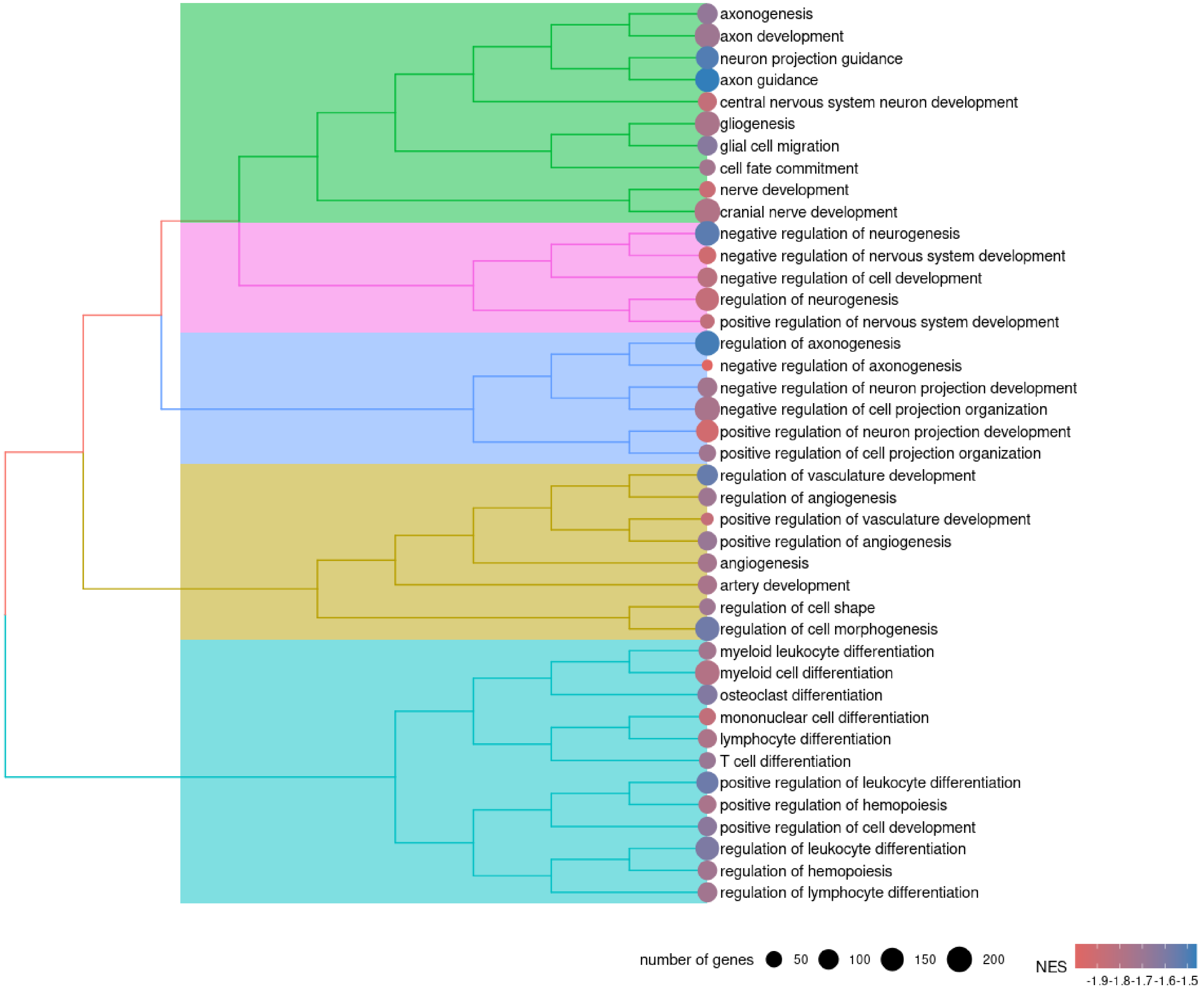

### MASH vs MASH + OATD-01 - Regulation of cell growth cluster

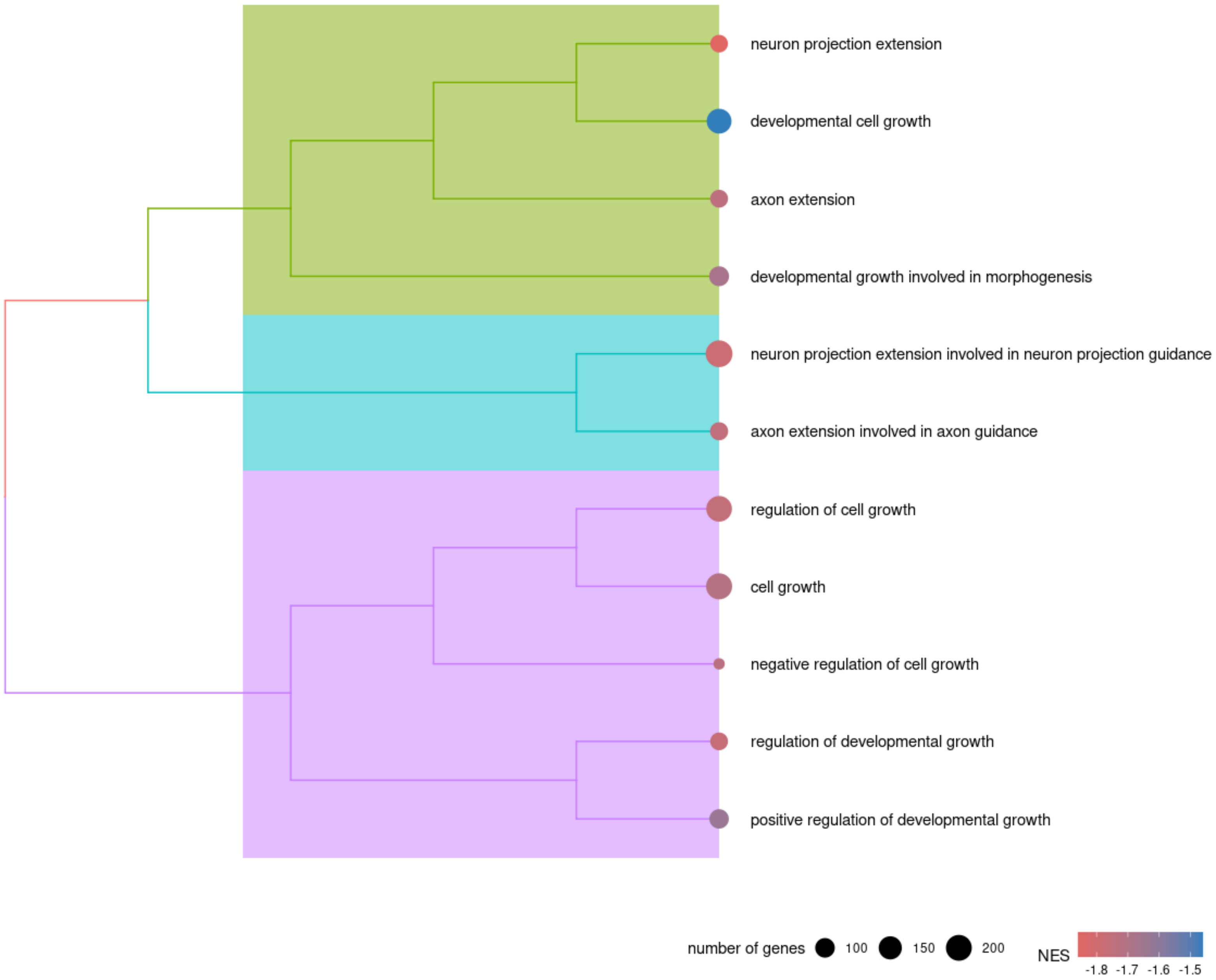

### MASH vs MASH + OATD-01 - Blood circulation cluster

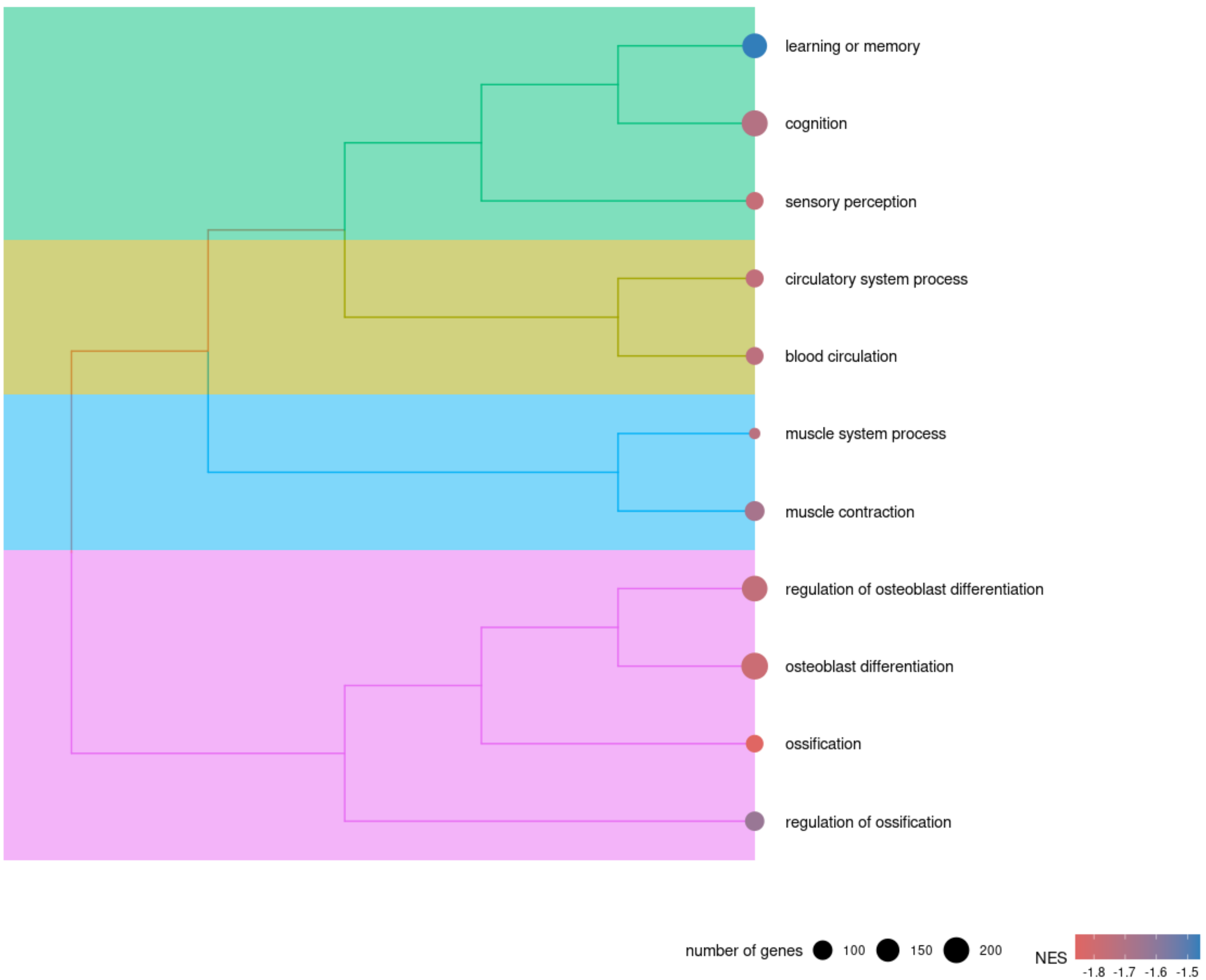

### MASH vs MASH + OATD-01 - Regulation of collagen metabolism cluster

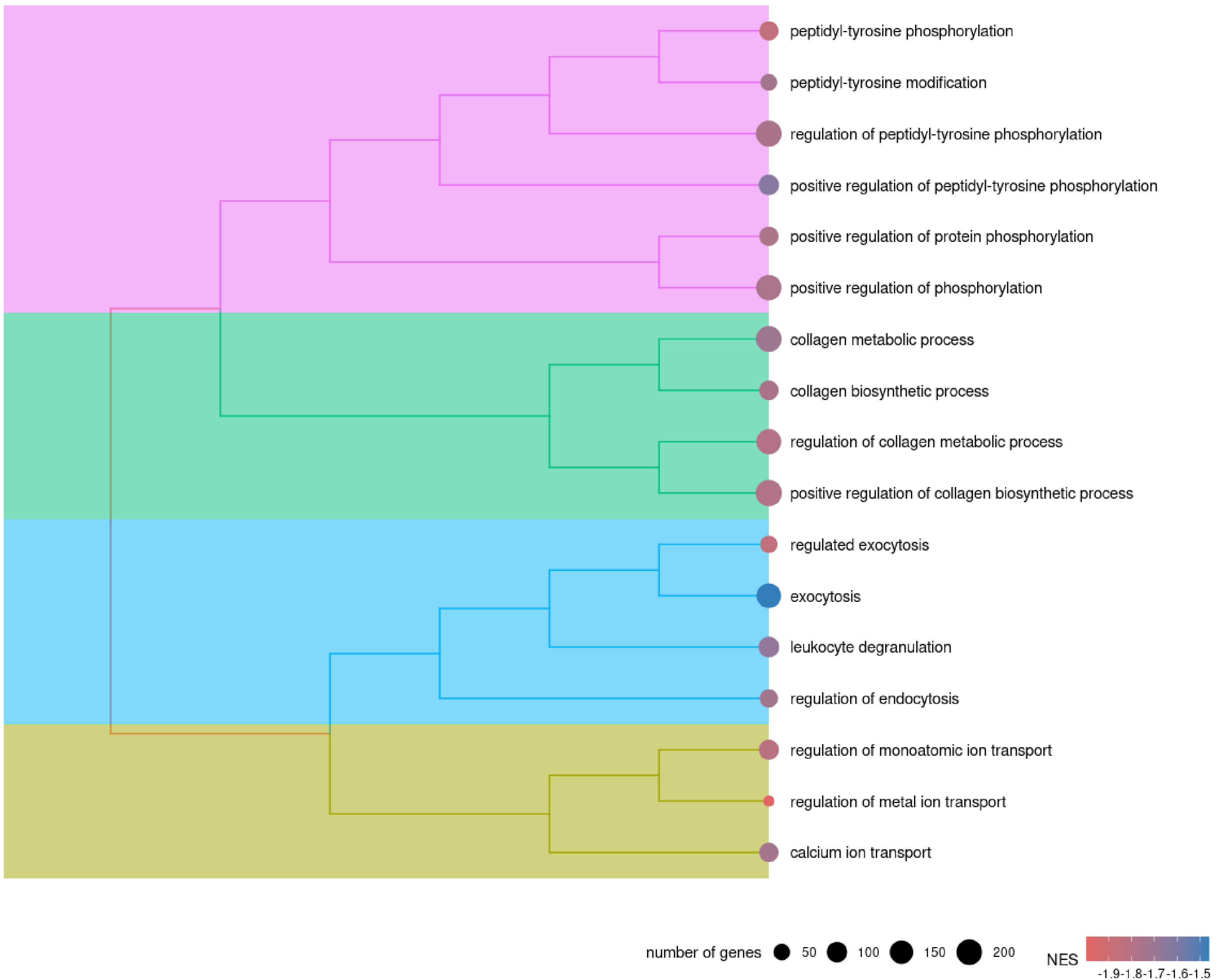

### MASH vs MASH + OATD-01 - Cell adhesion cluster

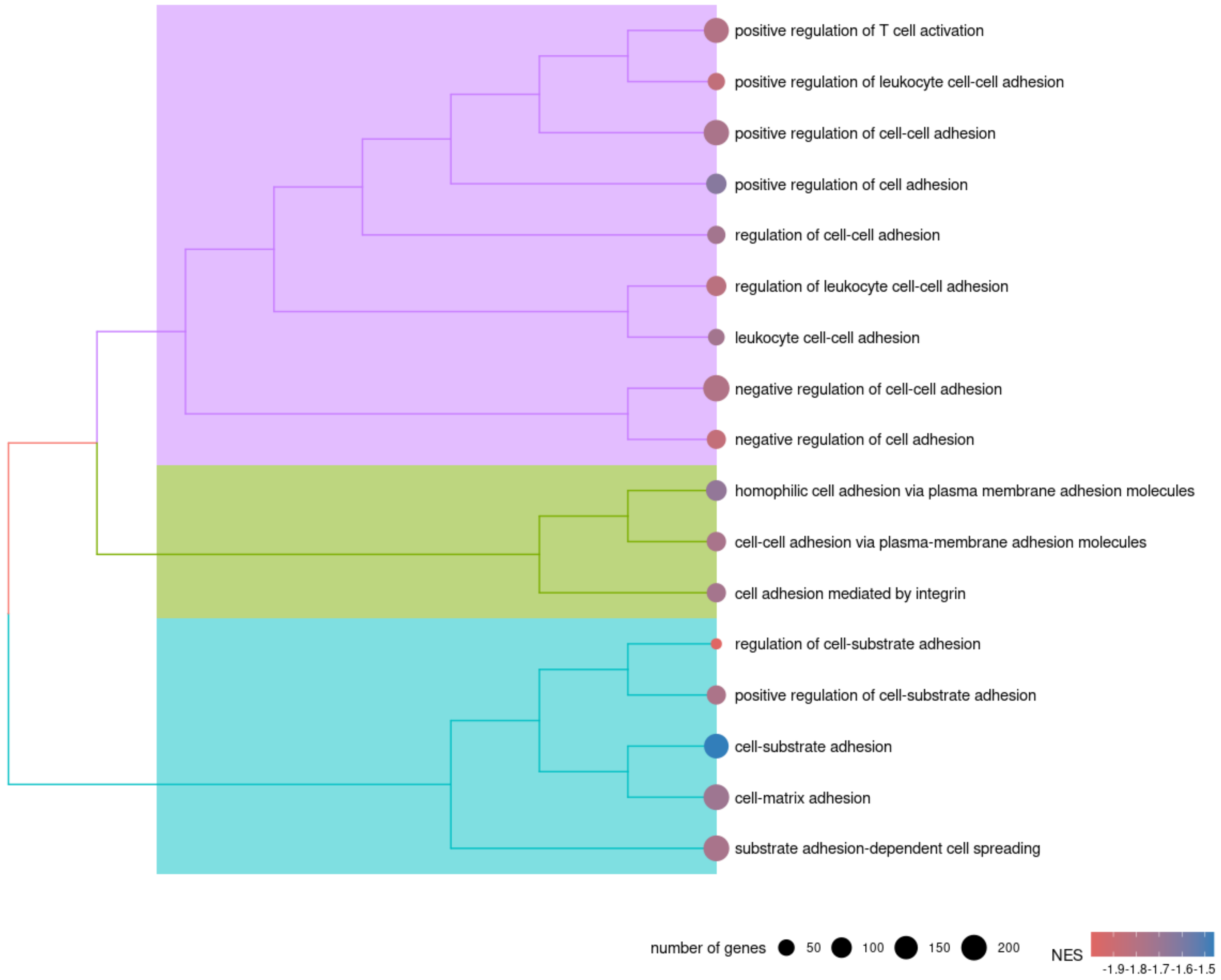

### MASH vs MASH + OATD-01 - Chemotaxis cluster

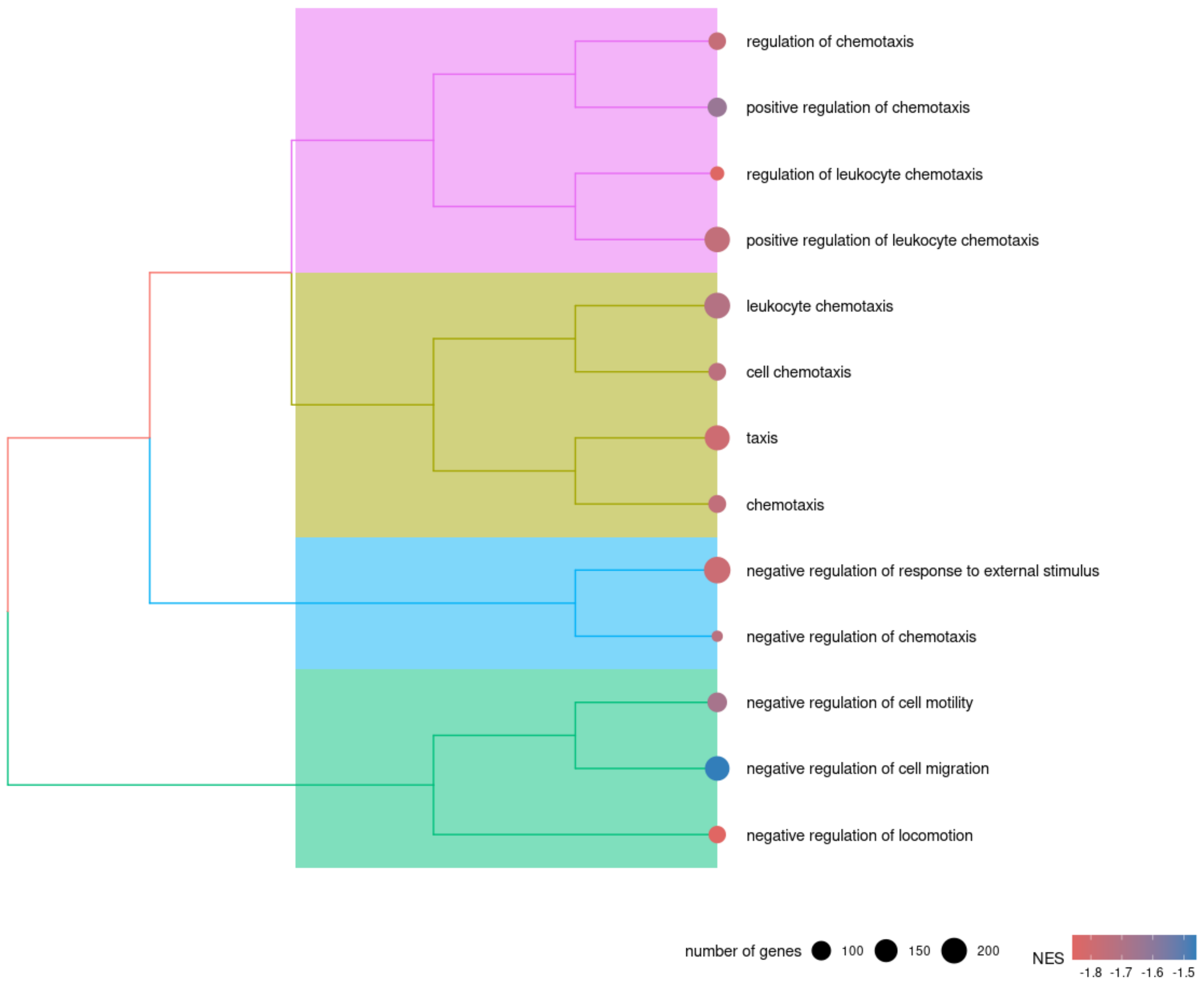

### MASH vs MASH + OATD-01 - Epithelial cells development cluster

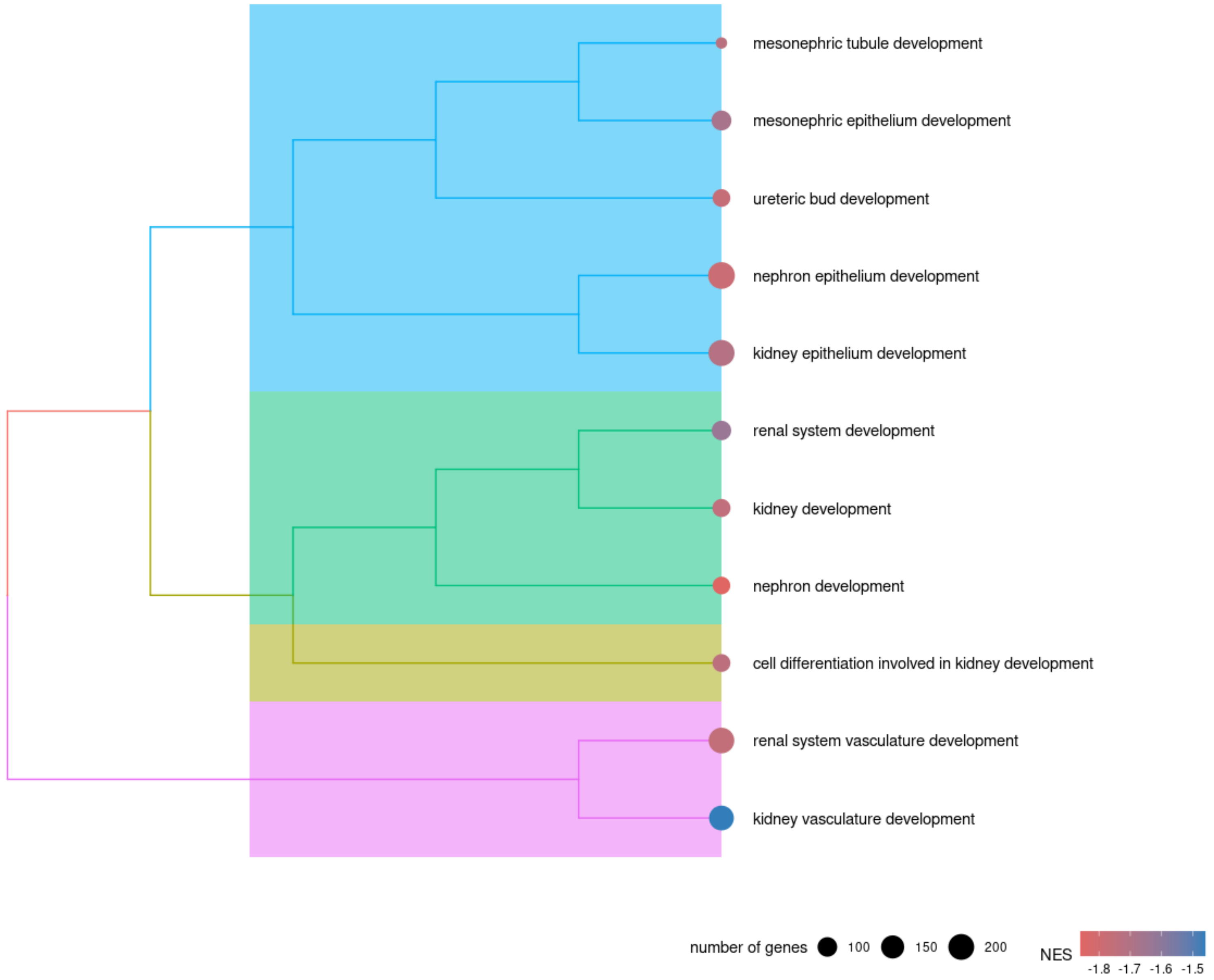

### MASH vs MASH + OATD-01 - Cellular response cluster

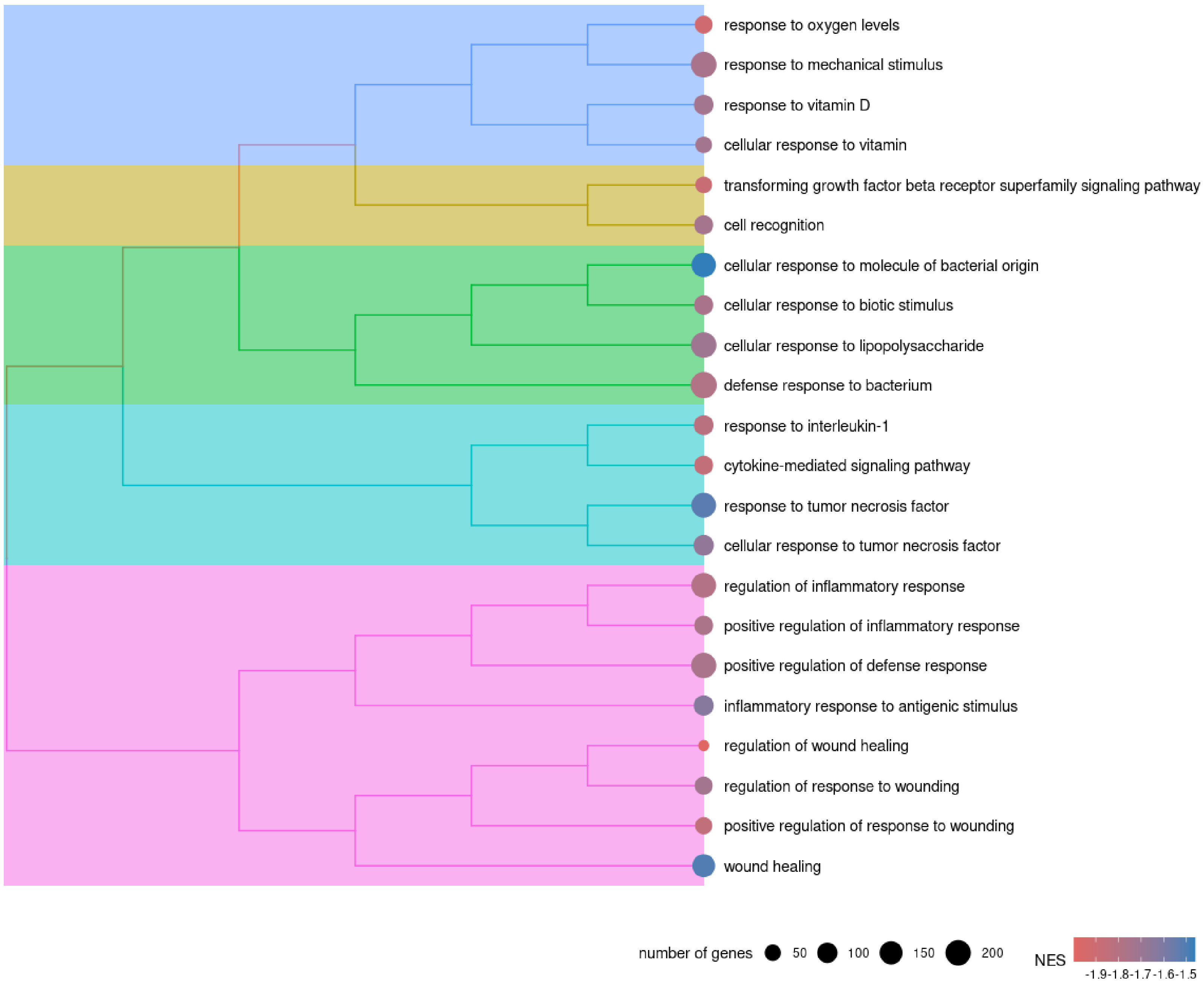

### MASH vs MASH + OATD-01 - Cellular response to growth factor cluster

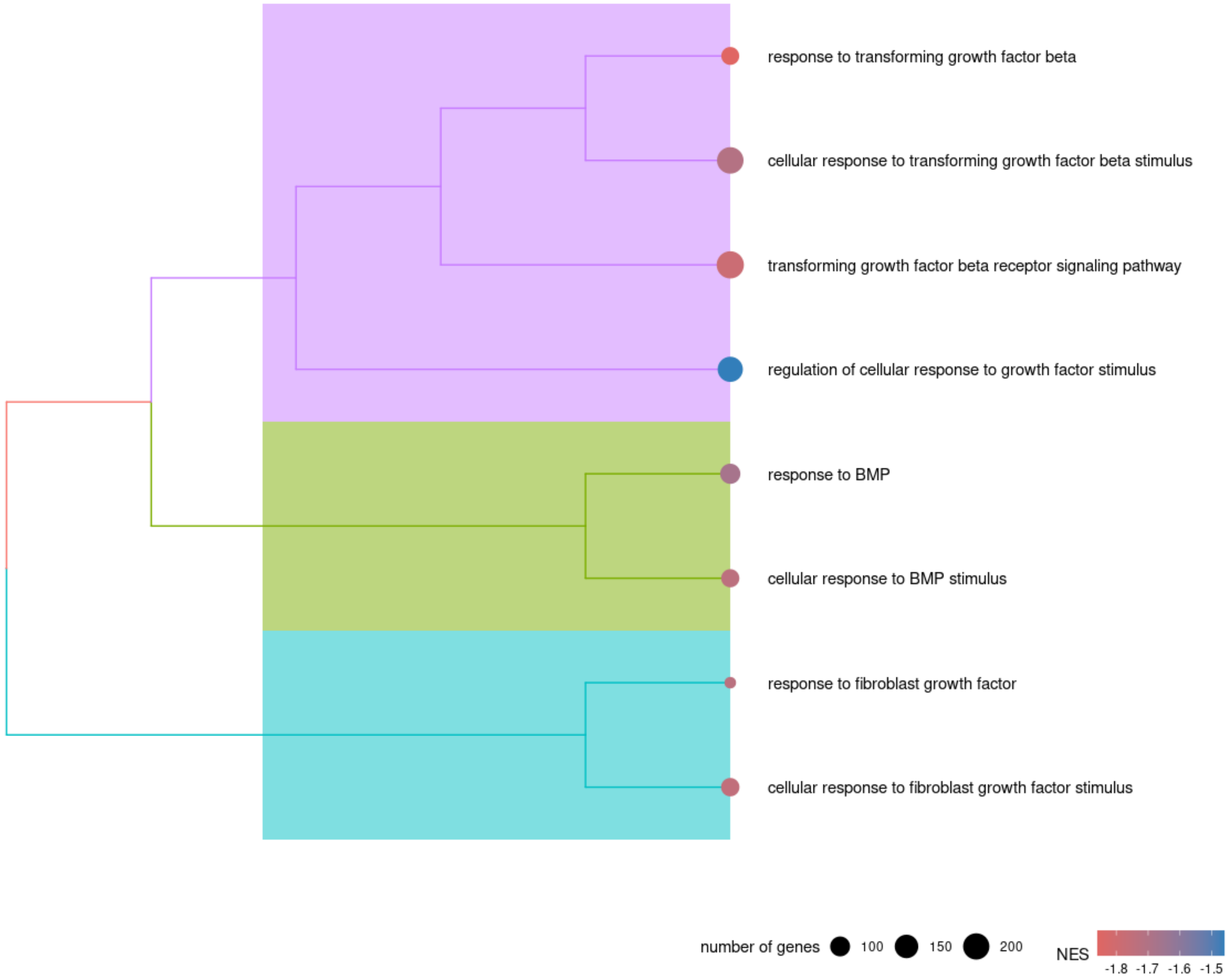

### MASH vs MASH + OATD-01 - Lymphocytes profileration cluster

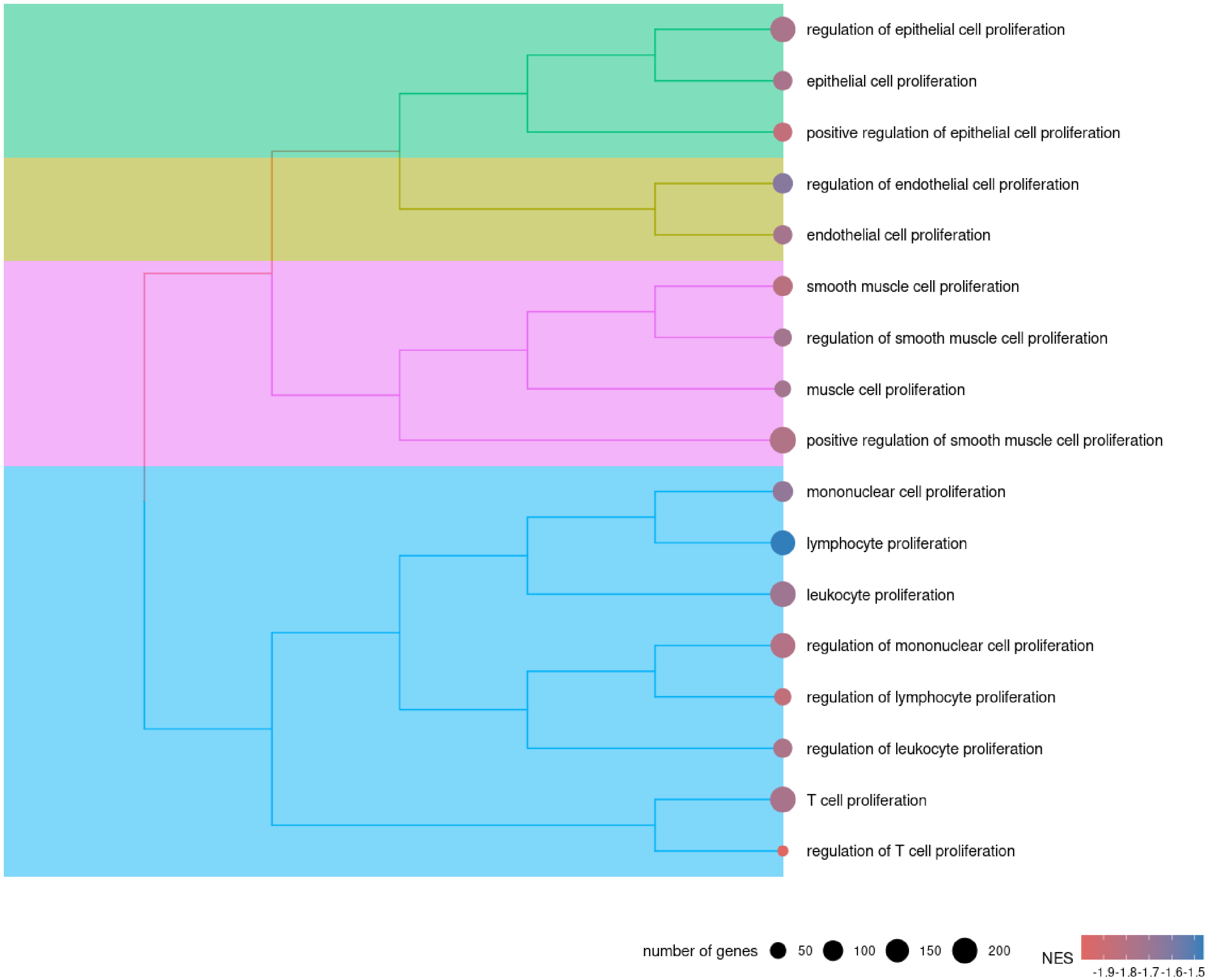

### MASH vs MASH + OATD-01 - TNF & Interleukin production cluster

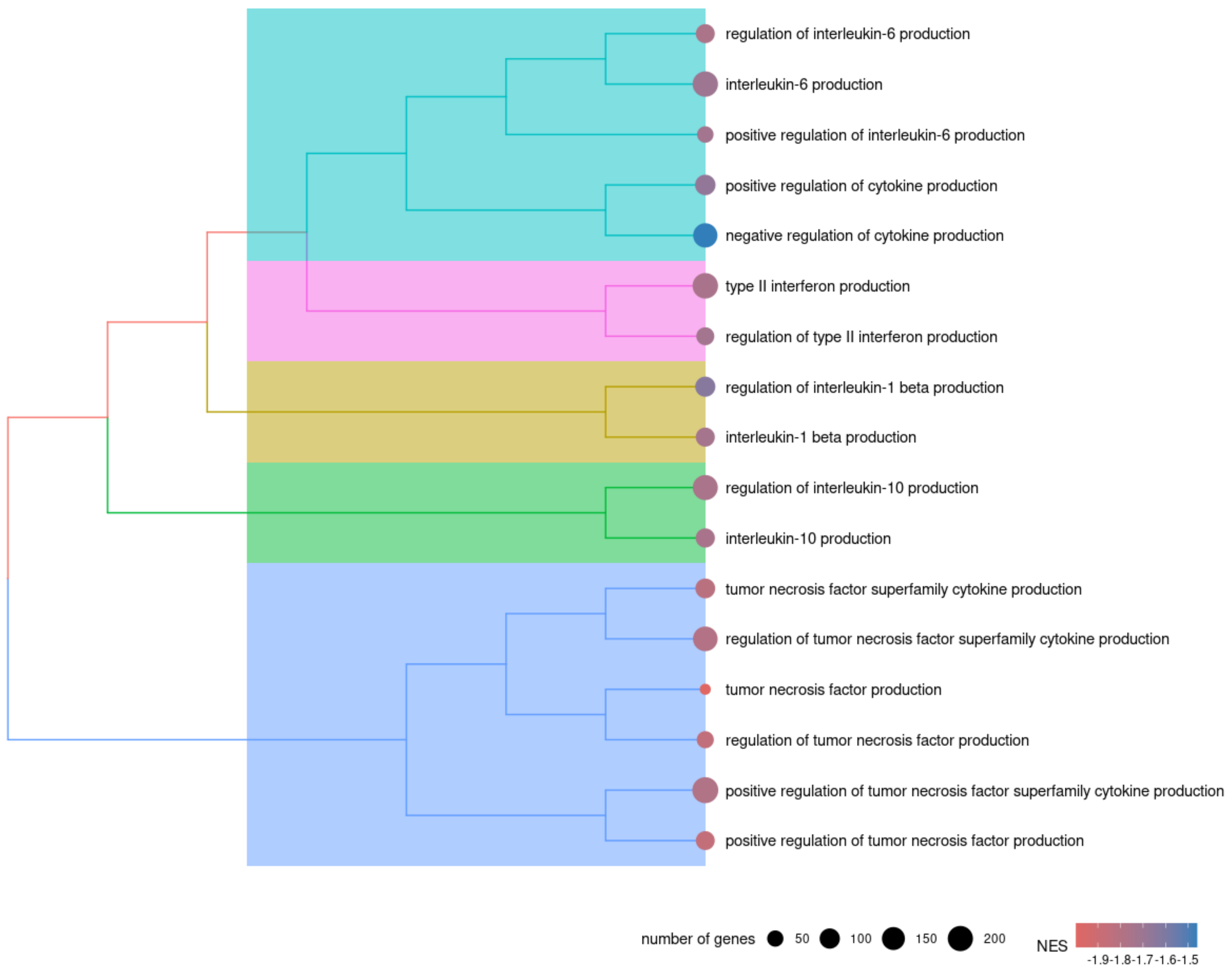
